# Local extraction of predatory fish may initiate behaviourally mediated trophic cascades in coral reefs in the Andaman Islands

**DOI:** 10.1101/2025.11.17.688842

**Authors:** Shawn Dsouza, Akshta Joshi, Bharat Ahuja, Samar Ahmad, Kartik Shanker

## Abstract

Humans disproportionately target predators due to their higher economic and social value. This has led to widespread declines of predators in ecosystems worldwide leading to far-reaching effects on ecosystem processes. We investigate how the loss of predators from nearshore ecosystems due to fishing affects individual prey behaviour using a gradient of protection as a natural experiment. We collected data on piscivorous predator assemblages, reef habitats and prey behaviour within and outside two marine protected areas (MPAs) in the South Andaman Islands, India. As expected, predator abundance was significantly greater within MPAs. Similarly, we found greater coral cover and reduced algal cover within MPAs. In response, prey species, which included herbivorous and invertivorous fish, exhibited reduced movement and heightened vigilance inside MPAs. Additionally, prey individuals foraged at higher rates within MPAs. Habitat complexity and resource availability had complex effects on prey behaviours that varied across guilds. Our findings suggest that fishing directly alters the seascape of fear through the removal of predators. Fishing may also shape prey behaviour and ecological interactions indirectly through habitat degradation. Overall, this study highlights the need to study both direct and indirect consequences of human interactions with wild animals to understand their combined impacts on ecosystems.

## Introduction

Among the myriad effects of human activities in the Anthropocene, the loss of top predators due to overexploitation and conflict has led to some of the largest impacts on ecosystems globally (1,2). Predators regulate key ecosystem functions by exerting top–down control on prey, and the effects of predator removal can propagate through food chains as trophic cascades (3). Although humans have been hunter–gatherers for most of their evolutionary history and often fill the niche of predators in many ecosystems, their impact has increased dramatically with modern tools and technology (4). In fact, human predators kill more prey than all predators combined across terrestrial and marine ecosystems (5).

Many prey species demonstrate explicit behavioural anti-predator strategies to mitigate the risk of depredation (6,7). However, the loss of predators due to human activities such as fishing may reduce the local intensity of predation risk experienced by prey. Thus, prey in areas with higher anthropogenic pressure on predators may be able to allocate their time to different behaviours than prey individuals in areas with low anthropogenic pressure. For example, fish outside marine protected areas may be able to spend more time foraging than fish within protected areas as there may be greater predator abundance within protected areas (8). In addition, reduced predator density can lead to a homogenisation of the spatial distribution of predation risk, also known as the “landscape of fear” (sensu 9; 6,7). Furthermore, changes in the spatial distribution of predation risk perceived by herbivorous fish species can lead to altered distribution of sea grass in patchy coral reefs (10). Such a change in prey behaviour due to altered risk that changes the abundance, distribution, and composition of lower trophic levels, is known as a behaviourally mediated trophic cascade (11).

These non-consumptive effects of predators are highly context dependent. Habitat characteristics such as refuge availability reduce predation risk for prey, thereby changing the spatial distribution of risk at the local scale (13). Bauman et al. (14) demonstrated that seagrass cover in a coral reef ecosystem reduced the response of prey to predator models by providing additional refuge. Prey may also respond differently based on predator hunting modes. For example, the landscape of risk from a sit-and-wait type or ambush predators like groupers (Epinephelidae) is different from a coursing predators like barracudas (Sphyraenidae) (15). Prey behaviours such as grouping may also reduce the overall risk experienced by prey individuals (16).

Fisheries tend to disproportionately target top predators and larger fish due to their higher economic value. (13,14,16). Historically, overfishing has led to the loss of top predators in many nearshore marine ecosystems. The reduced predation pressure has had widespread ecological consequences, including mesopredator release and in some cases, the collapse of fisheries (2). As predators have been fished out, fisheries have had to expand in spatial scale and effort while also targeting lower trophic levels (17), with downstream impacts across habitats. The loss of predators from coral reefs can also change the distribution of seagrass meadows which are vital for blue carbon sequestration (13–15, 17,18).

Here, we examine how fishing alters the seascape of risk in coral reefs by comparing predator assemblages and prey behaviour inside and outside two no-take protected areas in the Andaman Islands, India. We expected predator abundance to be lower outside protected areas due to their removal by fisheries. Similarly, we anticipated lower coral cover and reduced habitat complexity outside protected areas as a result of damage caused by fishing gear and boat anchors. These changes in predator distribution and habitat structure were expected to shift the spatial pattern of predation risk, leading fish to invest less time in anti-predator vigilance and more time in foraging outside protected areas, consistent with the risk-allocation hypothesis (20) (Figure 1).

**Figure 1:**
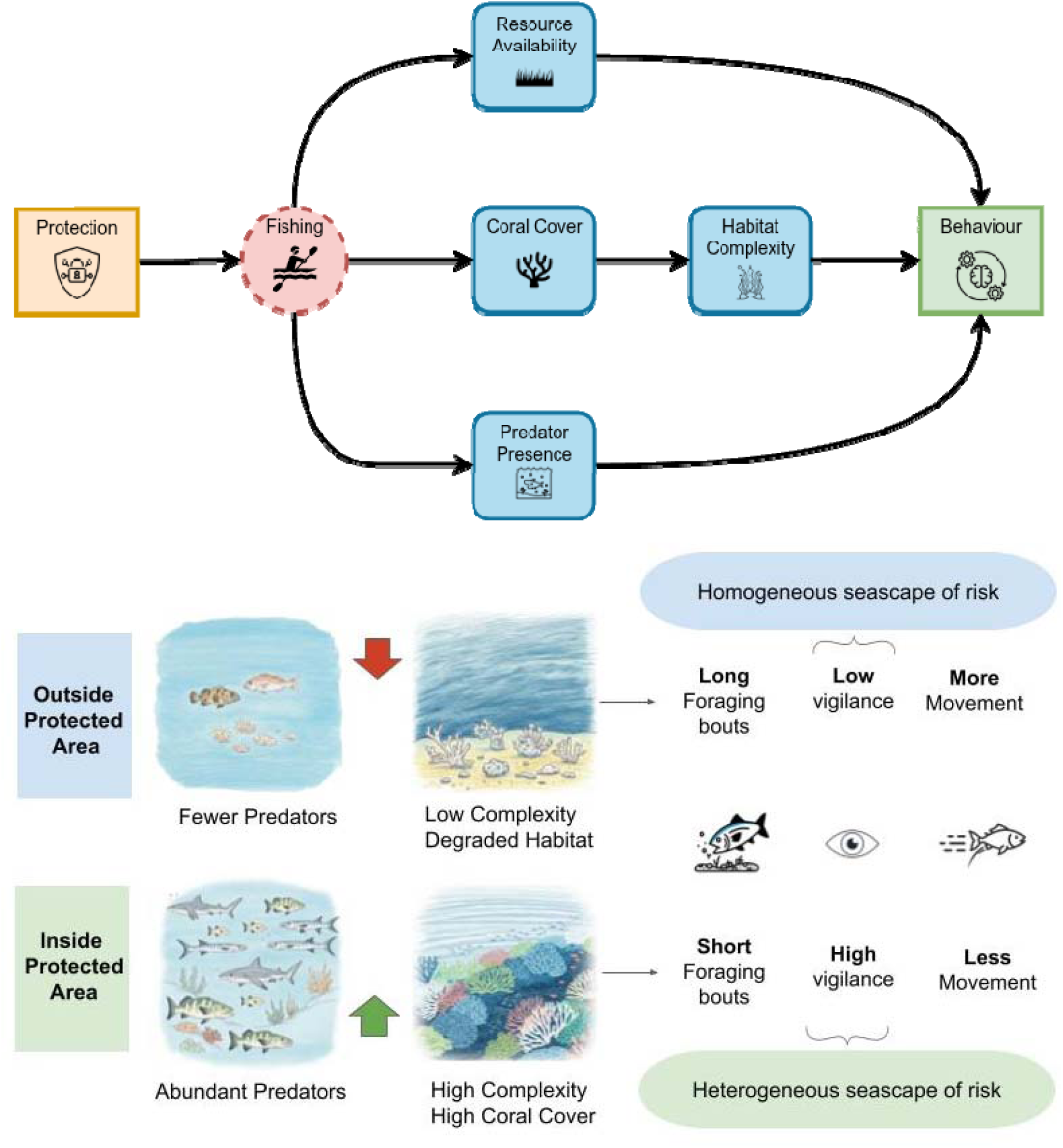
The effect of fishing and protection on predator assemblages, habitat and prey behaviour on coral reefs. Marine protected areas restrict access to fishing allowing predator assemblages and coral cover to recover, which has cascading downstream effects on reef fish behaviour (above). Fished reefs have fewer predators and more degraded habitats that lead to a homogenised seascape of fear. No take protected areas relieve the effects of fishing and human disturbance leading, thus predators are more abundant and habitats are more complex within protected areas, leading to a heterogeneous seascape of risk. As a consequence, prey fish species may forage for shorter durations at higher rates, be more vigilant and move less within than outside protected areas (below).

We first compared the abundance and size distribution of piscivorous fish (predators) across the sites inside and outside protected areas to determine differences in predation pressure due to fishing. We also compared the composition of benthic habitats across sites to determine whether protection influenced habitat complexity and resource availability. We determined the direct effect of protection on the foraging, vigilance and movement behaviour of herbivorous and invertivorous fish (prey) by comparing behavioural scans and time budgets across sites within and outside protected areas. Finally, we determined the effect of environmental conditions such as resource availability and habitat complexity on foraging, vigilance and movement behaviour of fish to quantify the indirect effects of protection on prey behaviour. We combine these results to discuss the impact of fishing on fish communities, behaviour and trophic dynamics.

## Methods

### Site selection

The study was conducted in the South Andaman Islands (India), primarily in and around Wandoor and Ritchie’s Archipelago (Figure 1). We selected 12 replicate sites based on previous surveys of coral cover and fish assemblages within and outside Mahatma Gandhi Marine National Park (MGMNP) near Wandoor and Rani Jhansi Marine National Park (RJMNP) situated north of Ritchie’s Archipelago. Fisheries in the Andaman and Nicobar Islands (ANI) primarily target predatory fish such as tuna, groupers, and snappers. The fishers in the area utilise mechanized boats and diverse artisanal gear such as handlines, shore seines, and gill nets, accommodating lucrative catch like. Exports have expanded rapidly in the past decade due to demand from domestic and international markets. The areas of Wandoor and Ritchie’s Archipelago are fished moderately, mainly with gillnets and, hook and line (21). Both MGMNP and RJMNP restrict fishing within their boundaries.

### In–water sampling

At each site we conducted estimation of predator assemblages, habitat complexity and collected video data for prey behavioural assays.

#### a) Characterising predator assemblages

Baited Remote Underwater Video Stations (BRUVS) were used at all sites at depth of 7-12 meters for a standardized 60-minute soak time (22–24). Videos were recorded in a GoPro camera and transcribed manually to determine species composition, relative abundance, size structure, and functional groups. We used the maximum number of individuals observed in a single frame (MaxN) as a proxy for relative abundance, which is a conservative estimate that avoids repeat counts (24–26). The total length (TL) of each individual fish was estimated against a calibrated 5 cm PVC pipe on the bait arm and categorized into 5 cm size classes (27). Finally, each species was assigned to a functional group (e.g., piscivore, invertivore) based on dietary information from FishBase (28),

#### b) Characterising benthic habitats

We sampled benthic cover of coral, algae and other substrates using on two 50m transects at each site. We placed and photographed twenty 50cm square quadrats at equal intervals on alternating sides of the transect line. We then used CPCe to determine the relative cover of different substrate types (29).

#### c) Characterising prey behaviour

For our behavioural sampling, we randomly selected five plots at each site with varying percentage cover of algae and variability in habitat complexity (rugosity) among plots. Each plot comprised a 1 x 1 m quadrat placed on the substrate of chosen reef patches. Plots were placed a minimum of 5 m apart. We estimated the resource abundance at each plot using photographs of the entire plot. We then used CPCe to determine the relative cover (%) of corals, sponges, algae and other substrates (sand, rock, rubble)(29).

We used remote underwater video surveys (RUVS) to observe fish behaviour. We deployed a GoPro camera within each sample plot to record herbivore fish bite rates and time budgets. Each camera has a field of view that covers approximately 2 m, we thus demarcated a 1 × 1 m^2^ quadrat for the purpose of analysis (14,30). All behavioural observations were conducted between 8:00 AM and 12:00 PM to account for changes in fish activity through the day (31).

After recording videos for behavioural observations, we estimated the rugosity of each plot using the chain transect method. We laid a 190 cm metal chain over the plot. As the chain is flexible, it adapts to the shape of the substrate thus changing the distance covered over the ground. Rugosity is the estimated ratio of straight-line length covered on the ground and the total length of the chain (32). We took three rugosity measurements at different angles per sample plot and used the mean of these measurements for analysis.

### Behavioural assays

We used videos captured from our in–water sampling to determine fish occurrence. We discarded the first 10 minutes of each recording to account for any disturbance caused during deployment of the cameras (14). We took five random 2-minute samples of the remaining video for processing for a total of 10 minutes sampled at each plot. We identified and counted both herbivore and predator species appearing within those samples of footage. These videos were further analysed using BORIS software for behavioural traits, viz, bite rates, foraging time, vigilance time and movement time (14).

We estimated the size class of sampled individuals visually using the quadrat sides (1m) as reference in steps of 10 cm (14,30). We quantified fish behaviour at three levels: (1) activity-states across individuals, (2) activity-states of individuals using time budgets, and (3) individual bite rates. Behaviours were classified into foraging, vigilance, and movement (Table 1). Activity states across individuals are expressed as the probability of individuals in the plot being seen in a particular state. For this, we recorded each individual’s behavioural state using scan samples taken 10–30 s after the individual first appeared in the video. The probability of each behavioural state was then calculated as the proportion of individuals exhibiting that state within a plot. Second, individual time budgets represented the proportion of total observation time spent in each behavioural state. This was estimated for individuals observed for more than 45s. Finally, when individuals were observed foraging, we additionally counted all bites within each foraging bout to estimate bite rates. Additionally, we noted any aggressive behaviour by the focal individuals against predators, conspecifics and heterospecifics. We did not observe any predation events or any disturbance due to human activities such as fishing or SCUBA diving. We also noted whether individuals were foraging in a group, and recorded group composition.

**Table 1:** Ethogram describing definitions of behaviours utilised for behavioural assays.

| Behavior code | Behavior type | Description |
| --- | --- | --- |
| Bite | Event | Pecking or biting at substrate |
| Vigilant | State | Head up, not feeding, focused or scanning |
| Moving | State | Displacement at a moderate pace |
| Escape | Event | Rapid displacement, finning |
| Agression | Event | Rapid displacement towards a target |
| Foraging | State | State of foraging: consecutive bites less than 5 seconds apart |

### Data Analysis

We used a causal inference framework along with Bayesian estimation to model predator assemblages, habitat composition and fish behaviour (31,32). Counterfactual simulations based on fitted models allowed us to estimate the effect of predictors by simulating outcomes under unobserved conditions (32). All models used sceptical priors for intercepts and coefficients (*Normal(0,0.5)*), and appropriate weak priors for standard deviations, scale, precision, and shape parameters (*Exponential*(*2*)) reflecting the prior assumption that predictors (e.g. protection, rugosity, etc.) had no influence on outcome variables (e.g. proportion of time spent vigilant, bite rates, etc.). For each estimate, we report the posterior probability of superiority—the probability that a parameter is greater than zero. Values near 0.5 indicate little or no effect, while values above 0.9 or below 0.1 indicate strong positive or negative effects, respectively. Model fit was assessed using Bayesian R^2^ which is measure of goodness of fit for Bayesian linear models (33). Further details on causal inference and counterfactual analyses are provided in Appendix A. All analyses were conducted in R 4.5.0 using the brms package (34, 35, 36).

#### a) Modelling predator assemblages and habitat composition

We modelled the abundance (maxN) of piscivorous fish as a Poisson process with a log link function, using protection status and island location (East/West) as predictors, and included a site-level random intercept (23). Fish size was modelled as an ordered categorical response with a cumulative logit link function and the same predictors (37). Counterfactual simulations were used to estimate the effect of protection on abundance and size, and results were expressed as log response ratios (LRR) comparing predicted values inside and outside protected areas (See appendix C for summary of model parameters).

We modelled the relative cover of corals, algae, sponges, and other substrates using a Dirichlet regression with a logit link function (39). The Dirichlet distribution describes the probability density of vectors that sum to one (simplex) and is hence appropriate for our benthic cover data. Protection status and location were included as predictors, with transects nested within sites as random effects. Counterfactual simulations provided estimates of protection effects on substrate composition, expressed as differences in predicted proportions between protected and unprotected sites (See appendix C for summary of model parameters).

#### b) Modelling fish behaviour

We modelled fish behaviour as four different processes: the abundance of fish on the plot (N = 55, Appendix D), the probability that each individual on the plot engages in either foraging, vigilance or movement behaviour (N = 239, Appendix E), the proportion of time that a fish individual engages in a given behaviour (N = 111, Appendix F) and finally the rate at which fish individuals forage (N = 56, Appendix G). For each process, we conducted counter-factual simulations to determine the direct effect of protection as well as the effect of plot – level and individual level predictors. Fish counts were over-dispersed and modelled with a negative binomial regression (log link). Predictors included protection status, rugosity, and biomass cover, with site as a random effect. We included the presence of damsel fish as a predictor for the abundance of fish on the plot. Guild identity and interactions with habitat variables were included as covariates. Protection effects were expressed as LRRs.

First, we modelled activity states across individuals from scan sampling data as the probability that herbivorous or invertivorous fish exhibited either foraging, movement, or vigilance behaviour. We modelled the probability of foraging, vigilance, or movement as separate Bernoulli processes with a logit link. Predictors included protection, habitat variables, predator presence, grouping status, guild, and size class. Plot and species (nested within family) were random effects. Effects were expressed as log odds ratios (LOR).

We then used focal individual sampling to determine the proportion of time spent in each state from individual time budgets. Proportions of time spent on each behaviour were modelled using an ordered beta regression to account for zero- and one-inflation (40). Predictors and random effects were the same as above. Effects were expressed as LORs.

Finally, for individuals that foraged, bite rates were modelled as a gamma process with a log link. Predictors and random effects were as above, and effects were reported as LRRs. All results are presented as probability (activity states), proportion (time budgets) and foraging rates (bite rates).

## Results

We recorded a total of 31 station-hours of video from BRUVS at 16 sites within and outside RJMNP and MGMNP. We sampled 24 transects for benthic cover and 55 plot-hours of videos for behavioural assays at 12 sites. Across the BRUVS, we recorded 837 fish individuals belonging to 72 species. Behavioural observations using RUVs were conducted on 434 individuals from 111 species. Among these, 46 species were invertivores (n = 160), 45 species were herbivores (n = 185) and 19 species were piscivores (n = 85). Additionally, 49 individuals were encountered as part of a group.

### Effect of protection on predator assemblages and habitats

Predator abundance was significantly greater within protected areas than outside protected areas (*LRR*=0.51[-0.008,1.013J, *P*(*LRR*>0)=0.947). However, the relative proportion of individuals across size classes did not differ across sites within and outside protected areas (Table 5 of appendix C). Coral cover was greater at sites within protected areas compared to outside protected areas (*δp*=16.1%(-30.3,63.9), *P*(*δp*>0)=0.736). In addition, the relative proportion of algae was lower within protected areas compared to outside protected (*δp*=-16.2%(-65.3,37.5), *P*(*δP*>0)=0.309).

### Direct effect of protection on fish behaviour

The number of herbivores that were encountered during behavioural assays did not differ at sites within and outside protected areas (*LRR*=-0.096(-0.636,0.446), *P*(*LRR*>0)=0.381). However, there were significantly fewer invertivores (*LRR*=-0.434(-1.080,0.201), *P*(*LRR*>0)=0.125) within protected areas than outside. Additionally, the number of piscivores encountered during behavioural assays was slightly lower inside protected areas compared to outside (*LRR*=-0.375(-1.115,0.377), *P*(*LRR*>0)=0.217).

The probability of fish exhibiting vigilance was greater within protected areas than outside protected areas (*LOR*=0.861(-0.189,1.958), *P*(*LOR*>0)=0.913). On the other hand, the probability of movement was lower within protected areas (*LOR*=-0.817(-1.977,0.266), *P*(*LOR*>0)=0.106). The probability of foraging did not change significantly with protection (*LOR*=0.181(-0.776,1.22), *P*(*LOR*>0)=0.625, Figure 2).

**Figure 2:**
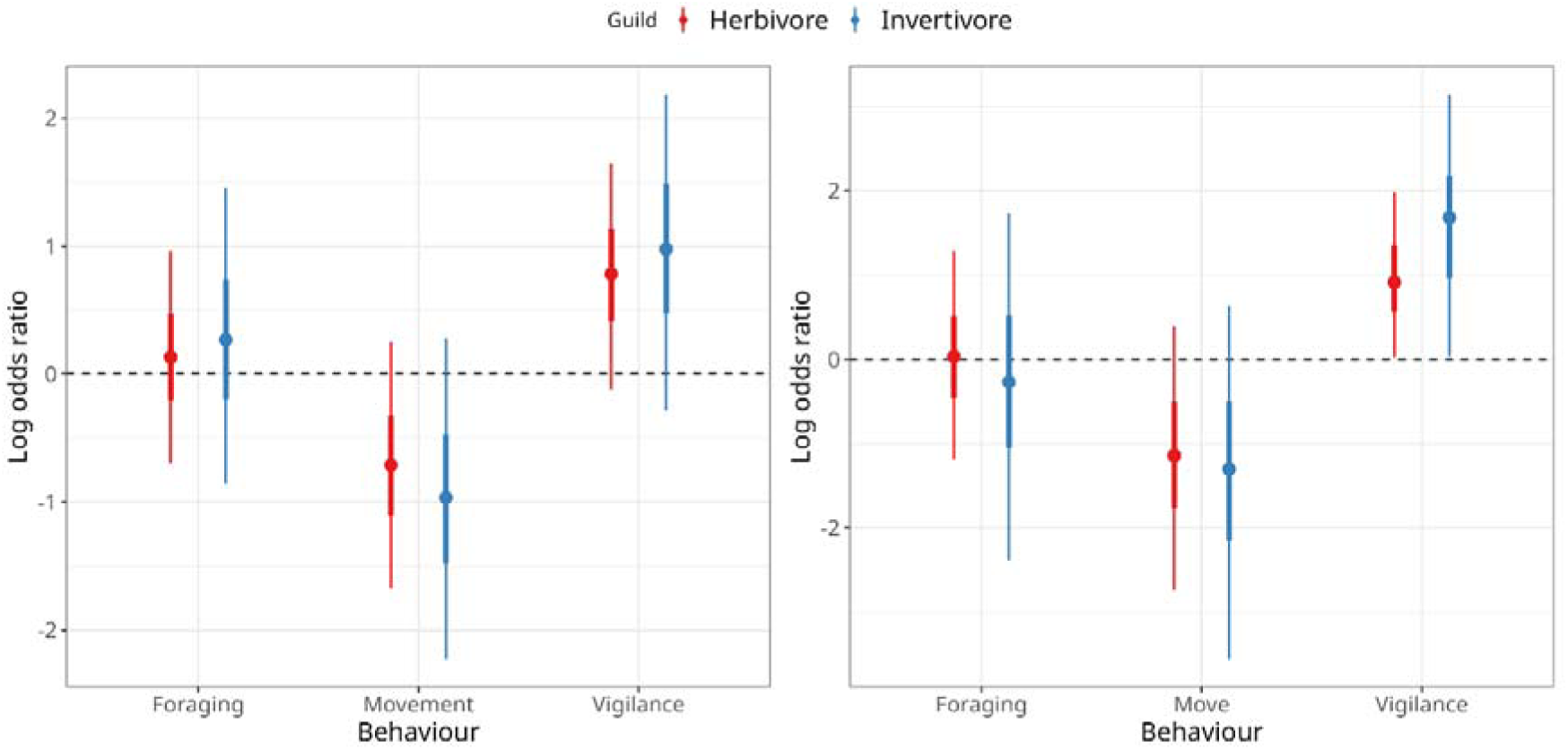
Left: Effect of protection on the proportion of fish exhibiting foraging, movement, or vigilance behaviour. Right: Effect of protection on the proportion of time spent in each behaviour. Points show the median posterior effect size; thick and thin lines indicate 50% and 90% credible intervals, respectively. The dashed line marks no effect. Points above it indicate higher likelihood or time spent in a behaviour inside protected areas, and points below indicate higher values outside protected areas.

The proportion of time fish spent vigilant was also significantly greater within protected areas (*LOR*=1.130(-0.142,2.562), *P*(*LOR*>0)=0.931). Similarly, the proportion of time fish spent moving was significantly lower within protected areas (*LOR*=-1.114(-2.914,0.477), *P*(*LOR*>0)=0.119). The proportion of time fish spent foraging was equal within and outside protected areas (*LOR*=-0.023(-1.511,0.532), *P*(*LOR*>0)=0.512, Figure 2). However, fish foraged at higher rates within protected areas (*LRR*=0.488(-0.255,1.726), *P*(*LRR*>0)=0.846, Figure 2). In addition, the effect of protection on foraging rates of invertivores was greater than that of herbivores (Table 1 of Appendix H).

### Effect of rugosity and algal cover on fish behaviour

The number of herbivores and invertivores encountered during behavioural assays increased significantly with increasing rugosity on the plot. However, the number of piscivores did not change (Table 3 of appendix D). Notably, the number of herbivores decreased significantly with increasing algal cover. The number of piscivores also decreased with algal cover but to a lesser extent. The number of invertivores did not change with algal cover (Table 5 of appendix D, Figure 3).

**Figure 3:**
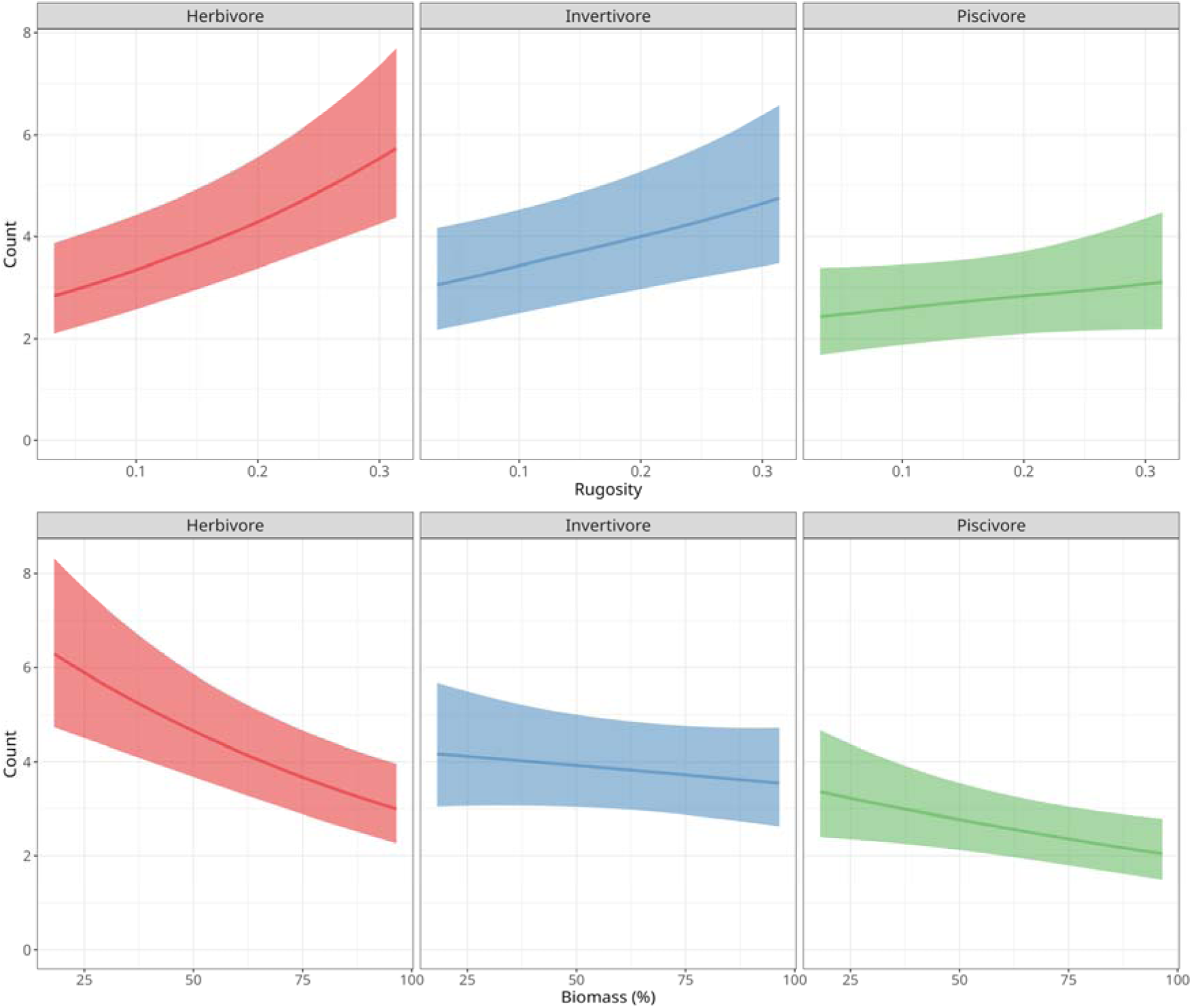
Effect of habitat complexity (rugosity) and resource availability (percentage cover of biomass) on the number of herbivores and invertivores observed in plots. Line indicates median posterior predictions of the regression model. Ribbons indicate 50% posterior credible intervals.

The probability of invertivores foraging was slightly higher and moving was slightly lower in areas with higher rugosity, but rugosity did not have a significant effect on the probability of foraging and movement of herbivores. In addition, the probability of vigilance of herbivores and invertivores did not change with rugosity (Table 2 of Appendix H). The proportion of time invertivorous fish spent foraging increased slightly with rugosity. However, herbivore foraging was not affected by rugosity (Table 3 of Appendix H). Similarly, the proportion of time fish spent moving decreased significantly for invertivores but not for herbivores (Table 3 of Appendix H). There was no effect of rugosity on the vigilance time of herbivores but invertivores were slightly more vigilant in more complex habitats (Table 3 of Appendix H). Invertivores also foraged at significantly higher rates in more complex sites, but herbivores did not (Table 4 of Appendix H).

The probability of foraging increased slightly with algal cover and probability of movement decreased slightly for both herbivores and invertivores. The probability of vigilance of both herbivores and invertivores did not change with algal cover (Table 5 of Appendix H). The proportion of time spent foraging and vigilant did not change significantly for both herbivores and invertivore. The proportion of time spent moving by herbivores did not change significantly with algal cover. However, the proportion of time spent moving by invertivores significantly decreased with algal cover (Table 6 of Appendix H). Additionally, the foraging rate of both herbivores and invertivores did not change significantly with algal cover (Table 7 of Appendix H).

### Effect of damselfish presence, predator presence and grouping on fish behaviour

The presence of damselfish on the plot had an overall negative effect on the number of fish on the plot (*LRR*=-0.457(-0.852,0.075), *P*(*LRR*>0)=0.024). Fish were less likely to forage when predators were present around the plot (Table 8 of Appendix H). The probability of vigilance and movement did not change significantly with predator presence (Table 8 of Appendix H). Similarly, the proportion of time fish spent moving and vigilant did not change significantly with predator presence (Table 9 of Appendix H). But fish individuals foraged in shorter bouts when predators were present (Table 9 of Appendix H). Overall, fish also foraged at slightly higher rates in the presence of predators (*LRR*=0.386(-0.21,0.995), *P*(*LRR*>0)=0.862).

Fish in groups were more likely to forage and foraged for longer bouts (Table 10 and 11 of Appendix H). Fish in groups were slightly less likely to be vigilant and spent a significantly lower proportion of time being vigilant. Grouping had no effect on fish movement behaviour (Table 10 and 11 of appendix H). In addition, fish in groups also foraged at lower rates (*LRR*=-0.29(-1.07,0.53), *P*(*LRR*>0)=0.274).

## Discussion

In the present study, we highlight how fishing alters the seascape of risk on coral reefs directly by altering predator assemblages and indirectly by altering habitats. To this end, we compared predator assemblages, habitat composition, and prey behaviour across two no-take marine protected areas (MPAs) and adjacent fished reefs in the Andaman Islands. We found greater piscivorous predator abundance and greater coral cover within protected areas. In addition, we found that reef-associated herbivorous and piscivorous fish were more likely to be vigilant and less likely to move within protected areas. Similarly, fish spent more time being vigilant and less time moving within protected areas (Figure 2). Prey fish species also foraged at higher rates within protected areas. Our findings underscore the importance of considering both consumptive and non-consumptive predator effects when evaluating the ecological impacts of human disturbance on ecosystems.

### Effect of fishing on predator abundance and habitat structure

Large bodied predators are economically valuable and are thus disproportionately targeted by fisheries (2). Thus, no–take restrictions tend to allow MPAs support greater predator biomass when effectively enforced (44). We found that predator abundance was significantly greater inside protected areas than on fished reefs, although predator size distributions did not differ between sites. This finding is consistent with earlier studies showing that MPAs enhance predator biomass but that changes in size structure often lag behind due to the long generation times and slow recovery rates of large-bodied reef predators (42–44). The absence of clear differences in size distributions may also signal ongoing fishing pressure adjacent to MPA boundaries, leading to possible spillover effects that prevent size-class recovery (41). The MPAs in our sites were established within the last 40 years (MGMNP in 1983 and RJMNP in 1996) and are relatively small (MGMNP: 281.5 km^2^; RJMNP: 256 km^2^). While our findings highlight an increase total predator presence, the capacity of MPAs to restore natural predator assemblage structures may require longer time periods or larger contiguous no-take zones.

We also observed that coral cover was consistently greater inside MPAs while algal cover was correspondingly higher at fished sites. This pattern reflects the broader literature demonstrating the role of protection in maintaining coral-dominated reef states by reducing fishing pressure on herbivores and preserving ecological interactions that suppress macroalgal proliferation (47,48). Although we expected lower predator biomass outside MPAs to release herbivores from predation and thus enhance herbivory (Figure 1), the higher algal cover we observed on fished reefs suggests that other stressors such as physical damage from fishing gear, anchor impacts, or slower coral recruitment may be limiting coral recovery in these areas. In contrast, coral recovery within MPAs likely enhances structural complexity, providing refuge for small-bodied fishes and prey groups. (46). Taken together, these results confirm that MPAs play a critical role not only in predator restoration but also in conserving benthic complexity and habitat quality—both of which mediate risk perception and subsequent prey behaviour.

### Direct effects of protection on prey behaviour

According to the risk–allocation hypothesis, prey must trade off the need of energy acquisition with threat of predation risk (20,50). Inside MPAs, higher predator abundance imposes elevated predation risk. Thus, prey individuals must respond by increasing vigilance. Simultaneously, reduced movement inside MPAs may reflect a strategy to minimize detection by predators or to exploit local refuges more efficiently (49). We found that the probability of both herbivorous and invertivorous fish being vigilant was greater within protected areas. In addition, that fishes inside protected areas altered their behavioural time budgets, spending more time for vigilance and less time for movement relative to individuals outside no-take protected areas (Figure 2).

Even though total foraging time did not change significantly between protected and fished sites, fish within MPAs foraged at higher rates. This suggests prey may compensate for increased vigilance costs by intensifying feeding bouts. Predator presence in and around our plots reduced the likelihood of foraging and shortened feeding bout duration with a corresponding increase in foraging effort. It has been found that prey adopt “burst foraging” strategies when predation risk is high, feeding rapidly yet cautiously to reduce exposure time to predators (53,56). Predator-driven trait-mediated effects such as these represent critical, non-consumptive forces that can cascade through reef trophic structures (55). Similar patterns of compensatory foraging effort under risk have been documented in coral reef fishes and other vertebrates where predators alter prey patch use or feeding intensities (51,52). Notably, invertivores showed greater sensitivity to predator-mediated behavioural shifts than herbivores (Figure 2). Variability across functional groups suggests that risk landscapes imposed by predators may structure prey communities differently, potentially influencing trophic dynamics and benthic cover trajectories.

### Effect of habitat complexity and resource availability on prey behaviour

Our findings emphasize the role of habitat complexity and resource availability in governing fish distributions and behaviours. Rugosity was positively associated with both herbivore and invertivore abundance during behavioural surveys (Figure 3). Thus, habitats with higher cover and thus higher structural complexity may provide refuge to prey individuals and thus be perceived as less risky (50,58). Additionally, structural complexity may alter the availability of certain food types such as invertebrates. Notably, invertivores were more likely to forage and less likely to move in highly rugose habitats. This trend was also reflected in the proportion of time spent in these states, likely reflecting their reliance on benthic microhabitats for prey capture. Herbivores, by contrast, appeared relatively less responsive to rugosity, possibly because their foraging is tied more closely to algal resource distribution than structural refuge (Figure 4).

**Figure 4:**
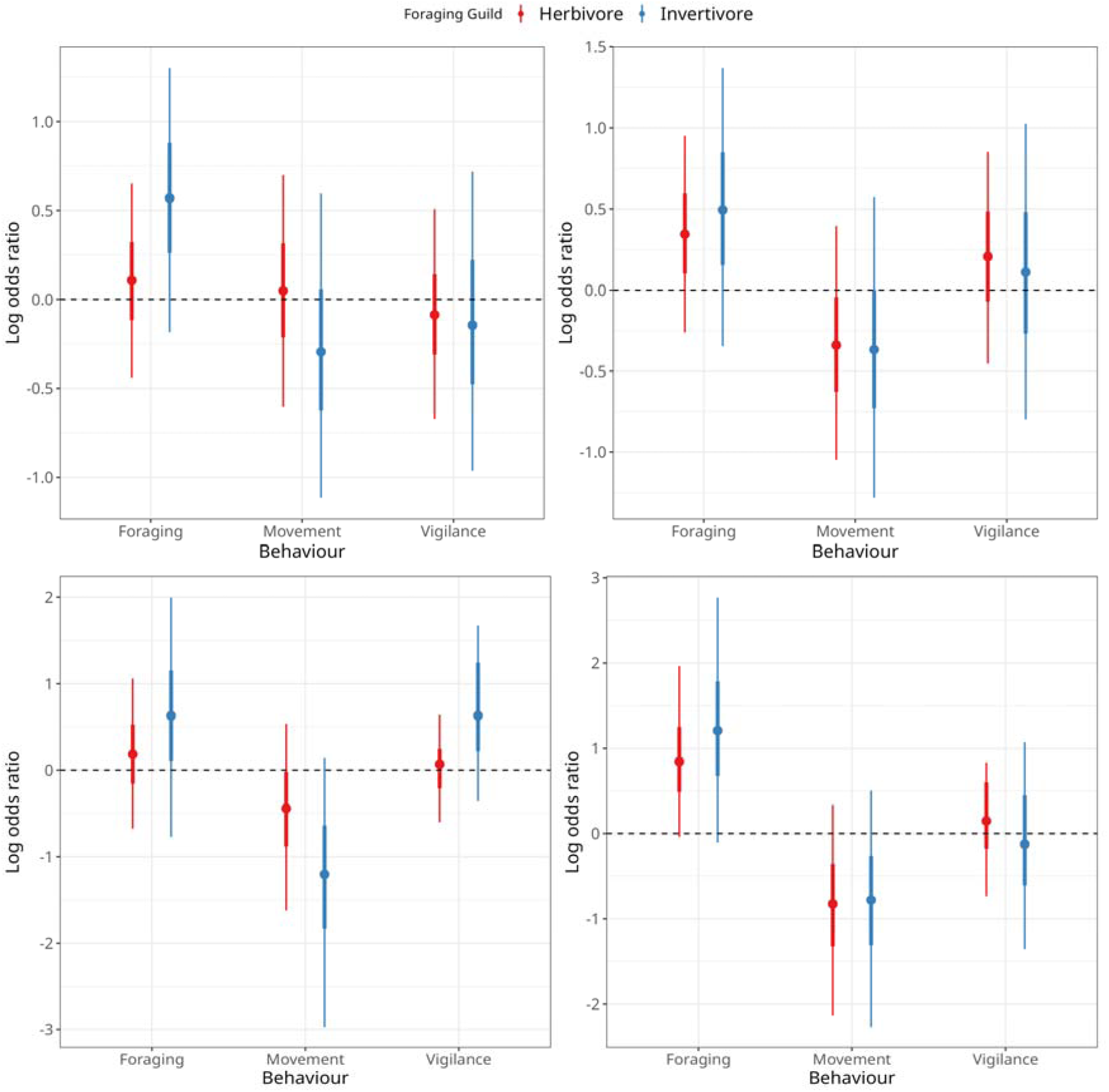
Top-left: Effect of habitat complexity (rugosity) on the proportion of fish exhibiting foraging, movement, or vigilance behaviour. Top-right: Effect of habitat complexity (rugosity) on the proportion of time spent in each behaviour. Bottom-left: Effect of resource availability (algal cover) on the proportion of fish exhibiting foraging, movement, or vigilance behaviour. Bottom-right: Effect of resource availability (algal cover) on the proportion of time spent in each behaviour. Points show the median posterior effect size; thick and thin lines indicate 50% and 90% credible intervals, respectively. The dashed line marks no effect. Points above it indicate higher likelihood or time spent in a behaviour inside protected areas, and points below indicate higher values outside protected areas.

Resource availability had a markedly different effect on the behaviour of fish than habitat complexity (Figure 3). We found significantly fewer herbivores in areas with high algal cover. Additionally, damselfish territoriality exerted an independent influence by reducing the overall number of fish on plots. Damselfishes, as ecosystem engineers, monopolize algal resources and exclude other herbivores or planktivores through aggressive territoriality (60). Damselfish presence therefore adds another layer of behavioural constraint, locally restructuring community assemblages independent of MPA protection status. Thus, productive algal-dominated reefs may paradoxically be less favourable for herbivore populations due to antagonistic interactions with territorial damselfish, declines in palatable algal taxa, or physiological constraints on diet (45).

Invertivores did not show strong behavioural responses to algal cover. This suggests that while habitat degradation alters herbivory, it may not directly constrain invertivorous feeding guilds. The complex interplay between habitat complexity and resource availability suggests that fishing alters predator-prey dynamics indirectly through habitat degradation. This highlights the multifaceted mechanisms through which fishing shapes coral reef ecosystems

### Individual variability in prey behaviour

Fish in groups were significantly less likely to be vigilant and more likely to forage than solitary fish. Similarly solitary fish spent less time foraging and more time being vigilant, consistent with the dilution effect and benefits of collective group defence highlighted across many taxa (59). These findings indicate that grouping acts as a natural behavioural buffer against predation risk, enabling individuals to allocate more time to foraging and less to anti-predator vigilance. Grouped individuals foraged at lower rates, despite foraging more often and for longer. This suggests that collective feeding reduces the urgency of individual consumption, reflecting a trade-off between group foraging consistency and individual foraging effort.

### Conclusion

Taken together, these results illustrate how fishing reshapes the seascape of risk directly by altering predator distributions, and indirectly by altering habitats (49). On fished reefs where predators are scarce, prey fish displayed reduced vigilance, greater mobility, and different foraging strategies. Such conditions reflect an altered landscape of risk where predation threat is minimal, enabling prey to forage openly with less need for vigilance and refuge. Conversely, in MPAs where predator abundance is higher, prey exhibited heightened vigilance and more risk-sensitive foraging behaviours (49). In this sense, MPAs reinforce the landscape of fear and constrain prey behaviour, even beyond actual predation events (52).

Importantly, these behavioural shifts can have cascading ecological consequences. Increased vigilance and altered foraging strategies inside MPAs could modify patterns of algal grazing and invertebrate consumption, influencing benthic community trajectories. Conversely, fished reefs, by lacking such non-consumptive effects, may experience more uniform foraging but less structural resilience. Thus, protection both restores consumptive predator–prey interactions (direct predation) and reinstates non-consumptive trait-mediated effects, thereby reinforcing trophic feedbacks that maintain coral reef resilience (58).

This study underscores the dual role of MPAs in conserving both biological assemblages and behavioural processes that regulate ecosystem function. Ultimately, fishing reduces predator biomass and alters benthic habitats, while protection restores habitat structure and predator presence, emphasising behavioural trade-offs and altering ecological interactions. Collectively, these findings emphasize that understanding the full impact of extraction on reef ecosystems requires accounting for both its immediate and cascading effects on fish assemblages.

## Data accessibility

Data and code for the paper is available at https://doi.org/10.5281/zenodo.17617857

## Acknowledgements

This manuscript is dedicated to Dr. Anne Heloise Theo, whose inputs were invaluable in the design of this study. We’d like to thank all the staff at Andaman Nicobar Environment Team (ANET), especially our boat captains and crew; Babu Kutty, Saw Thesorow, Jeevan Horo, Saw Watha (Agu) and Sebian Horo, for their logistical support. We thank Rahul Demello, Tanmay Wagh, Esha Gokhale, and Chaitanya Arjunwadkar for their suggestions on methods and assistance during field work. Finally, we’d like to thank the Andaman and Nicobar Islands forest department, the MGMNP ranger office, the RJMNP ranger office, and ANI Fisheries Department for providing us with permits to conduct our work.

## References

1. Estes JA, Terborgh J, Brashares JS, Power ME, Berger J, Bond WJ, et al. Trophic Downgrading of Planet Earth. Science. 2011 July 15;333(6040):301–6.

2. Jackson JBC, Kirby MX, Berger WH, Bjorndal KA, Botsford LW, Bourque BJ, et al. Historical Overfishing and the Recent Collapse of Coastal Ecosystems. Sci New Ser. 2001;293(5530):629–38.

3. Schmitz OJ, Wilmers CC, Leroux SJ, Doughty CE, Atwood TB, Galetti M, et al. Animals and the zoogeochemistry of the carbon cycle. Science. 2018 Dec 7;362(6419):eaar3213.

4. Treves A, Naughton-Treves L. Risk and opportunity for humans coexisting with large carnivores. J Hum Evol. 1999;36(3):275–82.

5. Darimont CT, Fox CH, Bryan HM, Reimchen TE. The unique ecology of human predators. Science. 2015 Aug 21;349(6250):858–60.

6. Brown JS, Kotler BP. Hazardous duty pay and the foraging cost of predation. Ecol Lett. 2004;7(10):999–1014.

7. Lima SL, Dill LM. Behavioral decisions made under the risk of predation: a review and prospectus. Can J Zool. 1990 Apr;68(4):619–40.

8. Madin EMP, Harborne AR, Harmer AMT, Luiz OJ, Atwood TB, Sullivan BJ, et al. Marine reserves shape seascapes on scales visible from space. Proc R Soc B Biol Sci. 2019;286(1901):20190053.

9. Laundre JW, Hernandez L, Ripple WJ. The Landscape of Fear: Ecological Implications of Being Afraid∼!2009-09-09∼!2009-11-16∼!2010-02-02∼! Open Ecol J. 2010 Mar 5;3(3):1–7.

10. Atwood TB, Madin EM, Harborne AR, Hammill E, Luiz OJ, Ollivier QR, et al. Predators shape sedimentary organic carbon storage in a coral reef ecosystem. Front Ecol Evol. 2018;6:110.

11. Schmitz OJ, Krivan V, Ovadia O. Trophic cascades: the primacy of trait-mediated indirect interactions: Primacy of trait-mediated indirect interactions. Ecol Lett. 2004 Feb 4;7(2):153–63.

12. Schmitz OJ, Beckerman AP, O’Brien KM. Behaviorally Mediated Trophic Cascades: Effects of Predation Risk on Food Web Interactions. Ecology. 1997;78(5):1388–99.

13. Wirsing AJ, Heithaus MR, Brown JS, Kotler BP, Schmitz OJ. The context dependence of non-consumptive predator effects. Ecol Lett. 2021;24(1):113–29.

14. Bauman AG, Seah JCL, Januchowski-Hartley FA, Hoey AS, Fong J, Todd PA. Fear effects associated with predator presence and habitat structure interact to alter herbivory on coral reefs. Biol Lett. 2019 Oct 31;15(10):20190409.

15. Catano LB, Barton MB, Boswell KM, Burkepile DE. Predator identity and time of day interact to shape the Risk–Reward trade-off for herbivorous coral reef fishes. Oecologia. 2017 Mar 1;183(3):763–73.

16. Lehtonen J, Jaatinen K. Safety in numbers: the dilution effect and other drivers of group life in the face of danger. Behav Ecol Sociobiol. 2016 Apr;70(4):449–58.

17. Bhathal B, Pauly D. “Fishing down marine food webs” and spatial expansion of coastal fisheries in India, 1950-2000. Fish Res. 2008;91(1):26–34.

18. Atwood TB, Madin EMP, Harborne AR, Hammill E, Luiz OJ, Ollivier QR, et al. Predators Shape Sedimentary Organic Carbon Storage in a Coral Reef Ecosystem. Front Ecol Evol. 2018;6:110.

19. Madin EMP, Harborne AR, Harmer AMT, Luiz OJ, Atwood TB, Sullivan BJ, et al. Marine reserves shape seascapes on scales visible from space. Proc R Soc B Biol Sci. 2019 Apr 24;286(1901):20190053.

20. Lima SL, Bednekoff PA. Temporal Variation in Danger Drives Antipredator Behavior: The Predation Risk Allocation Hypothesis. Am Nat. 1999 June 1;153(6):649–59.

21. Jaini M, Advani S, Shanker K, Oommen MA, Namboothri N. History, culture, infrastructure and export markets shape fisheries and reef accessibility in India’s contrasting oceanic islands. Environ Conserv. 2018;45(1):41–8.

22. Westerberg H, Westerberg K. Properties of odour plumes from natural baits. Fish Res. 2011 Aug 1;110(3):459–64.

23. Dorman SR, Harvey ES, Newman SJ. Bait Effects in Sampling Coral Reef Fish Assemblages with Stereo-BRUVs. PLOS ONE. 2012 July 27;7(7):e41538.

24. Whitmarsh SK, Fairweather PG, Huveneers C. What is Big BRUVver up to? Methods and uses of baited underwater video. Rev Fish Biol Fish. 2017 Mar 1;27(1):53–73.

25. Grimmel HMV, Bullock RW, Dedman SL, Guttridge TL, Bond ME. Assessment of faunal communities and habitat use within a shallow water system using non-invasive BRUVs methodology. Aquac Fish. 2020 Sept 1;5(5):224–33.

26. Cappo M a, Harvey E, Malcolm H, Speare P, others. Potential of video techniques to monitor diversity, abundance and size of fish in studies of marine protected areas. Aquat Prot Areas-What Works Best We Know. 2003;1:455–64.

27. Harvey E, Fletcher D, Shortis M. Estimation of reef fish length by divers and by stereo-video: a first comparison of the accuracy and precision in the field on living fish under operational conditions. Fish Res. 2002;57(3):255–65.

28. Froese R, Pauly D. FishBase [Internet]. 2025. Available from: https://www.fishbase.org

29. Kohler KE, Gill SM. Coral Point Count with Excel extensions (CPCe): A Visual Basic program for the determination of coral and substrate coverage using random point count methodology. Comput Geosci. 2006;32(9):1259–69.

30. Catano LB, Rojas MC, Malossi RJ, Peters JR, Heithaus MR, Fourqurean JW, et al. Reefscapes of fear: predation risk and reef hetero-geneity interact to shape herbivore foraging behaviour. J Anim Ecol. 2016;85(1):146–56.

31. Karkarey R, Alcoverro T, Kumar S, Arthur R. Coping with catastrophe: Foraging plasticity enables a benthic predator to survive in rapidly degrading coral reefs. Anim Behav. 2017;131:13–22.

32. Kuffner IB, Brock JC, Grober-Dunsmore R, Bonito VE, Hickey TD, Wright CW. Relationships between reef fish communities and remotely sensed rugosity measurements in Biscayne National Park, Florida, USA. Environ Biol Fishes. 2007;78(1):71–82.

33. McElreath R. Statistical Rethinking: A Bayesian Course with Examples in R and Stan [Internet]. New York: Chapman and Hall/CRC; 2018. Available from: 10.1201/9781315372495.

34. Gelman A, Carlin JB, Stern HS, Rubin DB. Bayesian Data Analysis [Internet]. New York: Chapman and Hall/CRC; 1995. Available from: 10.1201/9780429258411.

35. Froese R, Pauly D. FishBase 2000. Concepts Des Data Sources. 2000;1594.

36. Team RC, others. R: A language and environment for statistical computing. R Foundation for Statistical Computing, Vienna, Austria. Httpwww R-Proj Org. 2016;

37. Bürkner PC. brms: An R package for Bayesian multilevel models using Stan. J Stat Softw. 2017;80:1–28.

38. Bürkner PC, Vuorre M. Ordinal regression models in psychology: A tutorial. Adv Methods Pract Psychol Sci. 2019;2(1):77–101.

39. Sennhenn-Reulen H. Bayesian regression for a Dirichlet distributed response using Stan. ArXiv Prepr ArXiv180806399. 2018;

40. Kubinec R. Ordered beta regression: a parsimonious, well-fitting model for continuous data with lower and upper bounds. Polit Anal. 2023;31(4):519–36.

41. Mumby PJ, Dahlgren CP, Harborne AR, Kappel CV, Micheli F, Brumbaugh DR, et al. Fishing, Trophic Cascades, and the Process of Grazing on Coral Reefs. Science. 2006 Jan 6;311(5757):98–101.

42. Russ GR, Alcala AC, Maypa AP, Calumpong HP, White AT. Marine reserve benefits local fisheries. Ecol Appl. 2004;14(2):597–606.

43. Edgar GJ, Stuart-Smith RD, Willis TJ, Kininmonth S, Baker SC, Banks S, et al. Global conservation outcomes depend on marine protected areas with five key features. Nature. 2014 Feb;506(7487):216–20.

44. Williamson D, Russ G, Ayling A. No-take marine reserves increase abundance and biomass of reef fish on inshore fringing reefs of the Great Barrier Reef. Environ Conserv. 2004;31(2):149–59.

45. Hughes TP, Rodrigues MJ, Bellwood DR, Ceccarelli D, Hoegh-Guldberg O, McCook L, et al. Phase shifts, herbivory, and the resilience of coral reefs to climate change. Curr Biol CB. 2007 Feb 20;17(4):360–5.

46. Mumby PJ, Hastings A. The impact of ecosystem connectivity on coral reef resilience. J Appl Ecol. 2008;45(3):854–62.

47. Graham NA, Nash KL. The importance of structural complexity in coral reef ecosystems. Coral Reefs. 2013;32(2):315–26.

48. Mumby PJ, Hastings A. The impact of ecosystem connectivity on coral reef resilience. J Appl Ecol. 2008;45(3):854–62.

49. Madin EMP, Madin JS, Booth DJ. Landscape of fear visible from space. Sci Rep. 2011;1:14.

50. Lima SL, Dill LM. Behavioral decisions made under the risk of predation: A review and prospectus. Can J Zool. 1990;68(4):619–40.

51. Werner EE, Peacor SD. A review of trait-mediated indirect interactions in ecological communities. Ecology. 2003;84(5):1083–100.

52. Catano LB, Rojas MC, Malossi RJ, Peters JR, Heithaus MR, Fourqurean JW, et al. Reefscapes of fear: Predation risk and reef hetero-geneity interact to shape herbivore foraging behaviour. J Anim Ecol. 2016;85(1):146–56.

53. Brown JS, Kotler BP. Hazardous duty pay and the foraging cost of predation. Ecol Lett. 2004;7(10):999–1014.

54. Rizzari JR, Frisch AJ, Hoey AS, McCormick MI. Not worth the risk: apex predators suppress herbivory on coral reefs. Oikos. 2014;123(7):829–36.

55. Peacor SD, Werner EE. The contribution of trait-mediated indirect effects to the net effects of a predator. Proc Natl Acad Sci U S A. 2001 Mar 27;98(7):3904–8.

56. Rizzari JR, Frisch AJ, Hoey AS, McCormick MI. Not worth the risk: apex predators suppress herbivory on coral reefs. Oikos. 2014 July;123(7):829–36.

57. Almany GR. Differential Effects of Habitat Complexity, Predators and Competitors on Abundance of Juvenile and Adult Coral Reef Fishes. Oecologia. 2004;141(1):105–13.

58. Catano L, Shantz A, Burkepile D. Predation risk, competition, and territorial damselfishes as drivers of herbivore foraging on Caribbean coral reefs. Mar Ecol Prog Ser. 2014 Sept 24;511:193–207.

59. Krause J, Ruxton GD, Krause J, Ruxton GD. Living in Groups. Oxford, New York: Oxford University Press; 2002. 228 p. (Oxford Series in Ecology and Evolution).

60. Ceccarelli DM. Modification of benthic communities by territorial damselfish: a multi-species comparison. Coral Reefs. 2007 Dec 1;26(4):853–66.

