## Supplementary Materials for "Local extraction of predatory fish may initiate behaviourally mediated trophic cascades in coral reefs in the Andaman Islands"

### Appendix A: Causal inference and counter-factual analysis

Causal inference is a set of statistical procedures used to infer causal connections between two or more variables. It stands distinct from correlation or association in that it seeks to identify mechanisms or pathways that induce a change in the outcome variable directly attributable to the treatment or exposure variable. In the context of our study, we used causal inference to assess how protection status and species traits influence fish assemblages and behaviours at our sites.

Directed Acyclic Graphs (DAGs) are employed to visually represent and clarify our causal assumptions. These graphs help in delineating causal pathways and ensuring that the model does not incorporate feedback loops that could invalidate causal interpretations. This visualisation aids in meticulously defining the structural form of the causal models, helping ensure the coherence of causal claims (Pearl, 2000).

Generative Models are developed by using the DAGs as a reference. These models portray the hypothesised process from which our data emerges, enabling us to derive simulated data sets. We use these generated outputs to test and validate our statistical estimators, ensuring robustness and accuracy before applying them to actual data scenarios.

Bayesian estimation is a probabilistic approach to inference, which integrates prior knowledge with the data to update our beliefs about model parameters in the form of posterior distributions. This framework is pivotal in causal inference as it facilitates the integration of various sources of uncertainty in a coherent manner. For our analyses, the Bayesian procedure aids in precisely estimating the parameters of our regression models (Gelman et al., 2013).

For practical implementation, the posteriors from our models, which are typically complex and high-dimensional, cannot be solved analytically. To manage this, we utilise sampling algorithms to characterise the posterior distribution. Markov Chain Monte Carlo (MCMC) techniques, particularly the Hamiltonian Monte Carlo (HMC) implementation in Stan, are used to accurately characterise the posterior distributions of our model parameters while ensuring efficient sampling from these complex distributions (Carpenter et al., 2017).

In counterfactual analysis, we consider hypothetical alternatives to the events that actually occurred. For example, we simulate how fish assemblage and behaviour might differ under various protection levels or environmental conditions that were not originally observed. This involves evaluating the outcomes under alternative scenarios where a key variable or treatment is modified. We use this approach to isolate and estimate the causal effects of interest, especially in situations where experimental manipulation is unfeasible or impractical. This approach is crucial for understanding how different factors might influence outcomes in natural settings (Morgan & Winship, 2014; Imbens & Rubin, 2015).

**References**

1. Pearl, J. (2000). Causality: Models, Reasoning, and Inference. Cambridge University Press.
2. Gelman, A., Carlin, J. B., Stern, H. S., Dunson, D. B., Vehtari, A., & Rubin, D. B. (2013). Bayesian Data Analysis. Chapman and Hall/CRC.
3. Carpenter, B., Gelman, A., Hoffman, M. D., Lee, D., Goodrich, B., Betancourt, M., ... & Riddell, A. (2017). Stan: A probabilistic programming language. Journal of Statistical Software, 76(1).
4. Morgan, S. L., & Winship, C. (2014). Counterfactuals and Causal Inference. Cambridge University Press.
5. Imbens, G. W., & Rubin, D. B. (2015). Causal Inference for Statistics, Social, and Biomedical Sciences. Cambridge University Press.

### Appendix B: Description of fish assemblages

**Table 1:** Summary of traits of fish species encountered during behavioural observations.

| **Order** | **Family** | **Species** | **Guild** | **Column** | **Habitat** | **Abundance** |
| --- | --- | --- | --- | --- | --- | --- |
| Perciformes | Pomacentridae | *Pomacentrus albicaudatus* | Invertivore | Benthic |  | 31 |
| Perciformes | Acanthuridae | *Ctenochaetus striatus* | Herbivore, invertivore | Bentho-pelagic | Reef flats, lagoons, seaward reefs | 29 |
| Perciformes | Nemipteridae | *Scolopsis bilineatus* | Invertivore, piscivore | Benthic | Sand and rubble fringe in reefs | 22 |
| Eupercaria | Scaridae | *Chlorurus sordidus* | Herbivore | Benthic | Coral reefs, rubble | 16 |
| Eupercaria | Scaridae | *Chlorurus sordidus* | Herbivore | Benthic | Coral reefs, rubble | 16 |
| Perciformes | Pomacentridae | *Pomacentrus xanthosternus* | Invertivore | Benthic |  | 14 |
| Eupercaria | Labridae | *Thalassoma lunare* | Invertivore | Benthic | Coastal, lagoons, outer reefs | 13 |
| Eupercaria | Labridae | *Thalassoma lunare* | Invertivore | Benthic | Coastal, lagoons, outer reefs | 13 |
| Perciformes | Acanthuridae | *Acanthurus tristis* | Herbivore | Bentho-pelagic | Lagoons, seaward reefs | 12 |
| Eupercaria | Labridae | *Halichoeres hortulanus* | Invertivore | Bentho-pelagic | Lagoons, seaward reefs | 12 |
| Eupercaria | Labridae | *Halichoeres hortulanus* | Invertivore | Bentho-pelagic | Lagoons, seaward reefs | 12 |
| Perciformes | Mullidae | *Parupeneus macronemus* | Invertivore |  |  | 12 |
| Eupercaria | Labridae | *Halichoeres dussumieri* | Invertivore | Benthic | Shallow weedy areas | 11 |
| Eupercaria | Labridae | *Halichoeres dussumieri* | Invertivore | Benthic | Shallow weedy areas | 11 |
| Eupercaria | Labridae | *Halichoeres bicolor* | Invertivore | Bentho-pelagic | Lagoons, outer reefs | 10 |
| Perciformes | Lutjanidae | *Lutjanus decussatus* | Invertivore, piscivore | Benthic | Coastal reefs, lagoons and outer slopes | 10 |
| Perciformes | Chaetodontidae | *Chaetodon trifasciatus* | Invertivore | Benthic |  | 9 |
| Perciformes | Pomacentridae | *Chrysiptera talboti* | Invertivore | Benthic | Seaward reef slopes and deep lagoons | 9 |
| Perciformes | Balistidae | *Balistapus undulatus* | Invertivore | Bentho-pelagic |  | 8 |
| Eupercaria | Labridae | *Cheilinus trilobatus* | Invertivore, piscivore | Benthic | Lagoons, outward reefs | 8 |
| Eupercaria | Labridae | *Cheilinus trilobatus* | Invertivore, piscivore | Benthic | Lagoons, outward reefs | 8 |
| Eupercaria | Labridae | *Halichoeres chierchiae* | Invertivore | Bentho-pelagic | Lagoons, outer reefs | 8 |
| Perciformes | Pomacentridae | *Dischistodus melanotus* | Herbivore | Benthic |  | 7 |
| Perciformes | Synodontinae | *Synodus variegatus* | Piscivore | Benthic |  | 7 |
| Perciformes | Lethrinidae | *Monotaxis grandoculis* | Invertivore | Bentho-pelagic | Coastal reefs, lagoons, outer slopes | 6 |
| Perciformes | Monacanthidae | *Nelusetta ayraud* | Piscivore, invertivore | Bentho-pelagic |  | 6 |
| Perciformes | Gobiidae | *Valenciennea strigata* | Piscivore, invertivore | Benthic |  | 6 |
| Pecriformes | Serranidae | *Cephalopholis boenak* | Invertivore, piscivore | Benthic | Silty reefs in sheltered waters, coastal reefs | 5 |
| Perciformes | Pomacentridae | *Pomacentrus amboinensis* | Herbivore, invertivore | Benthic | Lagoons, coastal reefs, passages and outer reef slopes | 5 |
| Perciformes | Pomacentridae | *Pomacentrus andamanensis* | Herbivore | Benthic |  | 5 |
| Eupercaria | Scaridae | *Scarus schlegeli* | Herbivore | Benthic | Coastal, lagoon, outer reefs | 5 |
| Eupercaria | Scaridae | *Scarus schlegeli* | Herbivore | Benthic | Coastal, lagoon, outer reefs | 5 |
| Perciformes | Balistidae | *Sufflamen chrysopterum* | Invertivore | Benthic |  | 5 |
| Perciformes | Acanthuridae | *Acanthurus lineatus* | Herbivore | Bentho-pelagic | Seaward reefs | 4 |
| Perciformes | Pomacentridae | *Amblygliphidodon silolona* | Herbivore | Benthic |  | 4 |
| Eupercaria | Labridae | *Anampses geographicus* | Invertivore | Benthic | Reef tops, slopes | 4 |
| Eupercaria | Labridae | *Anampses geographicus* | Invertivore | Benthic | Reef tops, slopes | 4 |
| Perciformes | Nemipteridae | *Scolopsis lineata* | Invertivore, piscivore | Benthic | Sand and rubble fringe in reefs | 4 |
| Perciformes | Balistidae | *Suflamen chrysoterum* | Invertivore | Benthic |  | 4 |
| Perciformes | Pomacanthidae | *Centropyge nox* | Herbivore, invertivore | Benthic |  | 3 |
| Eupercaria | Labridae | *Cheilinus abudjubbe* | Invertivore | Benthic | Coral areas of lagoons, coastal reefs | 3 |
| Perciformes | Pomacentridae | *Chromis viridis* | Herbivore | Benthic | Subtidal reef flats and lagoons | 3 |
| Perciformes | Pomacentridae | *Chromis weberi* | Invertivore | Benthic | Steep outer reef slopes | 3 |
| Perciformes | Pomacentridae | *Pomacentrus moluccensis* | Herbivore, invertivore | Benthic | Shoreline, lagoons and outer reefs | 3 |
| Perciformes | Pomacentridae | *Pomacentrus philippinus* | Herbivore, invertivore | Benthic | Passages and outer reefs | 3 |
| Eupercaria | Scaridae | *Scarus rivulatus* | Herbivore, invertivore | Benthic | Silty coastal reefs, lagoons, seaward reefs | 3 |
| Eupercaria | Scaridae | *Scarus rivulatus* | Herbivore, invertivore | Benthic | Silty coastal reefs, lagoons, seaward reefs | 3 |
| Perciformes | Nemipteridae | *Scolopsis aurata* | Invertivore, piscivore | Benthic | Sandy bottoms of lagoons, coastal reefs | 3 |
| Perciformes | Scaridae | *Sparisoma chrysopterum* | Herbivore | Bentho-pelagic |  | 3 |
| Eupercaria | Labridae | *Stethojulis trilineata* | Invertivore | Benthic | Shallow clear water reefs | 3 |
| Eupercaria | Labridae | *Stethojulis trilineata* | Invertivore | Benthic | Shallow clear water reefs | 3 |
| Perciformes | Zanclidae | *Zanclus cornutus* | Herbivore, invertivore | Bentho-pelagic |  | 3 |
| Perciformes | Pomacentridae | *Acanthochromis polyacanthus* | Herbivore, invertivore | Benthic | Inshore and offschore coral reefs | 2 |
| Eupercaria | Labridae | *Anampses elegans* | Invertivore | Benthic | Coral reefs, rocky reefs | 2 |
| Eupercaria | Labridae | *Anampses elegans* | Invertivore | Benthic | Coral reefs, rocky reefs | 2 |
| Eupercaria | Labridae | *Anampses meleagrides* | Invertivore | Benthic | Coral, rubble and sand of seaward reefs | 2 |
| Eupercaria | Labridae | *Anampses meleagrides* | Invertivore | Benthic | Coral, rubble and sand of seaward reefs | 2 |
| Perciformes | Pomacanthidae | *Centropyge multispinis* | Herbivore, invertivore | Benthic |  | 2 |
| Eupercaria | Scaridae | *Cetoscarus ocellatus* | Herbivore | Benthic | Lagoons, seaward reefs | 2 |
| Perciformes | Chaetodontidae | *Chaetodon oxycephalus* | Invertivore | Benthic |  | 2 |
| Eupercaria | Labridae | *Coris batuensis* | Invertivore | Benthic | Lagoons, seaward reefs | 2 |
| Eupercaria | Labridae | *Halichoeres leucurus* | Invertivore | Benthic | Silty coastal reefs | 2 |
| Eupercaria | Labridae | *Halichoeres leucurus* | Invertivore | Benthic | Silty coastal reefs | 2 |
| Perciformes | Pomacentridae | *Neopomacentrus sororius* | Herbivore | Benthic | Rocky outcroppings, sand and silt areas | 2 |
| Perciformes | Mullidae | *Parupeneus cyclostomus* | Invertivore | Benthic | Lagoons, seaward reefs | 2 |
| Perciformes | Nemipteridae | *Pentapodus aureofasciatus* | Invertivore, piscivore |  | Mid-water over sand near reefs | 2 |
| Perciformes | Nemipteridae | *Pentapodus setosus* | Invertivore |  | Coastal reefs | 2 |
| Perciformes | Pomacentridae | *Plectroglyphiododon luteobrunneus* | Herbivore | Benthic |  | 2 |
| Perciformes | Pomacentridae | *Pomacentrus aurifrons* | Invertivore | Benthic | Coastal reefs | 2 |
| Perciformes | Labridae | *Pseudochelinus octotaenia* | Invertivore | Benthic |  | 2 |
| Eupercaria | Scaridae | *Scarus flavipectoralis* | Herbivore | Benthic | Coastal, lagoon, outer reefs | 2 |
| Eupercaria | Scaridae | *Scarus flavipectoralis* | Herbivore | Benthic | Coastal, lagoon, outer reefs | 2 |
| Eupercaria | Scaridae | *Scarus russelii* | Herbivore | Benthic | Outer reef slopes | 2 |
| Eupercaria | Scaridae | *Scarus russelii* | Herbivore | Benthic | Outer reef slopes | 2 |
| Perciformes | Nemipteridae | *Scolopsis ciliata* | Invertivore, piscivore | Benthic | Sandy bottoms of lagoons, coastal reefs | 2 |
| Perciformes | Siganidae | *Siganus doliatus* | Herbivore | Benthic |  | 2 |
| Perciformes | Siganidae | *Siganus puelloides* | Herbivore, invertivore | Benthic |  | 2 |
| Perciformes | Balistidae | *Sufflamen bursa* | Invertivore | Benthic |  | 2 |
| Eupercaria | Labridae | *Thalassoma jansenii* | Invertivore |  | Coastal, lagoon, outer reefs | 2 |
|  |  | *Zebrasoma rostratum* |  |  |  | 2 |
| Perciformes | Acanthuridae | *Acanthurus leucocheilus* | Herbivore |  |  | 1 |
| Perciformes | Acanthuridae | *Acanthurus nigrofuscus* | Herbivore | Bentho-pelagic | Lagoons, seaward reefs | 1 |
| Perciformes | Acanthuridae | *Acanthurus tennetii* | Herbivore |  |  | 1 |
| Perciformes | Pomacentridae | *Azurina cyanea* | Invertivore | Bentho-pelagic |  | 1 |
| Eupercaria | Labridae | *Bodianus mesothorax* | Invertivore | Benthic | Outer reef slopes, passes | 1 |
| Perciformes | Pomacanthidae | *Centropyge eibli* | Herbivore, invertivore | Benthic |  | 1 |
| Pecriformes | Serranidae | *Cephalopholis argus* | Invertivore, piscivore | Benthic | Tide pools, outer reefs | 1 |
| Perciformes | Chaetodontidae | *Chaetodon andamanensis* | Invertivore | Benthic |  | 1 |
| Perciformes | Chaetodontidae | *Chaetodon lunulatus* | Invertivore | Benthic |  | 1 |
| Perciformes | Chaetodontidae | *Chaetodon octofasciatus* | Invertivore | Benthic |  | 1 |
| Perciformes | Pomacanthidae | *Chaetodontoplus poliourus* | Herbivore, invertivore | Benthic |  | 1 |
| Eupercaria | Scaridae | *Chlorurus bowersi* | Herbivore | Benthic | Coral rich lagoons, upper edge of channels | 1 |
| Eupercaria | Scaridae | *Chlorurus bowersi* | Herbivore | Benthic | Coral rich lagoons, upper edge of channels | 1 |
| Eupercaria | Scaridae | *Chlorurus capisstratoides* | Herbivore | Benthic | Coral reefs, rubble | 1 |
| Perciformes | Pomacentridae | *Chrysiptera unimaculata* | Herbivore | Benthic | Reef flats | 1 |
| Eupercaria | Labridae | *Coris centralis* | Invertivore | Benthic | Outer reefs | 1 |
| Eupercaria | Labridae | *Coris centralis* | Invertivore | Benthic | Outer reefs | 1 |
| Eupercaria | Labridae | *Cymolutes praetextatus* | Invertivore | Benthic | Weedy aread near reefs, lagoons | 1 |
| Perciformes | Pomacentridae | *Dischistodus perspicillatus* | Herbivore | Benthic | Seagrass | 1 |
| Pecriformes | Serranidae | *Epinephelus fasciatus* | Invertivore, piscivore | Benthic | Coastal, lagoons, seaward reefs | 1 |
| Eupercaria | Labridae | *Halichoeres chloropterus* | Invertivore | Benthic | Silty reefs of lagoons, sheltered coasts | 1 |
| Eupercaria | Labridae | *Halichoeres chloropterus* | Invertivore | Benthic | Silty reefs of lagoons, sheltered coasts | 1 |
| Eupercaria | Labridae | *Halichoeres scapularis* | Invertivore | Benthic | Sand, rubble bottoms, seagrass beds near reefs | 1 |
| Eupercaria | Labridae | *Hemigymnus fasciatus* | Invertivore | Benthic | Lagoons, passes, outer slopes | 1 |
| Eupercaria | Labridae | *Hemigymnus fasciatus* | Invertivore | Benthic | Lagoons, passes, outer slopes | 1 |
| Eupercaria | Labridae | *Hemigymnus melapterus* | Invertivore | Benthic | Sand, rubble, coral areas | 1 |
| Eupercaria | Labridae | *Labroides dimidiatus* | Invertivore | Benthic | Coral reefs | 1 |
| Perciformes | Malacanthidae | *Malacanthus latovittatus* | Invertivore | Benthic |  | 1 |
| Perciformes | Balistidae | *Melichthys indicus* | Invertivore | Benthic |  | 1 |
| Perciformes | Acanthuridae | *Naso elegans* | Herbivore | Benthic |  | 1 |
| Eupercaria | Labridae | *Oxycheilinus orientalis* | Invertivore, piscivore |  | Lagoons, reefs | 1 |
| Perciformes | Mullidae | *Parupeneus barberinus* | Invertivore | Benthic | Lagoon, seaward reefs | 1 |
| Pecriformes | Serranidae | *Plectropomus leopardus* | Invertivore, piscivore | Benthic | Coastal, lagoon reefs | 1 |
| Perciformes | Haemulidae | *Plectrorhinchus vittatus* | Piscivore, invertivore | Bentho-pelagic |  | 1 |
| Perciformes | Pomacanthidae | *Pomacanthus sexstriatus* | Herbivore, invertivore | Benthic |  | 1 |
| Perciformes | Pomacentridae | *Pomacentrus alleni* | Invertivore | Benthic | Coral and rocky reefs | 1 |
| Perciformes | Pomacentridae | *Pomacentrus alleni* | Invertivore | Benthic | Rubble and dead reef areas | 1 |
| Perciformes | Pomacentridae | *Pomacentrus armillatus* | Herbivore | Benthic | Silty reefs | 1 |
| Perciformes | Pomacanthidae | *Pomocanthus xanthometopon* | Herbivore, invertivore | Benthic |  | 1 |
| Perciformes | Rachycentridae | *Rachycentron canadum* | Piscivore, invertivore | Bentho-pelagic |  | 1 |
| Eupercaria | Scaridae | *Scarus altipinnis* | Herbivore | Benthic | Shallow reefs, outer slopes | 1 |
| Eupercaria | Scaridae | *Scarus altipinnis* | Herbivore | Benthic | Shallow reefs, outer slopes | 1 |
| Eupercaria | Scaridae | *Scarus niger* | Herbivore | Benthic | Coastal, lagoon, outer reefs | 1 |
| Eupercaria | Scaridae | *Scarus niger* | Herbivore | Benthic | Coastal, lagoon, outer reefs | 1 |
| Eupercaria | Scaridae | *Scarus viridifucatus* | Herbivore | Benthic | Coastal, lagoon, outer reefs | 1 |
| Eupercaria | Scaridae | *Scarus viridifucatus* | Herbivore | Benthic | Coastal, lagoon, outer reefs | 1 |
| Perciformes | Siganidae | *Siganus guttatus* | Herbivore, invertivore | Benthic |  | 1 |
| Perciformes | Siganidae | *Siganus sutor* | Herbivore, invertivore | Benthic |  | 1 |
| Perciformes | Balistidae | *Sufflamen albicaudatum* | Invertivore | Benthic |  | 1 |
| Perciformes | Balistidae | *Sufflamen fraenatum* | Invertivore | Benthic |  | 1 |
| Perciformes | Synodontinae | *Synodus dermatogenys* | Piscivore | Benthic |  | 1 |

**Table 2:** Summary of traits of fish encountered on baited remote underwater video stations (BRUVS).

| **Order** | **Family** | **Species** | **Guild** | **Column** | **Habitat** | **Abundance** |
| --- | --- | --- | --- | --- | --- | --- |
| *Perciformes* | Lutjanidae | *Lutjanus decussatus* | Invertivore, Piscivore | Benthic | Coastal reefs, lagoons and outer slopes | 143 |
| *Pecriformes* | Serranidae | *Cephalopholis argus* | Invertivore, piscivore | Benthic | Tide pools, outer reefs | 97 |
| *Perciformes* | Balistidae | *Balistapus undulatus* | Invertivore | Bentho-pelagic |  | 61 |
| *Carangiformes* | Carangidae | *Caranx melampygus* | Piscivore | Pelagic | Outer reefs | 43 |
| *Perciformes* | Nemipteridae | *Scolopsis bilineatus* | Invertivore, piscivore | Benthic | Sand and rubble fringe in reefs | 41 |
| *Perciformes* | Lutjanidae | *Lutjanus bohar* | Invertivore, Piscivore | Benthic | Lagoons, outer reefs | 39 |
| *Perciformes* | Lethrinidae | *Monotaxis grandoculis* | Invertivore | Bentho-pelagic | Coastal reefs, lagoons, outer slopes | 36 |
| *Perciformes* | Lutjanidae | *Lutjanus gibbus* | Invertivore, Piscivore | Benthic | Lagoons, passages and outer reef slopes | 30 |
| *Perciformes* | Lutjanidae | *Lutjanus kasmira* | Invertivore, Piscivore | Bentho-pelagic | Shallow lagoons and outer reef slopes | 30 |
| *Perciformes* | Balistidae | *Sufflamen chrysopterum* | Invertivore | Benthic |  | 29 |
| *Perciformes* | Lethrinidae | *Lethrinus harak* | Invertivore, piscivore | Benthic | Sandy shallows next to shore, coastal reefs, lagoons | 24 |
| *Perciformes* | Nemipteridae | *Scolopsis ciliatus* | Invertivore, piscivore | Benthic | Sandy fringe of coastal reefs, lagoon reefs | 23 |
| *Carangiformes* | Carangidae | *Carangoides bajad* | Invertivore, piscivore | Benthic | Coastal reefs, outer slopes | 20 |
| *Perciformes* | Balistidae | *Melichthys indicus* | Invertivore | Benthic |  | 20 |
| *Perciformes* | Nemipteridae | *Scolopsis aurata* | Invertivore, piscivore | Benthic | Sandy bottoms of lagoons, coastal reefs | 19 |
| *Pecriformes* | Serranidae | *Aethaloperca rogaa* | Invertivore, piscivore | Benthic | Caves near seaward reefs | 16 |
| *Pecriformes* | Serranidae | *Epinephelus fasciatus* | Invertivore, piscivore | Benthic | Coastal, lagoons, seaward reefs | 14 |
| *Pecriformes* | Serranidae | *Plectropomus areolatus* | Piscivore | Benthic | Coastal reefs, lagoons, outer reefs | 11 |
| *Pecriformes* | Serranidae | *Cephalopholis miniata* | Invertivore, piscivore | Benthic | Coastal, lagoons, seaward reefs | 10 |
| *Perciformes* | Lethrinidae | *Lethrinus olivaceus* | Invertivore, piscivore | Benthic | Sand bottoms of lagoons and outer slopes | 10 |
| *Tetraodontiformes* | Balistidae | *Odonus niger* | Invertivore | Benthic |  | 7 |
| *Pecriformes* | Serranidae | *Variola louti* | Invertivore, piscivore | Benthic | Lagoons, outer reefs | 7 |
| *Perciformes* | Lethrinidae | *Gymnocranius grandoculis* | Invertivore, piscivore | Benthic | Sand or rubble bottoms | 6 |
| *Pecriformes* | Serranidae | *Plectropomus maculatus* | Invertivore, piscivore | Pelagic | Silty coastal reefs | 6 |
| *Tetraodontiformes* | Balistidae | *Balistoides viridescens* | Invertivore | Benthic |  | 5 |
| *Sygnathiformes* | Fistulariidae | *Fistularia commersonii* | Piscivore, Invertivore | Benthic |  | 5 |
| *Perciformes* | Nemipteridae | *Scolopsis affinis* | Invertivore, piscivore | Benthic | Sandy bottoms of lagoons, coastal reefs | 5 |
| *Carangiformes* | Carangidae | *Atule mate* | Piscivore, Invertivore | Pelagic |  | 4 |
| *Pecriformes* | Serranidae | *Epinephelus malabaricus* | Invertivore, piscivore | Benthic | Reefs, lagoons, estuaries | 4 |
| *Pecriformes* | Serranidae | *Epinephelus merra* | Invertivore, piscivore | Benthic | Coastal, lagoon, sheltered outer reefs | 4 |
| *Perciformes* | Lutjanidae | *Macolor niger* | Invertivore, Piscivore | Benthic | Lagoons, passages and outer reef slopes | 4 |
| *Carangiformes* | Carangidae | *Carangoides plagiotaenia* | Invertivore, piscivore | Benthic (?) | Along edge of steep outer reef slopes | 3 |
| *Anguilliformes* | Muraenidae | *Gymnothorax javanicus* | Piscivore, Invertivore | Benthic |  | 3 |
| *Perciformes* | Lethrinidae | *Lethrinus xanthochilus* | Invertivore, piscivore | Benthic | Sand and rubble bottoms near reefs | 3 |
| *Perciformes* | Lutjanidae | *Lutjanus fulviflamma* | Invertivore, Piscivore | Benthic | Estuaries, coastal reefs and outer slops | 3 |
| *Carangiformes* | Carangidae | *Scomberoides lysan* | Invertivore, piscivore | Pelagic | Coastal reefs, lagoons, outer slopes | 3 |
| *Perciformes* | Balistidae | *Sufflamen fraenatum* | Invertivore | Benthic |  | 3 |
| *Pecriformes* | Serranidae | *Cephalopholis boenak* | Invertivore, piscivore | Benthic | Silty reefs in sheltered waters, coastal reefs | 2 |
| *Pecriformes* | Serranidae | *Cephalopholis nigripinnis* | Piscivore | Benthic | Rock, coral reefs | 2 |
| *Pecriformes* | Serranidae | *Cephalopholis sonnerati* | Invertivore, piscivore | Benthic | Lagoon, outer reefs | 2 |
| *Pecriformes* | Serranidae | *Epinephelus caeruleopunctatus* | Invertivore, piscivore | Benthic | Coastal, lagoons, seaward reefs | 2 |
| *Anguilliformes* | Muraenidae | *Gymnothorax flavimarginatus* | Piscivore, Invertivore | Benthic |  | 2 |
| *Perciformes* | Lethrinidae | *Lethrinus erythropterus* | Invertivore, piscivore | Benthic | Coastal reefs, lagoons, outer slopes | 2 |
| *Perciformes* | Lethrinidae | *Lethrinus ornatus* | Invertivore, piscivore | Benthic | Seagrass beds, sand and rubble beds of coastal reefs, lagoons | 2 |
| *Perciformes* | Lutjanidae | *Lutjanus fulvus* | Invertivore, Piscivore | Benthic | Coastal reefs, lagoons and outer slopes | 2 |
| *Perciformes* | Lutjanidae | *Lutjanus lemniscatus* | Invertivore, Piscivore | Benthic | Coastal reefs, lagoons and outer slopes | 2 |
| *Pecriformes* | Serranidae | *Plectropomus leopardus* | Invertivore, piscivore | Benthic | Coastal, lagoon reefs | 2 |
| *Perciformes* | Balistidae | *Sufflamen bursa* | Invertivore | Benthic |  | 2 |
| *Myliobatiformes* | Aetobatidae | *Aetobatus narinari* | Invertivore | Benthic |  | 1 |
| *Myliobatiformes* | Aetobatidae | *Aetobatus ocellatus* | Invertivore | Benthic |  | 1 |
| *Pecriformes* | Serranidae | *Anyperodon leucogrammicus* | Piscivore | Benthic | Sheltered coastal reefs, outer reefs | 1 |
| *Perciformes* | Lutjanidae | *Aprion virescens* | Invertivore, piscivore | Bentho-pelagic | Lagoons, reef passes, outer slopes | 1 |
| *Carangiformes* | Carangidae | *Carangoides fulvoguttatus* | Invertivore, piscivore | Benthic | Coastal reefs, lagoons, outer slopes | 1 |
| *Carangiformes* | Carangidae | *Caranx ignobilis* | Invertivore, piscivore | Pelagic | Seaward reef slopes | 1 |
| *Carcharhiniformes* | Carcharhinidae | *Carcharhinus melanopterus* | Piscivore, Invertivore | Pelagic |  | 1 |
| *Eupercaria* | Labridae | *Cheilinus undulatus* | Invertivore, piscivore | Benthic | Lagoons, outer reefs | 1 |
| *Eupercaria* | Labridae | *Cheilinus undulatus* | Invertivore, piscivore | Benthic | Lagoons, outer reefs | 1 |
| *Pecriformes* | Serranidae | *Epinephelus longispinis* | Invertivore, piscivore | Benthic | Coastal reefs | 1 |
| *Pecriformes* | Serranidae | *Epinephelus macrospilos* | Invertivore, piscivore | Benthic | Coastal, lagoons, outer reefs | 1 |
| *Perciformes* | Lethrinidae | *Gymnocranius microdon* | Invertivore | Benthic | Sand or rubble bottoms near reefs and on outer slopes | 1 |
| *Perciformes* | Lethrinidae | *Lethrinus microdon* | Invertivore, piscivore | Benthic | Coastal reefs, lagoons, outer slopes | 1 |
| *Perciformes* | Lutjanidae | *Lutjanus monostigma* | Invertivore, Piscivore | Benthic | Outer reefs | 1 |
| *Perciformes* | Lutjanidae | *Lutjanus rivulatus* | Invertivore, Piscivore | Benthic | Seaward reefs, inshore flats | 1 |
| *Perciformes* | Lutjanidae | *Lutjanus russelii* | Invertivore, Piscivore | Benthic | Estuaries, coral reefs | 1 |
| *Perciformes* | Malacanthidae | *Malacanthus latovittatus* | Invertivore | Benthic |  | 1 |
| *Eupercaria* | Labridae | *Oxycheilinus digrammus* | Invertivore | Benthic | Lagoons, seaward reefs | 1 |
| *Eupercaria* | Haemulidae | *Plectorhinchus albovittatus* | Unknown | Benthic | Lagoons, seaward reefs | 1 |
| *Eupercaria* | Haemulidae | *Plectorhinchus chaetodonoides* | Invertivore, piscivore | Benthic | Ledges of coastal reefs, lagoons, seaward reefs | 1 |
| *Pecriformes* | Serranidae | *Plectropomus pessuliferus* | Piscivore | Benthic | Coastal, platforms, outer reefs | 1 |
| *Perciformes* | Scorpaenidae | *Pterois volitans* | Piscivore, Invertivore | Benthic |  | 1 |
| *Carangiformes* | Echeneidae | *Remora remora* | Invertivore | Pelagic |  | 1 |
| *Perciformes* | Nemipteridae | *Scolopsis lineata* | Invertivore, piscivore | Benthic | Sand and rubble fringe in reefs | 1 |
| *Myliobatiformes* | Dasyatidae | *Urogymnus granulatus* | Piscivore, Invertivore | Benthic |  | 1 |

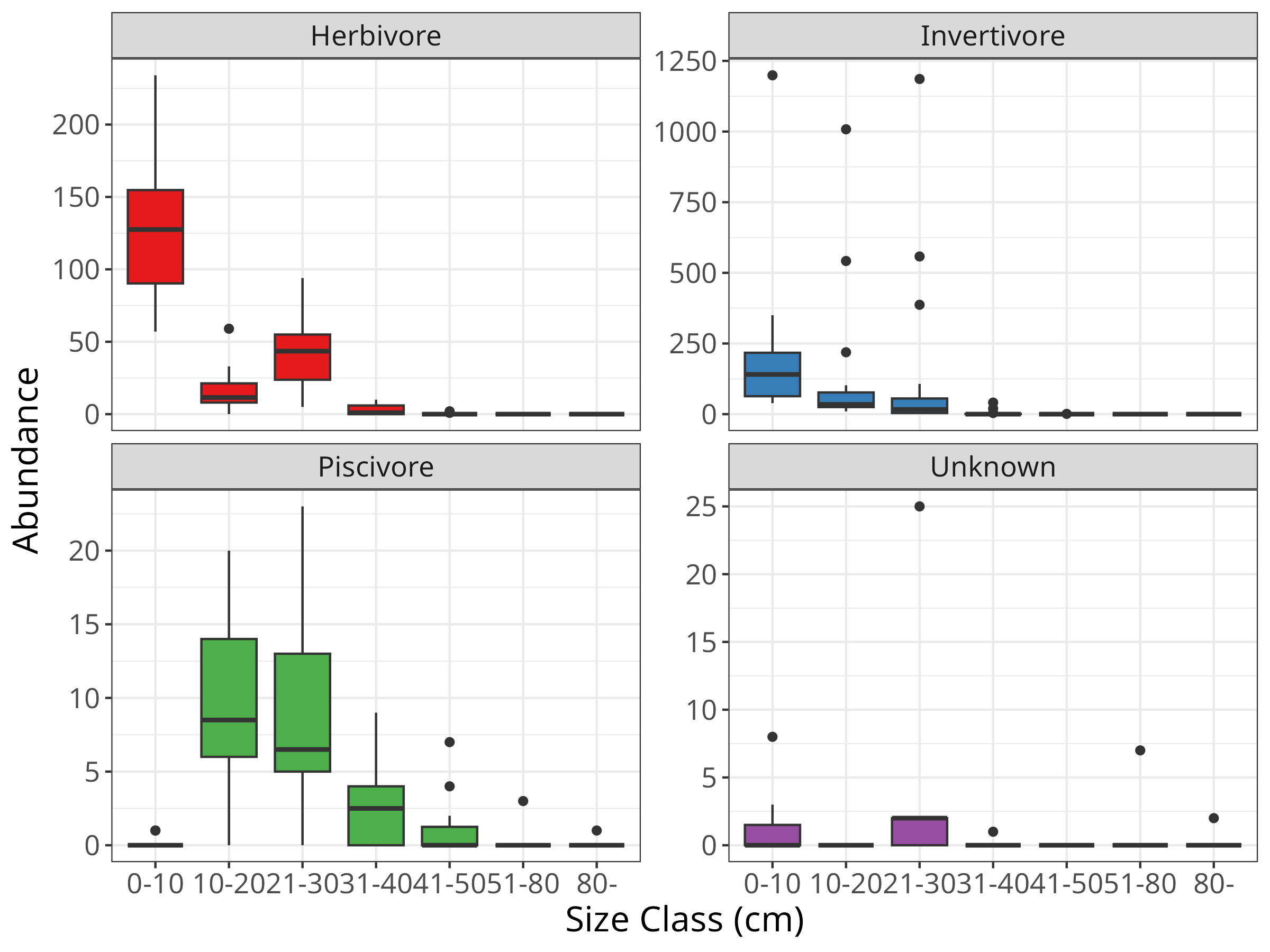

**Figure B1:** Size class distribution of fish encountered on transects.

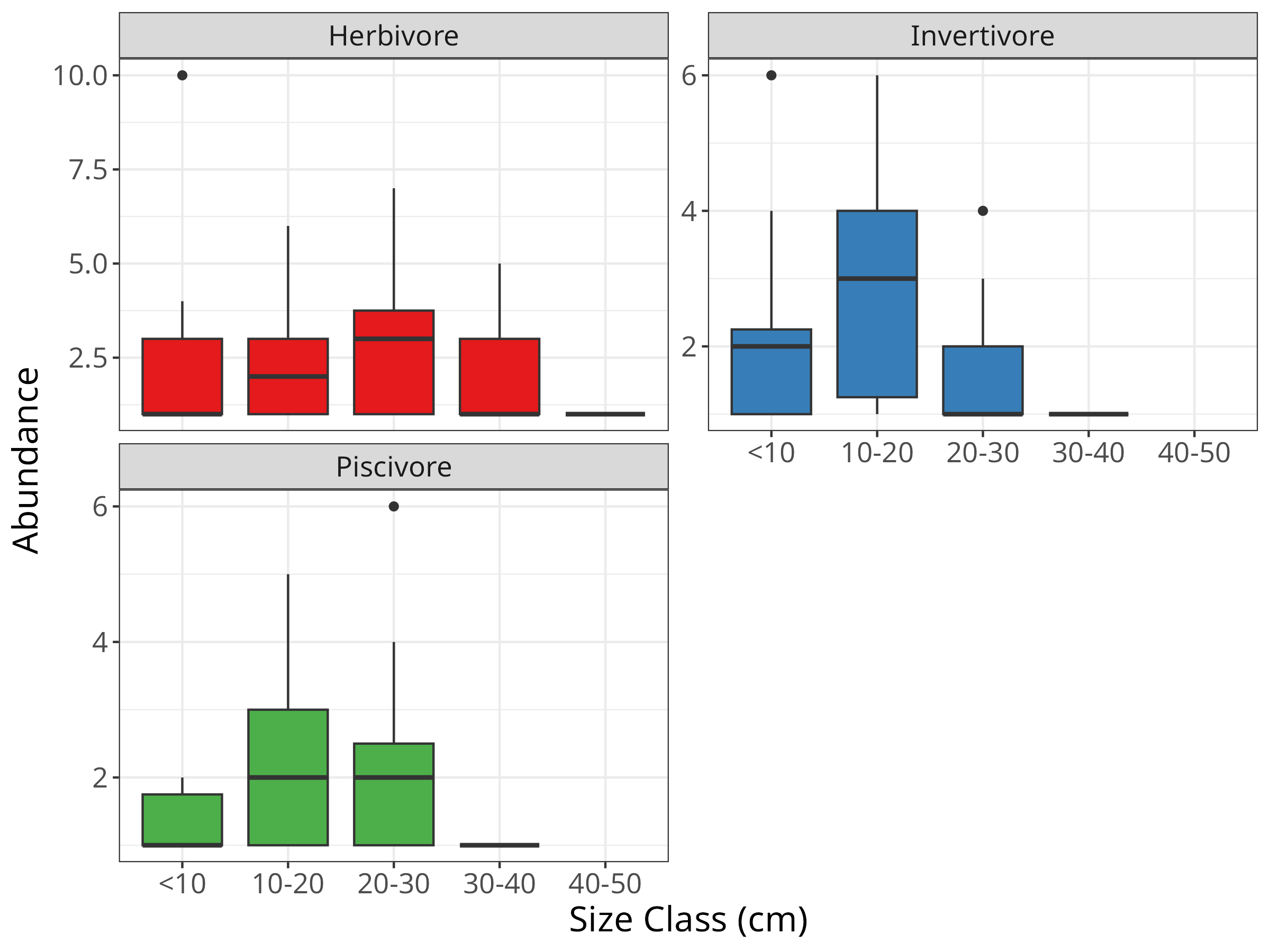

**Figure B2:** Size class distribution of fish encountered during behavioural observations.

### Appendix C: Summary of analysis of effect of protection status on predator assemblages and benthic habitats

**Table 1:** Summary of site level random intercepts modelling the effect of protection on the abundance of predators encountered in BRUVS.

| **Site** | **Mean** | **Median** | **Lower CI** | **Upper CI** |
| --- | --- | --- | --- | --- |
| Aquarium | 0.486 | 0.482 | 0.141 | 0.846 |
| Babu's Bar | -0.166 | -0.167 | -0.523 | 0.192 |
| Boat | -0.138 | -0.133 | -0.502 | 0.219 |
| Deep Deception | 0.323 | 0.317 | 0.005 | 0.665 |
| Hanging Garden | -0.237 | -0.230 | -0.646 | 0.145 |
| Horros's Hangout | 0.029 | 0.025 | -0.353 | 0.418 |
| I-95 | 0.169 | 0.172 | -0.195 | 0.523 |
| Juvi | 0.106 | 0.105 | -0.244 | 0.459 |
| Oshark | -0.495 | -0.490 | -0.934 | -0.096 |
| Pilot | -0.836 | -0.817 | -1.383 | -0.363 |
| Playground | 0.588 | 0.582 | 0.248 | 0.944 |
| Redskin | 0.503 | 0.498 | 0.184 | 0.845 |
| Slope | -0.054 | -0.051 | -0.446 | 0.310 |
| Tarmugli | 0.130 | 0.129 | -0.178 | 0.451 |
| Tarmugli-West | -0.158 | -0.153 | -0.548 | 0.229 |
| Twins | -0.253 | -0.253 | -0.632 | 0.125 |

**Table 2:** Summary of fixed effects modelling the effect of protection on the abundance of predators encountered in BRUVS.

| **Variable** | **Mean** | **Median** | **Lower CI** | **Upper CI** | **P(>0)** |
| --- | --- | --- | --- | --- | --- |
| Intercept | 2.530 | 2.535 | 2.259 | 2.795 | 1.000 |
| Protection | 0.363 | 0.362 | 0.027 | 0.702 | 0.961 |

**Table 3:** Goodness of fit modelling the effect of protection on the abundance of predators encountered in BRUVS.

| **Type** | **Estimate** | **SE** | **Lower CI** | **Upper CI** |
| --- | --- | --- | --- | --- |
| Conditional | 0.409 | 0.056 | 0.292 | 0.510 |
| Marginal | 0.085 | 0.071 | 0.001 | 0.258 |

**Table 4:** Summary of site level random intercepts modelling the effect of protection on the size of predators encountered in BRUVS.

| **Site** | **Mean** | **Median** | **Lower CI** | **Upper CI** |
| --- | --- | --- | --- | --- |
| Aquarium | -1.189 | -1.160 | -2.272 | -0.215 |
| Babu's Bar | 1.561 | 1.544 | 0.601 | 2.598 |
| Boat | 0.188 | 0.179 | -0.748 | 1.161 |
| Deep Deception | 0.664 | 0.641 | -0.276 | 1.702 |
| Hanging Garden | -0.681 | -0.669 | -1.795 | 0.359 |
| Horros's Hangout | 1.516 | 1.496 | 0.521 | 2.571 |
| I-95 | -1.149 | -1.130 | -2.276 | -0.080 |
| Juvi | 0.986 | 0.973 | 0.004 | 1.979 |
| Oshark | -2.345 | -2.306 | -3.579 | -1.211 |
| Pilot | -0.242 | -0.226 | -1.472 | 0.951 |
| Playground | -1.453 | -1.418 | -2.486 | -0.510 |
| Redskin | 0.615 | 0.608 | -0.329 | 1.587 |
| Slope | -2.486 | -2.457 | -3.746 | -1.328 |
| Tarmugli | 0.954 | 0.964 | 0.010 | 1.839 |
| Tarmugli-West | 0.925 | 0.931 | -0.090 | 1.900 |
| Twins | 1.082 | 1.060 | 0.125 | 2.075 |

**Table 5:** Summary of fixed effects modelling the effect of protection on the size of predators encountered in BRUVS.

| **Variable** | **Mean** | **Median** | **Lower CI** | **Upper CI** | **P(>0)** |
| --- | --- | --- | --- | --- | --- |
| Intercept[1] | -4.546 | -4.527 | -5.371 | -3.783 | 0.000 |
| Intercept[2] | -0.834 | -0.828 | -1.468 | -0.239 | 0.016 |
| Intercept[3] | 1.456 | 1.448 | 0.832 | 2.069 | 1.000 |
| Intercept[4] | 1.599 | 1.589 | 0.929 | 2.290 | 1.000 |
| Intercept[5] | 1.289 | 1.271 | 0.544 | 2.096 | 0.997 |
| Intercept[6] | 1.440 | 1.432 | 0.484 | 2.428 | 0.993 |
| Intercept[7] | -0.118 | -0.082 | -1.714 | 1.337 | 0.463 |
| Intercept[8] | 0.293 | 0.296 | -1.327 | 1.852 | 0.621 |
| Protection | -0.145 | -0.147 | -0.743 | 0.464 | 0.350 |
| Aspect | 0.540 | 0.534 | -0.764 | 1.896 | 0.764 |

**Table 6:** Effect (Log odds ration) of protection on the size of predators encountered in BRUVS.

| **Size** | **Median** | **90% Lower CI** | **50% Lower CI** | **50% Upper CI** | **90% Upper CI** | **P** |
| --- | --- | --- | --- | --- | --- | --- |
| 0-10 | 0.212 | -0.669 | -0.146 | 0.565 | 1.072 | 0.650 |
| 10-20 | 0.164 | -0.585 | -0.114 | 0.516 | 1.027 | 0.648 |
| 20-30 | 0.044 | -0.676 | -0.136 | 0.311 | 0.828 | 0.585 |
| 30-40 | -0.030 | -1.245 | -0.299 | 0.194 | 0.882 | 0.444 |
| 40-50 | -0.167 | -1.954 | -0.580 | 0.247 | 1.370 | 0.369 |
| 50-60 | -0.322 | -2.700 | -0.905 | 0.343 | 1.920 | 0.354 |
| 60-70 | -0.374 | -3.286 | -1.054 | 0.403 | 2.323 | 0.353 |
| 70-80 | -0.459 | -3.914 | -1.266 | 0.475 | 2.757 | 0.352 |
| >80 | -0.783 | -4.761 | -1.960 | 0.638 | 3.411 | 0.350 |

**Table 7:** Goodness of fit modelling the effect of protection on the size of predators encountered in BRUVS.

| **Type** | **Estimate** | **SE** | **Lower CI** | **Upper CI** |
| --- | --- | --- | --- | --- |
| Conditional | 0.222 | 0.025 | 0.175 | 0.275 |
| Marginal | 0.042 | 0.051 | 0.001 | 0.193 |

**Table 8:** Summary of transect level random intercepts modelling the effect of protection on the relative cover of various substrate types.

| **Site** | **Transect** | **Response** | **Median** | **Lower CI** | **Upper CI** |
| --- | --- | --- | --- | --- | --- |
| Boat | 1 | Biomass | -0.306 | -0.950 | 0.260 |
|  | 1 | Sponge | 0.018 | -0.422 | 0.454 |
|  | 1 | Substrate | -0.221 | -1.077 | 0.553 |
|  | 2 | Biomass | -0.329 | -0.949 | 0.244 |
|  | 2 | Sponge | 0.085 | -0.309 | 0.556 |
|  | 2 | Substrate | -0.265 | -1.132 | 0.532 |
| Aquarium | 3 | Biomass | -0.207 | -0.635 | 0.230 |
|  | 3 | Sponge | -0.067 | -0.456 | 0.264 |
|  | 3 | Substrate | -0.283 | -0.886 | 0.295 |
|  | 4 | Biomass | 0.081 | -0.340 | 0.551 |
|  | 4 | Sponge | 0.035 | -0.305 | 0.406 |
|  | 4 | Substrate | -0.202 | -0.790 | 0.392 |
| Pilot | 5 | Biomass | 0.192 | -0.242 | 0.657 |
|  | 5 | Sponge | 0.020 | -0.348 | 0.384 |
|  | 5 | Substrate | 0.366 | -0.233 | 0.943 |
|  | 6 | Biomass | -0.461 | -0.937 | -0.018 |
|  | 6 | Sponge | -0.225 | -0.720 | 0.108 |
|  | 6 | Substrate | -0.733 | -1.366 | -0.150 |
| Slope | 7 | Biomass | 0.615 | 0.171 | 1.111 |
|  | 7 | Sponge | 0.126 | -0.202 | 0.542 |
|  | 7 | Substrate | 0.855 | 0.263 | 1.480 |
|  | 8 | Biomass | 0.716 | 0.274 | 1.214 |
|  | 8 | Sponge | 0.218 | -0.115 | 0.714 |
|  | 8 | Substrate | 1.209 | 0.628 | 1.816 |
| Tarmugli | 9 | Biomass | -0.119 | -0.587 | 0.331 |
|  | 9 | Sponge | -0.031 | -0.410 | 0.294 |
|  | 9 | Substrate | -0.310 | -0.966 | 0.249 |
|  | 10 | Biomass | 0.016 | -0.442 | 0.460 |
|  | 10 | Sponge | -0.040 | -0.409 | 0.284 |
|  | 10 | Substrate | 0.407 | -0.205 | 0.992 |
| Babu's Bar | 11 | Biomass | 0.216 | -0.216 | 0.673 |
|  | 11 | Sponge | 0.033 | -0.322 | 0.382 |
|  | 11 | Substrate | 0.256 | -0.336 | 0.844 |
|  | 12 | Biomass | 0.030 | -0.411 | 0.465 |
|  | 12 | Sponge | 0.039 | -0.320 | 0.404 |
|  | 12 | Substrate | 0.057 | -0.545 | 0.642 |
| Deep Deception | 13 | Biomass | -0.580 | -1.056 | -0.124 |
|  | 13 | Sponge | -0.210 | -0.689 | 0.092 |
|  | 13 | Substrate | -0.724 | -1.330 | -0.140 |
|  | 14 | Biomass | -0.140 | -0.601 | 0.301 |
|  | 14 | Sponge | -0.093 | -0.490 | 0.206 |
|  | 14 | Substrate | -0.636 | -1.260 | -0.078 |
| Juvi | 15 | Biomass | 0.200 | -0.235 | 0.661 |
|  | 15 | Sponge | 0.054 | -0.286 | 0.415 |
|  | 15 | Substrate | -0.056 | -0.629 | 0.507 |
|  | 16 | Biomass | -0.242 | -0.699 | 0.187 |
|  | 16 | Sponge | -0.150 | -0.606 | 0.160 |
|  | 16 | Substrate | -0.680 | -1.303 | -0.099 |
| Hanging Garden | 17 | Biomass | -0.231 | -0.666 | 0.228 |
|  | 17 | Sponge | -0.058 | -0.459 | 0.272 |
|  | 17 | Substrate | -0.441 | -1.051 | 0.128 |
|  | 18 | Biomass | 0.234 | -0.218 | 0.685 |
|  | 18 | Sponge | 0.079 | -0.251 | 0.474 |
|  | 18 | Substrate | -0.150 | -0.753 | 0.445 |
| Twins | 19 | Biomass | 0.377 | -0.158 | 0.957 |
|  | 19 | Sponge | -0.022 | -0.489 | 0.412 |
|  | 19 | Substrate | 0.065 | -0.753 | 0.812 |
|  | 20 | Biomass | 0.383 | -0.180 | 0.983 |
|  | 20 | Sponge | -0.002 | -0.446 | 0.424 |
|  | 20 | Substrate | 0.242 | -0.556 | 0.998 |
| Oh Shark | 21 | Biomass | -0.265 | -0.735 | 0.182 |
|  | 21 | Sponge | -0.234 | -0.713 | 0.080 |
|  | 21 | Substrate | -0.641 | -1.309 | -0.031 |
|  | 22 | Biomass | -0.492 | -0.972 | -0.038 |
|  | 22 | Sponge | -0.185 | -0.650 | 0.133 |
|  | 22 | Substrate | -0.819 | -1.459 | -0.209 |
| Playground | 23 | Biomass | 0.384 | -0.059 | 0.853 |
|  | 23 | Sponge | 0.095 | -0.233 | 0.514 |
|  | 23 | Substrate | 0.340 | -0.279 | 0.938 |
|  | 24 | Biomass | 0.480 | 0.044 | 0.958 |
|  | 24 | Sponge | 0.159 | -0.160 | 0.610 |
|  | 24 | Substrate | 0.811 | 0.198 | 1.399 |

**Table 9:** Summary of fixed effects modelling the effect of protection on the relative cover of various substrate types.

| **Response** | **Variable** | **Median** | **Lower CI** | **Upper CI** |
| --- | --- | --- | --- | --- |
| Biomass | Intercept | 2.229 | 1.368 | 3.010 |
|  | Protection | -0.895 | -1.249 | -0.486 |
|  | Aspect | -0.174 | -0.539 | 0.220 |
| Sponge | Intercept | -0.614 | -1.256 | 0.012 |
|  | Protection | -0.712 | -0.991 | -0.417 |
|  | Aspect | 0.030 | -0.265 | 0.321 |
| Substrate | Intercept | 0.388 | -0.712 | 1.453 |
|  | Protection | -0.703 | -1.179 | -0.217 |
|  | Aspect | -0.193 | -0.698 | 0.305 |
|  | Phi | 4.058 | 3.76 | 4.38 |

### Appendix D: Summary of analysis of protection and habitat variables on the abundance of fish encountered during behavioural sampling.

**Table 1:** Summary of site level random intercepts.

| **Site** | **Mean** | **Median** | **Lower CI** | **Upper CI** |
| --- | --- | --- | --- | --- |
| 1 | 0.199 | 0.146 | -0.051 | 0.621 |
| 2 | -0.055 | -0.023 | -0.385 | 0.214 |
| 3 | 0.150 | 0.096 | -0.087 | 0.545 |
| 4 | -0.037 | -0.017 | -0.342 | 0.222 |
| 5 | -0.056 | -0.029 | -0.364 | 0.187 |
| 6 | -0.023 | -0.012 | -0.310 | 0.257 |
| 7 | -0.114 | -0.064 | -0.512 | 0.142 |
| 8 | 0.072 | 0.035 | -0.162 | 0.415 |
| 9 | -0.017 | -0.009 | -0.305 | 0.280 |
| 10 | 0.119 | 0.063 | -0.122 | 0.519 |
| 11 | 0.048 | 0.020 | -0.207 | 0.367 |
| 12 | -0.012 | -0.006 | -0.283 | 0.255 |

**Table 2:** Summary of fixed-effects.

| **Variable** | **Mean** | **Median** | **Lower CI** | **Upper CI** |
| --- | --- | --- | --- | --- |
| Intercept | 1.556 | 1.565 | 1.268 | 1.819 |
| Protected | -0.274 | -0.284 | -0.602 | 0.075 |
| Rugosity | 0.176 | 0.176 | -0.056 | 0.414 |
| Biomass | -0.221 | -0.221 | -0.434 | -0.007 |
| Guild (Invertivore) | -0.103 | -0.101 | -0.351 | 0.149 |
| Rugosity: Guild (Invertivore) | -0.169 | -0.169 | -0.478 | 0.132 |
| Biomass: Guild (Invertivore) | 0.197 | 0.195 | -0.092 | 0.477 |
| Shape | 3.617 | 3.519 | 2.445 | 5.140 |

**Table 3:** Summary of goodness of fit.

| **Type** | **Estimate** | **SE** | **Lower CI** | **Upper CI** |
| --- | --- | --- | --- | --- |
| Conditional | 0.275 | 0.088 | 0.114 | 0.449 |
| Marginal | 0.211 | 0.089 | 0.062 | 0.399 |

**Table 4:** Effect (log response ratio) of habitat complexity (rugosity) on the number of herbivores, invertivores and piscivores encountered during behavioural assays.

| **Foraging Guild** | **Median** | **90% Lower CI** | **50% Lower CI** | **50% Upper CI** | **90% Upper CI** | **P(>0)** |
| --- | --- | --- | --- | --- | --- | --- |
| Herbivore | 0.260 | -0.003 | 0.148 | 0.373 | 0.532 | 0.946 |
| Invertivore | 0.174 | -0.123 | 0.051 | 0.294 | 0.473 | 0.830 |
| Piscivore | 0.108 | -0.264 | -0.051 | 0.257 | 0.472 | 0.678 |

**Table 5:** Effect (log response ratio) of resource availability (algal cover) on the number of herbivores, invertivores and piscivores encountered during behavioural assays.

| **Foraging Guild** | **Median** | **90% Lower CI** | **50% Lower CI** | **50% Upper CI** | **90% Upper CI** | **P(>0)** |
| --- | --- | --- | --- | --- | --- | --- |
| Herbivore | -0.267 | -0.557 | -0.379 | -0.147 | 0.020 | 0.065 |
| Invertivore | -0.058 | -0.389 | -0.194 | 0.072 | 0.261 | 0.378 |
| Piscivore | -0.177 | -0.557 | -0.329 | -0.030 | 0.201 | 0.208 |

### ­Appendix E: Summary of analysis of protection, plot-level predictors and individual-level predictors on the probability of foraging, movement and vigilance.

**Table 1:** Summary of plot-level random intercepts for foraging probability.

| **Site** | **Plot** | **Mean** | **Median** | **Lower CI** | **Upper CI** |
| --- | --- | --- | --- | --- | --- |
| Aquarium | *1* | -0.162 | -0.061 | -1.059 | 0.495 |
|  | *2* | -0.175 | -0.074 | -1.143 | 0.536 |
|  | *3* | 0.078 | 0.020 | -0.755 | 1.052 |
|  | *5* | -0.131 | -0.052 | -1.043 | 0.590 |
| Pilot | *8* | 0.085 | 0.021 | -0.719 | 1.053 |
|  | *9* | 0.078 | 0.020 | -0.737 | 1.021 |
| Slope | *12* | 0.100 | 0.027 | -0.692 | 1.072 |
|  | *14* | 0.040 | 0.012 | -0.734 | 0.878 |
| Boat | *16* | 0.184 | 0.077 | -0.488 | 1.149 |
|  | *18* | 0.099 | 0.035 | -0.717 | 1.067 |
|  | *19* | 0.262 | 0.116 | -0.363 | 1.277 |
|  | *20* | 0.205 | 0.088 | -0.410 | 1.145 |
| Twins | *21* | -0.116 | -0.051 | -1.029 | 0.645 |
|  | *22* | -0.369 | -0.179 | -1.637 | 0.318 |
|  | *23* | -0.177 | -0.063 | -1.185 | 0.578 |
|  | *24* | -0.126 | -0.049 | -0.985 | 0.549 |
|  | *25* | 0.017 | 0.007 | -0.769 | 0.832 |
| Tarmugli | *26* | -0.118 | -0.039 | -1.010 | 0.598 |
|  | *28* | -0.213 | -0.074 | -1.235 | 0.472 |
|  | *30* | 0.158 | 0.061 | -0.572 | 1.098 |
| Babu's Bar | *31* | 0.468 | 0.261 | -0.190 | 1.774 |
|  | *32* | -0.104 | -0.034 | -0.976 | 0.641 |
|  | *33* | 0.062 | 0.016 | -0.722 | 0.954 |
|  | *34* | 0.141 | 0.047 | -0.615 | 1.116 |
|  | *35* | -0.062 | -0.018 | -0.931 | 0.704 |
| Deep Deception | *36* | 0.040 | 0.009 | -0.808 | 0.921 |
|  | *37* | 0.126 | 0.042 | -0.634 | 1.049 |
|  | *38* | -0.040 | -0.007 | -0.958 | 0.830 |
|  | *39* | -0.226 | -0.088 | -1.293 | 0.463 |
|  | *40* | -0.027 | -0.008 | -0.846 | 0.770 |
| Juvi | *41* | -0.031 | -0.008 | -0.919 | 0.778 |
|  | *42* | -0.088 | -0.029 | -0.941 | 0.637 |
|  | *43* | -0.025 | -0.009 | -0.873 | 0.741 |
|  | *44* | 0.095 | 0.036 | -0.724 | 1.037 |
| Oshark | *46* | 0.184 | 0.081 | -0.538 | 1.153 |
|  | *47* | 0.060 | 0.018 | -0.651 | 0.849 |
|  | *48* | 0.172 | 0.071 | -0.488 | 1.099 |
|  | *49* | -0.040 | -0.016 | -0.879 | 0.765 |
|  | *50* | -0.138 | -0.051 | -1.081 | 0.585 |
| Playground | *51* | 0.110 | 0.041 | -0.565 | 0.984 |
|  | *52* | 0.269 | 0.117 | -0.383 | 1.307 |
|  | *53* | -0.136 | -0.057 | -1.069 | 0.580 |
|  | *54* | 0.252 | 0.108 | -0.481 | 1.343 |
|  | *55* | 0.300 | 0.158 | -0.342 | 1.308 |
| Tarmugli-West | *56* | -0.325 | -0.170 | -1.451 | 0.347 |
|  | *57* | -0.331 | -0.149 | -1.522 | 0.350 |
|  | *58* | 0.114 | 0.039 | -0.673 | 1.081 |
|  | *59* | -0.294 | -0.143 | -1.424 | 0.408 |
|  | *60* | -0.187 | -0.070 | -1.171 | 0.502 |

**Table 2:** Summary of species-level random intercepts for foraging probability.

| **Family** | **Species** | **Mean** | **Median** | **Lower CI** | **Upper CI** |
| --- | --- | --- | --- | --- | --- |
| Acanthuridae | *Acanthurus leucocheilus* | 0.601 | 0.527 | -1.675 | 3.167 |
|  | *Acanthurus lineatus* | -0.686 | -0.641 | -2.523 | 1.059 |
|  | *Acanthurus tennetii* | 0.581 | 0.529 | -1.739 | 3.056 |
|  | *Acanthurus tristis* | 0.503 | 0.488 | -0.865 | 1.965 |
|  | *Ctenochaetus striatus* | 2.191 | 2.105 | 0.815 | 3.771 |
|  | *Zebrasoma rostratum* | -1.282 | -1.193 | -3.738 | 0.824 |
| Balistidae | *Balistapus undulatus* | -0.316 | -0.327 | -1.751 | 1.095 |
|  | *Melichthys indicus* | 0.805 | 0.702 | -1.431 | 3.319 |
|  | *Sufflamen albicaudatum* | 0.798 | 0.728 | -1.571 | 3.475 |
|  | *Sufflamen bursa* | 0.923 | 0.831 | -1.305 | 3.411 |
|  | *Sufflamen chrysopterum* | -0.152 | -0.129 | -1.751 | 1.371 |
|  | *Sufflamen fraenatum* | 1.159 | 1.082 | -1.074 | 3.665 |
|  | *Suflamen chrysoterum* | 1.495 | 1.380 | -0.402 | 3.732 |
| Chaetodontidae | *Chaetodon andamanensis* | -1.120 | -1.032 | -3.629 | 1.107 |
|  | *Chaetodon lunulatus* | 0.736 | 0.645 | -1.511 | 3.267 |
|  | *Chaetodon octofasciatus* | 0.929 | 0.863 | -1.259 | 3.378 |
|  | *Chaetodon oxycephalus* | 0.831 | 0.732 | -1.327 | 3.310 |
|  | *Chaetodon trifasciatus* | 1.652 | 1.572 | -0.231 | 3.911 |
| Labridae | *Anampses elegans* | -0.906 | -0.852 | -3.272 | 1.202 |
|  | *Anampses geographicus* | -1.271 | -1.159 | -3.665 | 0.795 |
|  | *Coris batuensis* | 0.032 | 0.029 | -1.991 | 2.065 |
|  | *Coris centralis* | -0.878 | -0.800 | -3.389 | 1.343 |
|  | *Halichoeres bicolor* | -1.026 | -0.992 | -2.537 | 0.397 |
|  | *Halichoeres chierchiae* | 0.669 | 0.660 | -0.724 | 2.125 |
|  | *Halichoeres dussumieri* | 1.461 | 1.423 | -0.005 | 3.028 |
|  | *Halichoeres hortulanus* | -1.053 | -1.025 | -2.546 | 0.325 |
|  | *Halichoeres leucurus* | 1.006 | 0.946 | -1.135 | 3.470 |
|  | *Hemigymnus melapterus* | 0.676 | 0.616 | -1.523 | 3.131 |
|  | *Labroides dimidiatus* | -0.821 | -0.738 | -3.349 | 1.432 |
|  | *Stethojulis trilineata* | 0.840 | 0.782 | -1.357 | 3.246 |
|  | *Thalassoma jansenii* | -1.042 | -0.915 | -3.350 | 0.923 |
|  | *Thalassoma lunare* | 1.273 | 1.176 | -0.713 | 3.592 |
| Lethrinidae | *Monotaxis grandoculis* | -1.185 | -1.043 | -3.575 | 0.850 |
| Malacanthidae | *Malacanthus latovittatus* | -1.123 | -1.042 | -3.527 | 1.002 |
| Mullidae | *Parupeneus cyclostomus* | 0.833 | 0.781 | -1.373 | 3.374 |
|  | *Parupeneus macronemus* | 1.305 | 1.257 | -0.225 | 3.022 |
|  | *Upeneus oligospilus* | 0.931 | 0.857 | -1.276 | 3.402 |
| Pomacanthidae | *Centropyge eibli* | 0.946 | 0.846 | -1.196 | 3.415 |
|  | *Centropyge nox* | 0.837 | 0.745 | -1.337 | 3.325 |
|  | *Pomocanthus xanthometopon* | 0.832 | 0.747 | -1.338 | 3.256 |
| Pomacentridae | *Acanthochromis polyacanthus* | 0.036 | 0.040 | -1.904 | 1.943 |
|  | *Amblygliphidodon silolona* | -1.653 | -1.564 | -3.964 | 0.249 |
|  | *Chromis viridis* | 1.126 | 1.004 | -0.948 | 3.627 |
|  | *Chromis weberi* | -1.143 | -1.065 | -3.631 | 1.020 |
|  | *Chrysiptera talboti* | -2.071 | -1.960 | -4.283 | -0.213 |
|  | *Chrysiptera unimaculata* | 0.799 | 0.706 | -1.349 | 3.237 |
|  | *Dischistodus melanotus* | -2.088 | -1.965 | -4.246 | -0.360 |
|  | *Dischistodus perspicillatus* | -0.880 | -0.813 | -3.300 | 1.256 |
|  | *Neopomacentrus sororius* | -1.765 | -1.680 | -4.216 | 0.328 |
|  | *Plectroglyphiododon luteobrunneus* | -1.340 | -1.253 | -3.784 | 0.650 |
|  | *Pomacentrus albicaudatus* | -0.383 | -0.366 | -1.248 | 0.438 |
|  | *Pomacentrus alleni* | -0.665 | -0.551 | -3.152 | 1.477 |
|  | *Pomacentrus amboinensis* | 0.366 | 0.359 | -1.242 | 2.024 |
|  | *Pomacentrus andamanensis* | -1.917 | -1.799 | -4.095 | -0.142 |
|  | *Pomacentrus aurifrons* | -1.595 | -1.512 | -4.024 | 0.448 |
|  | *Pomacentrus moluccensis* | -1.315 | -1.225 | -3.608 | 0.674 |
|  | *Pomacentrus philippinus* | -0.018 | -0.012 | -1.909 | 1.910 |
|  | *Pomacentrus xanthosternus* | 0.790 | 0.764 | -0.408 | 2.052 |
| Scaridae | *Cetoscarus ocellatus* | 1.124 | 1.047 | -0.869 | 3.512 |
|  | *Chlorurus sordidus* | 0.543 | 0.518 | -0.817 | 2.011 |
|  | *Scarus altipinnis* | 0.666 | 0.592 | -1.508 | 3.257 |
|  | *Scarus flavipectoralis* | -0.405 | -0.409 | -2.354 | 1.572 |
|  | *Scarus niger* | 0.776 | 0.677 | -1.399 | 3.261 |
|  | *Scarus rivulatus* | 1.207 | 1.101 | -0.750 | 3.553 |
|  | *Scarus russelii* | 0.926 | 0.856 | -1.298 | 3.308 |
|  | *Scarus schlegeli* | -0.848 | -0.805 | -2.516 | 0.723 |
|  | *Sparisoma chrysopterum* | -1.954 | -1.857 | -4.210 | -0.081 |
| Siganidae | *Siganus doliatus* | 0.837 | 0.717 | -1.303 | 3.285 |
|  | *Siganus guttatus* | -0.763 | -0.683 | -3.204 | 1.461 |
|  | *Siganus puelloides* | -0.437 | -0.413 | -2.432 | 1.455 |
| Zanclidae | *Zanclus cornutus* | 0.739 | 0.660 | -1.532 | 3.221 |

**Table 3:** Summary of fixed-effects for foraging probability.

| **Variable** | **Mean** | **Median** | **Lower CI** | **Upper CI** | **P(>0)** |
| --- | --- | --- | --- | --- | --- |
| Intercept | 0.532 | 0.532 | -0.357 | 1.376 | 0.846 |
| Protected | 0.036 | 0.029 | -0.565 | 0.646 | 0.532 |
| Rugosity | 0.156 | 0.154 | -0.282 | 0.585 | 0.734 |
| Biomass | 0.461 | 0.464 | -0.055 | 0.981 | 0.931 |
| Predator | -0.584 | -0.588 | -1.179 | 0.022 | 0.056 |
| Group | 0.611 | 0.620 | -0.149 | 1.373 | 0.900 |
| Rugosity:Guild (Invertivore) | 0.285 | 0.288 | -0.318 | 0.899 | 0.785 |
| Biomass:Guild (Invertivore) | 0.111 | 0.108 | -0.504 | 0.733 | 0.617 |
| Protected:Guild (Invertivore) | -0.069 | -0.069 | -0.812 | 0.662 | 0.444 |

**Table 4:** Summary of goodness of fit for foraging probability.

| **Type** | **Estimate** | **SE** | **Lower CI** | **Upper CI** |
| --- | --- | --- | --- | --- |
| Conditional | 0.345 | 0.053 | 0.237 | 0.443 |
| Marginal | 0.089 | 0.027 | 0.039 | 0.146 |

**Table 5:** Summary of plot-level random intercepts for vigilance probability.

| **Site** | **Plot** | **Mean** | **Median** | **Lower CI** | **Upper CI** |
| --- | --- | --- | --- | --- | --- |
| Aquarium | 1 | -0.005 | 0.000 | -0.895 | 0.867 |
|  | 2 | -0.544 | -0.272 | -2.085 | 0.199 |
|  | 3 | -0.064 | -0.009 | -1.140 | 0.867 |
|  | 5 | -0.190 | -0.053 | -1.321 | 0.582 |
| Pilot | 8 | -0.074 | -0.013 | -1.192 | 0.886 |
|  | 9 | -0.045 | -0.012 | -1.059 | 0.935 |
| Slope | 12 | -0.052 | -0.007 | -1.089 | 0.920 |
|  | 14 | 0.265 | 0.100 | -0.491 | 1.459 |
| Boat | 16 | -0.398 | -0.173 | -1.759 | 0.327 |
|  | 18 | -0.068 | -0.007 | -1.110 | 0.847 |
|  | 19 | -0.118 | -0.031 | -1.059 | 0.598 |
|  | 20 | -0.265 | -0.104 | -1.368 | 0.412 |
| Twins | 21 | 0.230 | 0.085 | -0.505 | 1.338 |
|  | 22 | 0.051 | 0.009 | -0.769 | 0.941 |
|  | 23 | 0.281 | 0.114 | -0.457 | 1.503 |
|  | 24 | -0.267 | -0.100 | -1.484 | 0.488 |
|  | 25 | 0.039 | 0.007 | -0.767 | 0.948 |
| Tarmugli | 26 | 0.019 | 0.006 | -0.876 | 0.950 |
|  | 28 | 0.402 | 0.169 | -0.367 | 1.837 |
|  | 30 | -0.139 | -0.044 | -1.211 | 0.674 |
| Babu's Bar | 31 | -0.363 | -0.148 | -1.683 | 0.359 |
|  | 32 | 0.121 | 0.034 | -0.656 | 1.158 |
|  | 33 | -0.041 | -0.012 | -0.995 | 0.838 |
|  | 34 | -0.170 | -0.048 | -1.306 | 0.649 |
|  | 35 | -0.228 | -0.083 | -1.386 | 0.547 |
| Deep Deception | 36 | 0.026 | 0.003 | -0.885 | 1.044 |
|  | 37 | 0.079 | 0.018 | -0.796 | 1.125 |
|  | 38 | 0.100 | 0.023 | -0.816 | 1.232 |
|  | 39 | -0.044 | -0.009 | -1.030 | 0.832 |
|  | 40 | 0.315 | 0.138 | -0.395 | 1.502 |
| Juvi | 41 | -0.130 | -0.023 | -1.242 | 0.754 |
|  | 42 | 0.187 | 0.075 | -0.540 | 1.186 |
|  | 43 | 0.131 | 0.044 | -0.641 | 1.126 |
|  | 44 | -0.231 | -0.082 | -1.418 | 0.569 |
| Oshark | 46 | 0.059 | 0.015 | -0.798 | 1.002 |
|  | 47 | -0.039 | -0.004 | -0.926 | 0.775 |
|  | 48 | -0.063 | -0.011 | -1.001 | 0.708 |
|  | 49 | 0.133 | 0.035 | -0.700 | 1.197 |
|  | 50 | 0.049 | 0.014 | -0.829 | 0.977 |
| Playground | 51 | -0.254 | -0.089 | -1.444 | 0.503 |
|  | 52 | -0.234 | -0.075 | -1.467 | 0.524 |
|  | 53 | -0.230 | -0.067 | -1.454 | 0.567 |
|  | 54 | -0.197 | -0.058 | -1.355 | 0.588 |
|  | 55 | -0.145 | -0.045 | -1.141 | 0.621 |
| Tarmugli-West | 56 | -0.185 | -0.068 | -1.220 | 0.535 |
|  | 57 | -0.289 | -0.108 | -1.493 | 0.468 |
|  | 58 | 0.244 | 0.092 | -0.532 | 1.396 |
|  | 59 | -0.231 | -0.077 | -1.364 | 0.534 |
|  | 60 | 0.033 | 0.009 | -0.859 | 0.961 |

**Table 6:** Summary of species-level random intercepts for vigilance probability.

| **Family** | **Species** | **Mean** | **Median** | **Lower CI** | **Upper CI** |
| --- | --- | --- | --- | --- | --- |
| Acanthuridae | *Acanthurus leucocheilus* | -0.297 | -0.224 | -2.675 | 1.827 |
|  | *Acanthurus lineatus* | -0.729 | -0.609 | -2.987 | 1.125 |
|  | *Acanthurus tennetii* | -0.261 | -0.180 | -2.704 | 2.007 |
|  | *Acanthurus tristis* | -1.258 | -1.075 | -3.519 | 0.462 |
|  | *Ctenochaetus striatus* | -1.552 | -1.400 | -3.666 | 0.038 |
|  | *Zebrasoma rostratum* | 0.470 | 0.446 | -1.435 | 2.415 |
| Balistidae | *Balistapus undulatus* | -0.950 | -0.816 | -3.184 | 0.832 |
|  | *Melichthys indicus* | -0.320 | -0.250 | -2.694 | 1.832 |
|  | *Sufflamen albicaudatum* | -0.441 | -0.376 | -2.932 | 1.752 |
|  | *Sufflamen bursa* | -0.520 | -0.407 | -3.013 | 1.651 |
|  | *Sufflamen chrysopterum* | -0.903 | -0.764 | -3.148 | 0.936 |
|  | *Sufflamen fraenatum* | -0.247 | -0.179 | -2.671 | 1.988 |
|  | *Suflamen chrysoterum* | -0.673 | -0.542 | -2.906 | 1.222 |
| Chaetodontidae | *Chaetodon andamanensis* | -0.305 | -0.244 | -2.722 | 1.888 |
|  | *Chaetodon lunulatus* | -0.386 | -0.309 | -2.732 | 1.683 |
|  | *Chaetodon octofasciatus* | -0.487 | -0.422 | -2.787 | 1.538 |
|  | *Chaetodon oxycephalus* | -0.415 | -0.296 | -2.814 | 1.624 |
|  | *Chaetodon trifasciatus* | -1.023 | -0.924 | -3.076 | 0.741 |
| Labridae | *Anampses elegans* | -0.547 | -0.486 | -2.879 | 1.507 |
|  | *Anampses geographicus* | -0.453 | -0.359 | -2.837 | 1.521 |
|  | *Coris batuensis* | -0.565 | -0.459 | -2.893 | 1.427 |
|  | *Coris centralis* | -0.282 | -0.221 | -2.709 | 1.891 |
|  | *Halichoeres bicolor* | 0.693 | 0.686 | -0.715 | 2.086 |
|  | *Halichoeres chierchiae* | -1.151 | -1.024 | -3.317 | 0.584 |
|  | *Halichoeres dussumieri* | -1.254 | -1.136 | -3.477 | 0.491 |
|  | *Halichoeres hortulanus* | -1.194 | -1.064 | -3.366 | 0.560 |
|  | *Halichoeres leucurus* | -0.329 | -0.252 | -2.738 | 1.801 |
|  | *Hemigymnus melapterus* | -0.293 | -0.238 | -2.738 | 1.908 |
|  | *Labroides dimidiatus* | -0.315 | -0.204 | -2.768 | 1.872 |
|  | *Stethojulis trilineata* | -0.335 | -0.249 | -2.644 | 1.773 |
|  | *Thalassoma jansenii* | -0.567 | -0.464 | -2.911 | 1.444 |
|  | *Thalassoma lunare* | -0.652 | -0.564 | -2.939 | 1.286 |
| Lethrinidae | *Monotaxis grandoculis* | 0.796 | 0.761 | -0.995 | 2.780 |
| Malacanthidae | *Malacanthus latovittatus* | 1.438 | 1.369 | -0.633 | 3.855 |
| Mullidae | *Parupeneus cyclostomus* | -0.408 | -0.335 | -2.762 | 1.754 |
|  | *Parupeneus macronemus* | -1.035 | -0.892 | -3.230 | 0.724 |
|  | *Upeneus oligospilus* | -0.398 | -0.316 | -2.731 | 1.700 |
| Pomacanthidae | *Centropyge eibli* | -0.524 | -0.468 | -2.796 | 1.418 |
|  | *Centropyge nox* | -0.458 | -0.362 | -2.906 | 1.633 |
|  | *Pomocanthus xanthometopon* | -0.362 | -0.267 | -2.723 | 1.686 |
| Pomacentridae | *Acanthochromis polyacanthus* | 0.597 | 0.562 | -1.232 | 2.561 |
|  | *Amblygliphidodon silolona* | 1.982 | 1.887 | 0.131 | 4.124 |
|  | *Chromis viridis* | -0.603 | -0.505 | -2.853 | 1.312 |
|  | *Chromis weberi* | -0.268 | -0.203 | -2.656 | 1.839 |
|  | *Chrysiptera talboti* | 2.215 | 2.137 | 0.568 | 4.011 |
|  | *Chrysiptera unimaculata* | -0.418 | -0.351 | -2.772 | 1.647 |
|  | *Dischistodus melanotus* | 0.806 | 0.782 | -0.552 | 2.273 |
|  | *Dischistodus perspicillatus* | -0.515 | -0.444 | -2.850 | 1.523 |
|  | *Neopomacentrus sororius* | 0.572 | 0.537 | -1.318 | 2.567 |
|  | *Plectroglyphiododon luteobrunneus* | -0.839 | -0.728 | -3.205 | 1.141 |
|  | *Pomacentrus albicaudatus* | 1.057 | 1.043 | 0.175 | 1.977 |
|  | *Pomacentrus alleni* | -0.390 | -0.279 | -2.909 | 1.752 |
|  | *Pomacentrus amboinensis* | 0.256 | 0.255 | -1.495 | 1.988 |
|  | *Pomacentrus andamanensis* | 2.433 | 2.336 | 0.630 | 4.640 |
|  | *Pomacentrus aurifrons* | 1.531 | 1.388 | -0.356 | 3.834 |
|  | *Pomacentrus moluccensis* | 1.777 | 1.678 | -0.142 | 4.082 |
|  | *Pomacentrus philippinus* | -0.880 | -0.763 | -3.101 | 0.993 |
|  | *Pomacentrus xanthosternus* | -0.030 | -0.029 | -1.393 | 1.306 |
| Scaridae | *Cetoscarus ocellatus* | -0.553 | -0.452 | -2.793 | 1.451 |
|  | *Chlorurus sordidus* | -1.141 | -1.012 | -3.307 | 0.557 |
|  | *Scarus altipinnis* | -0.311 | -0.234 | -2.766 | 1.900 |
|  | *Scarus flavipectoralis* | -0.492 | -0.385 | -2.917 | 1.591 |
|  | *Scarus niger* | -0.386 | -0.308 | -2.766 | 1.705 |
|  | *Scarus rivulatus* | -0.610 | -0.476 | -2.937 | 1.410 |
|  | *Scarus russelii* | -0.270 | -0.190 | -2.751 | 1.919 |
|  | *Scarus schlegeli* | -0.105 | -0.062 | -1.806 | 1.408 |
|  | *Sparisoma chrysopterum* | -0.661 | -0.542 | -2.989 | 1.289 |
| Siganidae | *Siganus doliatus* | -0.432 | -0.381 | -2.876 | 1.659 |
|  | *Siganus guttatus* | -0.398 | -0.317 | -2.804 | 1.753 |
|  | *Siganus puelloides* | -0.497 | -0.384 | -2.844 | 1.526 |
| Zanclidae | *Zanclus cornutus* | -0.347 | -0.270 | -2.788 | 1.881 |

**Table 7:** Summary of fixed-effects for vigilance probability.

| **Variable** | **Mean** | **Median** | **Lower CI** | **Upper CI** | **P(>0)** |
| --- | --- | --- | --- | --- | --- |
| Intercept | -0.374 | -0.380 | -1.071 | 0.370 | 0.203 |
| Protected | 0.557 | 0.571 | -0.042 | 1.093 | 0.938 |
| Rugosity | -0.013 | -0.009 | -0.366 | 0.354 | 0.482 |
| Biomass | 0.038 | 0.038 | -0.413 | 0.493 | 0.555 |
| Predator | 0.432 | 0.432 | -0.140 | 0.992 | 0.894 |
| Group | -0.668 | -0.660 | -1.464 | 0.046 | 0.064 |
| Rugosity:Guild (Invertivore) | 0.335 | 0.330 | -0.202 | 0.902 | 0.827 |
| Biomass:Guild (Invertivore) | -0.082 | -0.079 | -0.690 | 0.504 | 0.422 |
| Protected:Guild (Invertivore) | 0.227 | 0.244 | -0.536 | 1.000 | 0.687 |

**Table 8:** Summary of goodness of fit for foraging probability.

| **Type** | **Estimate** | **SE** | **Lower CI** | **Upper CI** |
| --- | --- | --- | --- | --- |
| Conditional | 0.365 | 0.071 | 0.216 | 0.493 |
| Marginal | 0.066 | 0.033 | 0.019 | 0.144 |

**Table 9:** Summary of plot-level random intercepts for movement probability.

| **Site** | **Plot** | **Mean** | **Median** | **Lower CI** | **Upper CI** |
| --- | --- | --- | --- | --- | --- |
| Aquarium | 1 | 0.391 | 0.349 | -0.926 | 1.796 |
|  | 2 | 1.477 | 1.423 | 0.078 | 3.083 |
|  | 3 | -0.372 | -0.299 | -2.425 | 1.487 |
|  | 5 | 0.882 | 0.799 | -0.760 | 2.785 |
| Pilot | 8 | -0.369 | -0.283 | -2.418 | 1.518 |
|  | 9 | -0.410 | -0.321 | -2.478 | 1.424 |
| Slope | 12 | -0.484 | -0.403 | -2.531 | 1.297 |
|  | 14 | -1.164 | -1.064 | -3.131 | 0.473 |
| Boat | 16 | 0.236 | 0.233 | -1.274 | 1.778 |
|  | 18 | -0.347 | -0.276 | -2.337 | 1.471 |
|  | 19 | -0.710 | -0.639 | -2.260 | 0.644 |
|  | 20 | -0.604 | -0.573 | -2.052 | 0.677 |
| Twins | 21 | -0.552 | -0.460 | -2.592 | 1.220 |
|  | 22 | 0.913 | 0.853 | -0.589 | 2.623 |
|  | 23 | -0.345 | -0.261 | -2.476 | 1.582 |
|  | 24 | 0.793 | 0.777 | -0.560 | 2.241 |
|  | 25 | -0.487 | -0.398 | -2.516 | 1.290 |
| Tarmugli | 26 | 0.080 | 0.074 | -1.350 | 1.482 |
|  | 28 | -0.866 | -0.750 | -2.888 | 0.786 |
|  | 30 | -0.428 | -0.365 | -2.018 | 1.006 |
| Babu's Bar | 31 | -1.199 | -1.060 | -3.292 | 0.377 |
|  | 32 | -0.384 | -0.276 | -2.463 | 1.436 |
|  | 33 | -0.323 | -0.246 | -2.412 | 1.526 |
|  | 34 | -0.179 | -0.128 | -2.319 | 1.777 |
|  | 35 | 0.844 | 0.787 | -0.734 | 2.640 |
| Deep Deception | 36 | -0.476 | -0.400 | -2.446 | 1.248 |
|  | 37 | -0.868 | -0.773 | -2.841 | 0.782 |
|  | 38 | -0.318 | -0.268 | -2.309 | 1.561 |
|  | 39 | 0.814 | 0.715 | -0.881 | 2.724 |
|  | 40 | -1.191 | -1.067 | -3.158 | 0.373 |
| Juvi | 41 | 0.474 | 0.408 | -1.233 | 2.347 |
|  | 42 | -0.446 | -0.380 | -2.111 | 0.992 |
|  | 43 | -0.824 | -0.689 | -2.863 | 0.761 |
|  | 44 | 0.104 | 0.102 | -1.501 | 1.711 |
| Oshark | 46 | -1.084 | -0.975 | -3.064 | 0.477 |
|  | 47 | -0.394 | -0.346 | -1.761 | 0.825 |
|  | 48 | -0.595 | -0.541 | -2.030 | 0.676 |
|  | 49 | -0.449 | -0.375 | -2.376 | 1.201 |
|  | 50 | 0.065 | 0.076 | -1.581 | 1.665 |
| Playground | 51 | -0.092 | -0.064 | -1.631 | 1.402 |
|  | 52 | -0.561 | -0.510 | -2.218 | 0.871 |
|  | 53 | 0.560 | 0.536 | -0.766 | 1.944 |
|  | 54 | -0.710 | -0.601 | -2.710 | 0.988 |
|  | 55 | -0.942 | -0.855 | -2.544 | 0.409 |
| Tarmugli-West | 56 | 1.550 | 1.480 | 0.096 | 3.252 |
|  | 57 | 1.715 | 1.605 | 0.100 | 3.777 |
|  | 58 | -1.067 | -0.948 | -3.085 | 0.594 |
|  | 59 | 1.267 | 1.191 | -0.217 | 3.045 |
|  | 60 | 0.523 | 0.454 | -0.884 | 2.152 |

**Table 10:** Summary of species-level random intercepts for movement probability.

| **Family** | **Species** | **Mean** | **Median** | **Lower CI** | **Upper CI** |
| --- | --- | --- | --- | --- | --- |
| Acanthuridae | *Acanthurus leucocheilus* | -0.185 | -0.119 | -2.105 | 1.559 |
|  | *Acanthurus lineatus* | 0.868 | 0.801 | -0.553 | 2.567 |
|  | *Acanthurus tennetii* | -0.147 | -0.114 | -1.895 | 1.579 |
|  | *Acanthurus tristis* | 0.265 | 0.235 | -0.984 | 1.565 |
|  | *Ctenochaetus striatus* | -1.351 | -1.279 | -2.852 | -0.090 |
|  | *Zebrasoma rostratum* | 0.059 | 0.037 | -1.537 | 1.724 |
| Balistidae | *Balistapus undulatus* | 0.641 | 0.587 | -0.640 | 2.105 |
|  | *Melichthys indicus* | -0.327 | -0.262 | -2.223 | 1.365 |
|  | *Sufflamen albicaudatum* | -0.183 | -0.132 | -2.069 | 1.569 |
|  | *Sufflamen bursa* | -0.271 | -0.206 | -2.097 | 1.390 |
|  | *Sufflamen chrysopterum* | 0.408 | 0.370 | -0.869 | 1.826 |
|  | *Sufflamen fraenatum* | -0.585 | -0.489 | -2.431 | 0.998 |
|  | *Suflamen chrysoterum* | -0.742 | -0.628 | -2.606 | 0.780 |
| Chaetodontidae | *Chaetodon andamanensis* | 0.657 | 0.530 | -0.955 | 2.613 |
|  | *Chaetodon lunulatus* | -0.314 | -0.254 | -2.075 | 1.374 |
|  | *Chaetodon octofasciatus* | -0.346 | -0.260 | -2.271 | 1.326 |
|  | *Chaetodon oxycephalus* | -0.365 | -0.275 | -2.121 | 1.173 |
|  | *Chaetodon trifasciatus* | -0.676 | -0.565 | -2.525 | 0.847 |
| Labridae | *Anampses elegans* | 0.645 | 0.537 | -0.863 | 2.487 |
|  | *Anampses geographicus* | 0.623 | 0.521 | -0.897 | 2.436 |
|  | *Coris batuensis* | -0.088 | -0.065 | -1.777 | 1.557 |
|  | *Coris centralis* | 0.385 | 0.305 | -1.189 | 2.222 |
|  | *Halichoeres bicolor* | 0.099 | 0.082 | -1.185 | 1.454 |
|  | *Halichoeres chierchiae* | -0.017 | -0.012 | -1.371 | 1.361 |
|  | *Halichoeres dussumieri* | -0.600 | -0.529 | -2.095 | 0.696 |
|  | *Halichoeres hortulanus* | 1.200 | 1.144 | -0.062 | 2.668 |
|  | *Halichoeres leucurus* | -0.636 | -0.522 | -2.560 | 0.977 |
|  | *Hemigymnus melapterus* | -0.309 | -0.249 | -2.166 | 1.404 |
|  | *Labroides dimidiatus* | 0.389 | 0.321 | -1.214 | 2.231 |
|  | *Stethojulis trilineata* | -0.299 | -0.245 | -2.047 | 1.293 |
|  | *Thalassoma jansenii* | 0.504 | 0.413 | -1.021 | 2.323 |
|  | *Thalassoma lunare* | -0.458 | -0.341 | -2.402 | 1.168 |
| Lethrinidae | *Monotaxis grandoculis* | 0.108 | 0.084 | -1.485 | 1.730 |
| Malacanthidae | *Malacanthus latovittatus* | -0.243 | -0.178 | -2.099 | 1.413 |
| Mullidae | *Parupeneus cyclostomus* | -0.256 | -0.168 | -2.359 | 1.507 |
|  | *Parupeneus macronemus* | -0.829 | -0.751 | -2.399 | 0.483 |
|  | *Upeneus oligospilus* | -0.259 | -0.194 | -2.099 | 1.453 |
| Pomacanthidae | *Centropyge eibli* | -0.320 | -0.247 | -2.158 | 1.364 |
|  | *Centropyge nox* | -0.282 | -0.224 | -2.131 | 1.415 |
|  | *Pomocanthus xanthometopon* | -0.266 | -0.205 | -2.137 | 1.483 |
| Pomacentridae | *Acanthochromis polyacanthus* | -0.519 | -0.399 | -2.394 | 0.962 |
|  | *Amblygliphidodon silolona* | -0.587 | -0.486 | -2.446 | 0.950 |
|  | *Chromis viridis* | -0.354 | -0.272 | -2.203 | 1.205 |
|  | *Chromis weberi* | 0.767 | 0.659 | -0.872 | 2.745 |
|  | *Chrysiptera talboti* | -0.526 | -0.450 | -2.094 | 0.821 |
|  | *Chrysiptera unimaculata* | -0.292 | -0.220 | -2.194 | 1.379 |
|  | *Dischistodus melanotus* | 0.349 | 0.315 | -0.972 | 1.692 |
|  | *Dischistodus perspicillatus* | 0.740 | 0.649 | -0.793 | 2.588 |
|  | *Neopomacentrus sororius* | 0.639 | 0.545 | -0.876 | 2.458 |
|  | *Plectroglyphiododon luteobrunneus* | 1.110 | 1.010 | -0.392 | 3.028 |
|  | *Pomacentrus albicaudatus* | -1.145 | -1.095 | -2.344 | -0.085 |
|  | *Pomacentrus alleni* | 0.323 | 0.250 | -1.354 | 2.270 |
|  | *Pomacentrus amboinensis* | 0.058 | 0.073 | -1.338 | 1.376 |
|  | *Pomacentrus andamanensis* | -0.941 | -0.802 | -2.858 | 0.571 |
|  | *Pomacentrus aurifrons* | -0.231 | -0.162 | -2.155 | 1.483 |
|  | *Pomacentrus moluccensis* | -0.621 | -0.516 | -2.520 | 0.895 |
|  | *Pomacentrus philippinus* | 0.612 | 0.543 | -0.978 | 2.440 |
|  | *Pomacentrus xanthosternus* | -1.149 | -1.040 | -2.781 | 0.124 |
| Scaridae | *Cetoscarus ocellatus* | -0.419 | -0.341 | -2.259 | 1.140 |
|  | *Chlorurus sordidus* | -0.173 | -0.111 | -1.508 | 1.031 |
|  | *Scarus altipinnis* | -0.320 | -0.251 | -2.182 | 1.370 |
|  | *Scarus flavipectoralis* | 0.321 | 0.267 | -1.266 | 2.048 |
|  | *Scarus niger* | -0.251 | -0.206 | -2.077 | 1.429 |
|  | *Scarus rivulatus* | -0.539 | -0.430 | -2.386 | 1.016 |
|  | *Scarus russelii* | -0.408 | -0.296 | -2.385 | 1.252 |
|  | *Scarus schlegeli* | 0.446 | 0.399 | -0.842 | 1.855 |
|  | *Sparisoma chrysopterum* | 1.226 | 1.121 | -0.333 | 3.211 |
| Siganidae | *Siganus doliatus* | -0.250 | -0.177 | -2.095 | 1.471 |
|  | *Siganus guttatus* | 0.513 | 0.438 | -1.107 | 2.374 |
|  | *Siganus puelloides* | 0.507 | 0.468 | -1.063 | 2.204 |
| Zanclidae | *Zanclus cornutus* | -0.311 | -0.219 | -2.201 | 1.384 |

**Table 11:** Summary of fixed-effects for movement probability.

| **Variable** | **Mean** | **Median** | **Lower CI** | **Upper CI** | **P(>0)** |
| --- | --- | --- | --- | --- | --- |
| Intercept | -1.657 | -1.654 | -2.117 | -1.193 | 0.000 |
| Protected | 0.198 | 0.200 | -0.227 | 0.605 | 0.780 |
| Rugosity | 0.064 | 0.062 | -0.191 | 0.324 | 0.659 |
| Biomass | 0.140 | 0.141 | -0.186 | 0.458 | 0.753 |
| Predator | 0.270 | 0.267 | -0.145 | 0.690 | 0.862 |
| Group | -0.195 | -0.201 | -0.742 | 0.368 | 0.274 |
| Rugosity:Guild (Invertivore) | 0.186 | 0.191 | -0.238 | 0.592 | 0.778 |
| Biomass:Guild (Invertivore) | -0.146 | -0.135 | -0.642 | 0.339 | 0.315 |
| Protected:Guild (Invertivore) | 0.558 | 0.563 | -0.070 | 1.161 | 0.926 |

**Table 12:** Summary of goodness of fit for movement probability.

| **Type** | **Estimate** | **SE** | **Lower CI** | **Upper CI** |
| --- | --- | --- | --- | --- |
| Conditional | 0.347 | 0.070 | 0.202 | 0.478 |
| Marginal | 0.084 | 0.034 | 0.028 | 0.160 |

### ­Appendix F: Summary of analysis of protection, plot-level predictors and individual-level predictors on the time budget of fish.

**Table 1:** Summary of plot-level random intercepts for foraging time.

| **Site** | **Plot** | **Mean** | **Median** | **Lower CI** | **Upper CI** |
| --- | --- | --- | --- | --- | --- |
| Aquarium | 1 | 0.533 | 0.449 | -0.674 | 1.960 |
|  | 2 | -0.326 | -0.233 | -1.991 | 1.105 |
|  | 5 | 0.422 | 0.316 | -1.049 | 2.164 |
| Pilot | 8 | 0.504 | 0.413 | -0.943 | 2.235 |
| Slope | 14 | -0.067 | -0.028 | -1.751 | 1.499 |
| Boat | 16 | 1.146 | 1.094 | -0.046 | 2.557 |
|  | 19 | -0.235 | -0.151 | -1.915 | 1.292 |
|  | 20 | 0.605 | 0.531 | -0.521 | 1.945 |
| Twins | 21 | -0.343 | -0.298 | -1.613 | 0.861 |
|  | 22 | -0.357 | -0.291 | -1.744 | 0.869 |
|  | 23 | -0.192 | -0.146 | -1.467 | 0.998 |
|  | 24 | 0.193 | 0.164 | -1.204 | 1.653 |
|  | 25 | -0.240 | -0.216 | -1.296 | 0.773 |
| Tarmugli | 26 | -0.403 | -0.311 | -1.948 | 0.882 |
|  | 28 | -0.619 | -0.507 | -2.362 | 0.802 |
|  | 30 | -0.205 | -0.133 | -1.944 | 1.391 |
| Babu's Bar | 31 | 0.436 | 0.318 | -0.951 | 2.227 |
|  | 32 | -0.568 | -0.519 | -1.745 | 0.506 |
|  | 33 | 1.426 | 1.391 | 0.052 | 2.928 |
|  | 34 | 0.048 | 0.015 | -1.239 | 1.382 |
|  | 35 | -0.123 | -0.109 | -1.362 | 1.155 |
| Deep Deception | 36 | 0.581 | 0.470 | -0.670 | 2.169 |
|  | 37 | 0.618 | 0.487 | -0.816 | 2.458 |
|  | 38 | -0.145 | -0.094 | -1.773 | 1.436 |
|  | 39 | -0.253 | -0.207 | -1.585 | 1.027 |
|  | 40 | -0.736 | -0.613 | -2.498 | 0.666 |
| Juvi | 42 | 0.684 | 0.587 | -0.490 | 2.111 |
|  | 43 | -0.546 | -0.432 | -2.273 | 0.834 |
| Oshark | 46 | 0.121 | 0.089 | -0.971 | 1.280 |
|  | 47 | 0.239 | 0.183 | -1.055 | 1.733 |
|  | 48 | 0.074 | 0.049 | -1.201 | 1.411 |
|  | 49 | -0.003 | -0.011 | -1.053 | 1.131 |
|  | 50 | -0.229 | -0.200 | -1.594 | 1.110 |
| Playground | 51 | -0.003 | -0.010 | -1.216 | 1.230 |
|  | 52 | 0.371 | 0.295 | -0.968 | 1.846 |
|  | 53 | 0.045 | 0.013 | -1.043 | 1.245 |
|  | 54 | 1.382 | 1.309 | -0.002 | 2.993 |
|  | 55 | 0.684 | 0.591 | -0.513 | 2.203 |
| Tarmugli-West | 56 | -0.721 | -0.637 | -2.169 | 0.404 |
|  | 57 | -0.998 | -0.871 | -2.725 | 0.326 |
|  | 58 | -0.094 | -0.048 | -1.467 | 1.242 |
|  | 59 | -0.795 | -0.681 | -2.473 | 0.492 |
|  | 60 | -0.583 | -0.487 | -2.232 | 0.724 |

**Table 2:** Summary of species-level random intercepts for foraging time.

| **Family** | **Species** | **Mean** | **Median** | **Lower CI** | **Upper CI** |
| --- | --- | --- | --- | --- | --- |
| Acanthuridae | *Acanthurus tristis* | 1.682 | 1.585 | -0.280 | 3.962 |
|  | *Ctenochaetus striatus* | 2.142 | 2.048 | 0.000 | 4.636 |
|  | *Naso elegans* | 1.499 | 1.420 | -0.584 | 3.895 |
|  | *Zebrasoma rostratum* | -0.788 | -0.749 | -2.780 | 1.120 |
| Balistidae | *Balistapus undulatus* | 0.067 | 0.044 | -1.493 | 1.654 |
|  | *Sufflamen chrysopterum* | -0.966 | -0.791 | -4.020 | 1.537 |
| Chaetodontidae | *Chaetodon trifasciatus* | 1.569 | 1.466 | -0.476 | 3.928 |
| Labridae | *Anampses elegans* | -0.639 | -0.525 | -3.520 | 1.842 |
|  | *Anampses geographicus* | -0.976 | -0.798 | -3.975 | 1.522 |
|  | *Coris batuensis* | -0.713 | -0.548 | -3.721 | 1.827 |
|  | *Coris centralis* | -1.096 | -0.950 | -3.962 | 1.381 |
|  | *Halichoeres bicolor* | -0.533 | -0.537 | -2.005 | 0.923 |
|  | *Halichoeres chierchiae* | 3.017 | 2.814 | -0.162 | 6.985 |
|  | *Halichoeres dussumieri* | 2.401 | 2.381 | 0.658 | 4.188 |
|  | *Halichoeres hortulanus* | 1.151 | 1.100 | -0.617 | 3.217 |
|  | *Halichoeres leucurus* | 0.852 | 0.788 | -1.077 | 2.988 |
|  | *Labroides dimidiatus* | -0.952 | -0.813 | -3.860 | 1.519 |
|  | *Thalassoma jansenii* | -0.778 | -0.614 | -3.780 | 1.823 |
| Mullidae | *Parupeneus macronemus* | 2.019 | 1.964 | 0.234 | 3.947 |
|  | *Upeneus oligospilus* | 2.129 | 2.067 | -0.174 | 4.786 |
| Pomacentridae | *Acanthochromis polyacanthus* | 0.113 | 0.132 | -1.690 | 1.872 |
|  | *Amblygliphidodon silolona* | -1.749 | -1.520 | -4.672 | 0.425 |
|  | *Chrysiptera talboti* | -2.172 | -1.993 | -4.907 | -0.059 |
|  | *Chrysiptera unimaculata* | 1.718 | 1.675 | -0.344 | 3.943 |
|  | *Dischistodus melanotus* | -2.551 | -2.394 | -5.322 | -0.308 |
|  | *Neopomacentrus sororius* | -0.922 | -0.865 | -3.207 | 1.284 |
|  | *Pomacentrus albicaudatus* | 0.245 | 0.254 | -0.657 | 1.136 |
|  | *Pomacentrus alleni* | -0.758 | -0.609 | -3.626 | 1.724 |
|  | *Pomacentrus amboinensis* | -0.541 | -0.521 | -2.428 | 1.322 |
|  | *Pomacentrus andamanensis* | -0.714 | -0.658 | -2.504 | 0.878 |
|  | *Pomacentrus aurifrons* | -1.705 | -1.556 | -4.648 | 0.722 |
|  | *Pomacentrus moluccensis* | -1.196 | -1.141 | -3.171 | 0.567 |
|  | *Pomacentrus philippinus* | -1.298 | -1.137 | -4.293 | 1.213 |
|  | *Pomacentrus xanthosternus* | 0.681 | 0.677 | -0.746 | 2.086 |
| Scaridae | *Cetoscarus ocellatus* | 1.498 | 1.423 | -0.377 | 3.789 |
|  | *Chlorurus capisstratoides* | 2.085 | 2.018 | -0.074 | 4.407 |
|  | *Chlorurus sordidus* | 1.477 | 1.366 | -0.426 | 3.782 |
|  | *Scarus rivulatus* | 0.242 | 0.177 | -1.703 | 2.336 |
|  | *Scarus schlegeli* | -1.422 | -1.230 | -4.309 | 0.813 |
|  | *Sparisoma chrysopterum* | -1.530 | -1.407 | -4.500 | 0.934 |
| Siganidae | *Siganus doliatus* | 1.415 | 1.335 | -0.560 | 3.647 |
|  | *Siganus puelloides* | -0.169 | -0.176 | -2.085 | 1.858 |

**Table 3:** Summary of family-level random intercepts for precision

| **Family** | **Species** | **Mean** | **Median** | **Lower CI** | **Upper CI** |
| --- | --- | --- | --- | --- | --- |
| Acanthuridae | *Acanthurus tristis* | -0.209 | -0.080 | -1.468 | 0.699 |
|  | *Ctenochaetus striatus* | -0.219 | -0.087 | -1.456 | 0.702 |
|  | *Naso elegans* | -0.182 | -0.073 | -1.454 | 0.781 |
|  | *Zebrasoma rostratum* | 0.193 | 0.065 | -0.744 | 1.445 |
| Balistidae | *Balistapus undulatus* | 0.570 | 0.281 | -0.367 | 2.368 |
|  | *Sufflamen chrysopterum* | -0.003 | -0.002 | -1.187 | 1.146 |
| Chaetodontidae | *Chaetodon trifasciatus* | -0.180 | -0.067 | -1.458 | 0.815 |
| Labridae | *Anampses elegans* | 0.032 | 0.004 | -1.219 | 1.311 |
|  | *Anampses geographicus* | -0.001 | 0.001 | -1.264 | 1.224 |
|  | *Coris batuensis* | 0.006 | -0.002 | -1.279 | 1.278 |
|  | *Coris centralis* | 0.004 | 0.003 | -1.198 | 1.217 |
|  | *Halichoeres bicolor* | 0.230 | 0.124 | -0.514 | 1.283 |
|  | *Halichoeres chierchiae* | 0.001 | -0.002 | -1.225 | 1.209 |
|  | *Halichoeres dussumieri* | -0.227 | -0.109 | -1.365 | 0.630 |
|  | *Halichoeres hortulanus* | -0.399 | -0.193 | -1.856 | 0.455 |
|  | *Halichoeres leucurus* | 0.307 | 0.119 | -0.692 | 1.767 |
|  | *Labroides dimidiatus* | -0.010 | 0.000 | -1.235 | 1.207 |
|  | *Thalassoma jansenii* | -0.024 | -0.002 | -1.265 | 1.151 |
| Mullidae | *Parupeneus macronemus* | -0.004 | 0.002 | -0.988 | 0.982 |
|  | *Upeneus oligospilus* | -0.259 | -0.120 | -1.608 | 0.673 |
| Pomacentridae | *Acanthochromis polyacanthus* | 0.233 | 0.088 | -0.749 | 1.561 |
|  | *Amblygliphidodon silolona* | -0.009 | 0.003 | -1.242 | 1.184 |
|  | *Chrysiptera talboti* | 0.009 | 0.001 | -1.231 | 1.299 |
|  | *Chrysiptera unimaculata* | -0.219 | -0.086 | -1.505 | 0.689 |
|  | *Dischistodus melanotus* | 0.006 | -0.003 | -1.191 | 1.281 |
|  | *Neopomacentrus sororius* | 0.151 | 0.044 | -0.809 | 1.383 |
|  | *Pomacentrus albicaudatus* | 0.069 | 0.028 | -0.560 | 0.789 |
|  | *Pomacentrus alleni* | 0.003 | -0.003 | -1.226 | 1.257 |
|  | *Pomacentrus amboinensis* | 0.134 | 0.047 | -0.795 | 1.340 |
|  | *Pomacentrus andamanensis* | 0.314 | 0.117 | -0.645 | 1.830 |
|  | *Pomacentrus aurifrons* | -0.022 | -0.003 | -1.281 | 1.211 |
|  | *Pomacentrus moluccensis* | 0.186 | 0.065 | -0.801 | 1.497 |
|  | *Pomacentrus philippinus* | -0.006 | -0.004 | -1.139 | 1.137 |
|  | *Pomacentrus xanthosternus* | -0.259 | -0.126 | -1.425 | 0.595 |
| Scaridae | *Cetoscarus ocellatus* | -0.192 | -0.064 | -1.488 | 0.798 |
|  | *Chlorurus capisstratoides* | -0.303 | -0.119 | -1.733 | 0.611 |
|  | *Chlorurus sordidus* | -0.206 | -0.080 | -1.502 | 0.754 |
|  | *Scarus rivulatus* | 0.313 | 0.123 | -0.683 | 1.906 |
|  | *Scarus schlegeli* | 0.010 | -0.001 | -1.172 | 1.243 |
|  | *Sparisoma chrysopterum* | -0.003 | -0.001 | -1.229 | 1.181 |
| Siganidae | *Siganus doliatus* | -0.129 | -0.045 | -1.259 | 0.795 |
|  | *Siganus puelloides* | 0.341 | 0.137 | -0.591 | 1.879 |

**Table 4:** Summary of fixed-effects for foraging time.

| **Variable** | **Mean** | **Median** | **Lower CI** | **Upper CI** | **P(>0)** |
| --- | --- | --- | --- | --- | --- |
| Intercept | 0.532 | 0.532 | -0.357 | 1.376 | 0.846 |
| Protected | 0.036 | 0.029 | -0.565 | 0.646 | 0.532 |
| Rugosity | 0.156 | 0.154 | -0.282 | 0.585 | 0.734 |
| Biomass | 0.461 | 0.464 | -0.055 | 0.981 | 0.931 |
| Predator | -0.584 | -0.588 | -1.179 | 0.022 | 0.056 |
| Group | 0.611 | 0.620 | -0.149 | 1.373 | 0.900 |
| Rugosity:Guild (Invertivore) | 0.285 | 0.288 | -0.318 | 0.899 | 0.785 |
| Biomass:Guild (Invertivore) | 0.111 | 0.108 | -0.504 | 0.733 | 0.617 |
| Protected:Guild (Invertivore) | -0.069 | -0.069 | -0.812 | 0.662 | 0.444 |

**Table 5:** Summary of zero and one hyperparameters for foraging time.

| **Variable** | **Mean** | **Median** | **Lower CI** | **Upper CI** | **P(>0)** |
| --- | --- | --- | --- | --- | --- |
| Cut One | 1.938 | 1.937 | 1.613 | 2.276 | 1.000 |
| Cut Zero | 0.094 | 0.104 | -0.563 | 0.741 | 0.598 |

**Table 6:** Summary of goodness of fit for foraging probability.

| **Type** | **Estimate** | **SE** | **Lower CI** | **Upper CI** |
| --- | --- | --- | --- | --- |
| Conditional | 0.619 | 0.077 | 0.442 | 0.738 |
| Marginal | 0.196 | 0.058 | 0.081 | 0.305 |

**Table 7:** Summary of plot-level random intercepts for foraging and vigilance time.

| **Site** | **Plot** | **Mean** | **Median** | **Lower CI** | **Upper CI** |
| --- | --- | --- | --- | --- | --- |
| Aquarium | 1 | 0.133 | 0.108 | -0.954 | 1.239 |
|  | 2 | -1.432 | -1.312 | -3.429 | 0.017 |
|  | 5 | -0.284 | -0.236 | -1.880 | 1.091 |
| Pilot | 8 | -0.381 | -0.244 | -2.053 | 0.976 |
| Slope | 14 | 0.653 | 0.541 | -0.757 | 2.401 |
| Boat | 16 | -0.982 | -0.948 | -2.315 | 0.133 |
|  | 19 | 0.831 | 0.733 | -0.520 | 2.555 |
|  | 20 | -1.003 | -0.941 | -2.339 | 0.135 |
| Twins | 21 | 0.630 | 0.547 | -0.427 | 2.142 |
|  | 22 | 0.091 | 0.090 | -0.981 | 1.190 |
|  | 23 | 0.384 | 0.335 | -0.567 | 1.519 |
|  | 24 | -0.602 | -0.435 | -2.244 | 0.671 |
|  | 25 | 0.280 | 0.262 | -0.550 | 1.174 |
| Tarmugli | 26 | -0.276 | -0.222 | -1.640 | 0.934 |
|  | 28 | 1.384 | 1.282 | -0.086 | 3.222 |
|  | 30 | 0.307 | 0.240 | -1.639 | 2.072 |
| Babu's Bar | 31 | -0.467 | -0.375 | -2.067 | 0.840 |
|  | 32 | 0.372 | 0.318 | -0.529 | 1.374 |
|  | 33 | -0.333 | -0.312 | -1.340 | 0.626 |
|  | 34 | -0.804 | -0.789 | -2.015 | 0.269 |
|  | 35 | -0.206 | -0.169 | -1.295 | 0.827 |
| Deep Deception | 36 | -0.329 | -0.277 | -1.518 | 0.797 |
|  | 37 | -0.188 | -0.117 | -1.793 | 1.185 |
|  | 38 | 0.438 | 0.297 | -0.854 | 2.269 |
|  | 39 | -0.077 | -0.068 | -1.443 | 1.250 |
|  | 40 | 1.286 | 1.146 | -0.092 | 3.152 |
| Juvi | 42 | 0.306 | 0.237 | -0.720 | 1.627 |
|  | 43 | 0.617 | 0.535 | -0.765 | 2.315 |
| Oshark | 46 | 0.485 | 0.454 | -0.488 | 1.538 |
|  | 47 | 0.314 | 0.239 | -1.080 | 1.778 |
|  | 48 | 0.105 | 0.085 | -0.888 | 1.164 |
|  | 49 | -0.040 | -0.021 | -1.064 | 0.931 |
|  | 50 | 0.373 | 0.328 | -0.667 | 1.507 |
| Playground | 51 | -0.257 | -0.211 | -1.590 | 1.036 |
|  | 52 | -0.157 | -0.127 | -1.267 | 1.007 |
|  | 53 | -0.304 | -0.244 | -1.441 | 0.658 |
|  | 54 | -0.376 | -0.277 | -2.054 | 1.087 |
|  | 55 | -0.848 | -0.821 | -2.098 | 0.237 |
| Tarmugli-West | 56 | 0.142 | 0.113 | -0.796 | 1.090 |
|  | 57 | -0.521 | -0.477 | -1.644 | 0.495 |
|  | 58 | 0.898 | 0.784 | -0.274 | 2.452 |
|  | 59 | -0.722 | -0.560 | -2.449 | 0.488 |
|  | 60 | 0.061 | 0.042 | -1.063 | 1.154 |

**Table 8:** Summary of species-level random intercepts for vigilance time.

| **Family** | **Species** | **Mean** | **Median** | **Lower CI** | **Upper CI** |
| --- | --- | --- | --- | --- | --- |
| Acanthuridae | *Acanthurus tristis* | -0.186 | -0.051 | -1.348 | 0.645 |
|  | *Ctenochaetus striatus* | -0.226 | -0.066 | -1.522 | 0.604 |
|  | *Naso elegans* | -0.127 | -0.031 | -1.283 | 0.784 |
|  | *Zebrasoma rostratum* | -0.060 | -0.020 | -0.995 | 0.770 |
| Balistidae | *Balistapus undulatus* | -0.070 | -0.017 | -0.965 | 0.716 |
|  | *Sufflamen chrysopterum* | -0.089 | -0.019 | -1.234 | 0.805 |
| Chaetodontidae | *Chaetodon trifasciatus* | -0.108 | -0.026 | -1.256 | 0.869 |
| Labridae | *Anampses elegans* | -0.052 | -0.015 | -1.053 | 0.882 |
|  | *Anampses geographicus* | -0.122 | -0.031 | -1.246 | 0.766 |
|  | *Coris batuensis* | -0.040 | -0.008 | -1.049 | 0.880 |
|  | *Coris centralis* | -0.055 | -0.009 | -1.189 | 0.927 |
|  | *Halichoeres bicolor* | 0.108 | 0.042 | -0.550 | 0.921 |
|  | *Halichoeres chierchiae* | -0.115 | -0.030 | -1.234 | 0.745 |
|  | *Halichoeres dussumieri* | -0.473 | -0.203 | -2.029 | 0.261 |
|  | *Halichoeres hortulanus* | -0.300 | -0.104 | -1.647 | 0.469 |
|  | *Halichoeres leucurus* | -0.093 | -0.018 | -1.163 | 0.849 |
|  | *Labroides dimidiatus* | -0.092 | -0.012 | -1.240 | 0.826 |
|  | *Thalassoma jansenii* | -0.198 | -0.055 | -1.446 | 0.615 |
| Mullidae | *Parupeneus macronemus* | -0.191 | -0.054 | -1.406 | 0.623 |
|  | *Upeneus oligospilus* | -0.125 | -0.026 | -1.257 | 0.705 |
| Pomacentridae | *Acanthochromis polyacanthus* | 0.111 | 0.028 | -0.666 | 1.117 |
|  | *Amblygliphidodon silolona* | 0.470 | 0.174 | -0.347 | 2.182 |
|  | *Chrysiptera talboti* | 0.688 | 0.358 | -0.150 | 2.568 |
|  | *Chrysiptera unimaculata* | -0.189 | -0.049 | -1.464 | 0.672 |
|  | *Dischistodus melanotus* | 0.104 | 0.029 | -0.642 | 1.080 |
|  | *Neopomacentrus sororius* | 0.151 | 0.048 | -0.571 | 1.176 |
|  | *Pomacentrus albicaudatus* | 0.255 | 0.169 | -0.199 | 0.942 |
|  | *Pomacentrus alleni* | -0.121 | -0.025 | -1.247 | 0.752 |
|  | *Pomacentrus amboinensis* | 0.078 | 0.012 | -0.697 | 1.031 |
|  | *Pomacentrus andamanensis* | 0.426 | 0.205 | -0.252 | 1.780 |
|  | *Pomacentrus aurifrons* | 0.324 | 0.103 | -0.453 | 1.803 |
|  | *Pomacentrus moluccensis* | 0.336 | 0.102 | -0.452 | 1.800 |
|  | *Pomacentrus philippinus* | 0.080 | 0.020 | -0.716 | 1.037 |
|  | *Pomacentrus xanthosternus* | -0.176 | -0.050 | -1.356 | 0.676 |
| Scaridae | *Cetoscarus ocellatus* | 0.043 | 0.006 | -0.804 | 1.017 |
|  | *Chlorurus capisstratoides* | -0.127 | -0.031 | -1.284 | 0.733 |
|  | *Chlorurus sordidus* | -0.135 | -0.031 | -1.312 | 0.792 |
|  | *Scarus rivulatus* | -0.129 | -0.030 | -1.294 | 0.774 |
|  | *Scarus schlegeli* | 0.012 | -0.002 | -0.882 | 0.962 |
|  | *Sparisoma chrysopterum* | 0.170 | 0.062 | -0.552 | 1.147 |
| Siganidae | *Siganus doliatus* | -0.116 | -0.022 | -1.210 | 0.780 |
|  | *Siganus puelloides* | -0.190 | -0.047 | -1.417 | 0.662 |

**Table 9:** Summary of family-level random intercepts for precision of vigilance time.

| **Family** | **Species** | **Mean** | **Median** | **Lower CI** | **Upper CI** |
| --- | --- | --- | --- | --- | --- |
| Acanthuridae | *Acanthurus tristis* | -0.050 | -0.110 | -2.673 | 2.748 |
|  | *Ctenochaetus striatus* | 0.038 | 0.045 | -2.636 | 2.708 |
|  | *Naso elegans* | 0.075 | 0.044 | -2.673 | 2.551 |
|  | *Zebrasoma rostratum* | -0.017 | -0.016 | -1.441 | 1.353 |
| Balistidae | *Balistapus undulatus* | 0.816 | 0.668 | -1.298 | 3.235 |
|  | *Sufflamen chrysopterum* | -0.027 | -0.023 | -2.701 | 2.622 |
| Chaetodontidae | *Chaetodon trifasciatus* | -0.002 | 0.028 | -2.624 | 2.651 |
| Labridae | *Anampses elegans* | -1.230 | -1.124 | -3.212 | 0.388 |
|  | *Anampses geographicus* | -0.074 | -0.046 | -3.658 | 2.949 |
|  | *Coris batuensis* | -1.356 | -1.256 | -3.324 | 0.343 |
|  | *Coris centralis* | 0.079 | 0.066 | -2.615 | 2.764 |
|  | *Halichoeres bicolor* | -1.825 | -1.792 | -3.275 | -0.568 |
|  | *Halichoeres chierchiae* | -0.027 | -0.044 | -2.691 | 2.623 |
|  | *Halichoeres dussumieri* | -0.023 | -0.064 | -2.747 | 2.650 |
|  | *Halichoeres hortulanus* | 0.025 | 0.047 | -2.784 | 2.453 |
|  | *Halichoeres leucurus* | 0.019 | 0.037 | -2.663 | 2.692 |
|  | *Labroides dimidiatus* | -0.013 | 0.047 | -2.738 | 2.572 |
|  | *Thalassoma jansenii* | 0.073 | 0.071 | -2.798 | 2.835 |
| Mullidae | *Parupeneus macronemus* | -0.032 | -0.060 | -2.645 | 2.915 |
|  | *Upeneus oligospilus* | -0.164 | -0.099 | -3.193 | 2.483 |
| Pomacentridae | *Acanthochromis polyacanthus* | 0.136 | 0.092 | -1.871 | 2.180 |
|  | *Amblygliphidodon silolona* | -0.108 | -0.087 | -2.849 | 2.756 |
|  | *Chrysiptera talboti* | -0.011 | 0.097 | -2.978 | 2.520 |
|  | *Chrysiptera unimaculata* | 0.039 | -0.012 | -2.601 | 2.893 |
|  | *Dischistodus melanotus* | -0.042 | -0.018 | -2.892 | 2.921 |
|  | *Neopomacentrus sororius* | 1.108 | 0.764 | -1.137 | 5.367 |
|  | *Pomacentrus albicaudatus* | 0.869 | 0.871 | -0.261 | 1.996 |
|  | *Pomacentrus alleni* | 0.047 | 0.037 | -2.645 | 2.558 |
|  | *Pomacentrus amboinensis* | 0.873 | 0.744 | -1.215 | 3.395 |
|  | *Pomacentrus andamanensis* | 0.879 | 0.774 | -1.146 | 3.323 |
|  | *Pomacentrus aurifrons* | 0.069 | 0.083 | -2.618 | 2.674 |
|  | *Pomacentrus moluccensis* | 0.642 | 0.668 | -1.989 | 3.091 |
|  | *Pomacentrus philippinus* | -0.074 | -0.084 | -2.557 | 2.606 |
|  | *Pomacentrus xanthosternus* | 0.087 | 0.050 | -2.614 | 2.820 |
| Scaridae | *Cetoscarus ocellatus* | -0.010 | -0.060 | -2.720 | 2.605 |
|  | *Chlorurus capisstratoides* | 0.056 | 0.098 | -2.835 | 2.577 |
|  | *Chlorurus sordidus* | -0.169 | -0.059 | -3.722 | 2.709 |
|  | *Scarus rivulatus* | 0.349 | 0.269 | -2.187 | 2.806 |
|  | *Scarus schlegeli* | -0.968 | -0.904 | -2.494 | 0.334 |
|  | *Sparisoma chrysopterum* | -0.004 | 0.033 | -2.827 | 2.725 |
| Siganidae | *Siganus doliatus* | 0.077 | 0.012 | -2.666 | 3.051 |
|  | *Siganus puelloides* | -0.014 | 0.000 | -2.621 | 2.636 |

**Table 10:** Summary of fixed-effects for vigilance time.

| **Variable** | **Mean** | **Median** | **Lower CI** | **Upper CI** | **P(>0)** |
| --- | --- | --- | --- | --- | --- |
| Intercept | -0.374 | -0.380 | -1.071 | 0.370 | 0.203 |
| Protected | 0.557 | 0.571 | -0.042 | 1.093 | 0.938 |
| Rugosity | -0.013 | -0.009 | -0.366 | 0.354 | 0.482 |
| Biomass | 0.038 | 0.038 | -0.413 | 0.493 | 0.555 |
| Predator | 0.432 | 0.432 | -0.140 | 0.992 | 0.894 |
| Group | -0.668 | -0.660 | -1.464 | 0.046 | 0.064 |
| Rugosity:Guild (Invertivore) | 0.335 | 0.330 | -0.202 | 0.902 | 0.827 |
| Biomass:Guild (Invertivore) | -0.082 | -0.079 | -0.690 | 0.504 | 0.422 |
| Protected:Guild (Invertivore) | 0.227 | 0.244 | -0.536 | 1.000 | 0.687 |

**Table 11:** Summary of zero and one hyperparameters for vigilance time.

| **Variable** | **Mean** | **Median** | **Lower CI** | **Upper CI** | **P(>0)** |
| --- | --- | --- | --- | --- | --- |
| Cut One | 1.938 | 1.937 | 1.613 | 2.276 | 1.000 |
| Cut Zero | 0.094 | 0.104 | -0.563 | 0.741 | 0.598 |

**Table 12:** Summary of goodness of fit for vigilance time.

| **Type** | **Estimate** | **SE** | **Lower CI** | **Upper CI** |
| --- | --- | --- | --- | --- |
| Conditional | 0.454 | 0.085 | 0.268 | 0.603 |
| Marginal | 0.203 | 0.066 | 0.079 | 0.325 |

**Table 13**: Summary of plot-level random intercepts for movement time.

| **Site** | **Plot** | **Mean** | **Median** | **Lower CI** | **Upper CI** |
| --- | --- | --- | --- | --- | --- |
| Aquarium | 1 | 0.152 | 0.126 | -1.187 | 1.584 |
|  | 2 | 1.689 | 1.646 | 0.173 | 3.396 |
|  | 5 | -0.378 | -0.295 | -2.495 | 1.563 |
| Pilot | 8 | -0.443 | -0.344 | -2.529 | 1.398 |
| Slope | 14 | -0.702 | -0.615 | -2.699 | 1.142 |
| Boat | 16 | 0.570 | 0.525 | -0.792 | 2.096 |
|  | 19 | -0.611 | -0.502 | -2.600 | 1.098 |
|  | 20 | 0.646 | 0.606 | -0.566 | 1.998 |
| Twins | 21 | -0.584 | -0.458 | -2.544 | 1.100 |
|  | 22 | 0.674 | 0.601 | -0.889 | 2.439 |
|  | 23 | -0.604 | -0.457 | -2.738 | 1.104 |
|  | 24 | 1.442 | 1.399 | -0.135 | 3.249 |
|  | 25 | -0.587 | -0.469 | -2.610 | 1.067 |
| Tarmugli | 26 | 0.841 | 0.803 | -0.542 | 2.377 |
|  | 28 | -0.949 | -0.818 | -2.873 | 0.617 |
|  | 30 | -0.265 | -0.205 | -2.365 | 1.653 |
| Babu's Bar | 31 | -0.540 | -0.432 | -2.551 | 1.195 |
|  | 32 | -0.580 | -0.476 | -2.634 | 1.098 |
|  | 33 | -0.485 | -0.366 | -2.494 | 1.265 |
|  | 34 | -0.245 | -0.173 | -2.370 | 1.667 |
|  | 35 | -0.283 | -0.187 | -2.383 | 1.599 |
| Deep Deception | 36 | -0.342 | -0.268 | -2.395 | 1.448 |
|  | 37 | -0.480 | -0.395 | -2.469 | 1.302 |
|  | 38 | -0.534 | -0.446 | -2.560 | 1.247 |
|  | 39 | 0.609 | 0.539 | -0.852 | 2.238 |
|  | 40 | -0.702 | -0.567 | -2.733 | 0.939 |
| Juvi | 42 | -0.735 | -0.607 | -2.672 | 0.846 |
|  | 43 | -0.417 | -0.312 | -2.558 | 1.520 |
| Oshark | 46 | 0.198 | 0.173 | -1.215 | 1.683 |
|  | 47 | -0.693 | -0.557 | -2.735 | 0.962 |
|  | 48 | -0.356 | -0.286 | -2.302 | 1.406 |
|  | 49 | -0.531 | -0.425 | -2.515 | 1.145 |
|  | 50 | -0.288 | -0.207 | -2.360 | 1.556 |
| Playground | 51 | -0.500 | -0.400 | -2.497 | 1.252 |
|  | 52 | -0.578 | -0.469 | -2.558 | 1.118 |
|  | 53 | 0.344 | 0.310 | -1.055 | 1.862 |
|  | 54 | -0.362 | -0.257 | -2.414 | 1.471 |
|  | 55 | -0.558 | -0.453 | -2.569 | 1.167 |
| Tarmugli-West | 56 | 1.517 | 1.482 | 0.234 | 2.894 |
|  | 57 | 1.698 | 1.657 | 0.192 | 3.285 |
|  | 58 | 0.246 | 0.195 | -1.159 | 1.783 |
|  | 59 | 1.910 | 1.869 | 0.286 | 3.635 |
|  | 60 | 0.424 | 0.346 | -0.850 | 1.928 |

**Table 14:** Summary of species-level random intercepts for movement time.

| **Family** | **Species** | **Mean** | **Median** | **Lower CI** | **Upper CI** |
| --- | --- | --- | --- | --- | --- |
| Acanthuridae | *Acanthurus tristis* | -0.120 | -0.024 | -1.408 | 0.913 |
|  | *Ctenochaetus striatus* | -0.320 | -0.113 | -1.809 | 0.555 |
|  | *Naso elegans* | -0.128 | -0.025 | -1.441 | 0.938 |
|  | *Zebrasoma rostratum* | 0.318 | 0.120 | -0.589 | 1.707 |
| Balistidae | *Balistapus undulatus* | 0.079 | 0.019 | -0.746 | 1.027 |
|  | *Sufflamen chrysopterum* | 0.145 | 0.044 | -0.751 | 1.290 |
| Chaetodontidae | *Chaetodon trifasciatus* | -0.152 | -0.035 | -1.582 | 1.027 |
| Labridae | *Anampses elegans* | 0.009 | 0.000 | -1.059 | 1.032 |
|  | *Anampses geographicus* | 0.518 | 0.187 | -0.446 | 2.488 |
|  | *Coris batuensis* | -0.075 | -0.018 | -1.172 | 0.912 |
|  | *Coris centralis* | 0.212 | 0.069 | -0.672 | 1.469 |
|  | *Halichoeres bicolor* | -0.076 | -0.024 | -0.901 | 0.677 |
|  | *Halichoeres chierchiae* | -0.118 | -0.023 | -1.413 | 0.969 |
|  | *Halichoeres dussumieri* | -0.435 | -0.187 | -1.972 | 0.473 |
|  | *Halichoeres hortulanus* | 0.565 | 0.299 | -0.313 | 2.186 |
|  | *Halichoeres leucurus* | -0.246 | -0.087 | -1.526 | 0.665 |
|  | *Labroides dimidiatus* | 0.143 | 0.038 | -0.755 | 1.299 |
|  | *Thalassoma jansenii* | 0.173 | 0.064 | -0.711 | 1.338 |
| Mullidae | *Parupeneus macronemus* | -0.306 | -0.129 | -1.663 | 0.531 |
|  | *Upeneus oligospilus* | -0.192 | -0.053 | -1.594 | 0.805 |
| Pomacentridae | *Acanthochromis polyacanthus* | -0.329 | -0.129 | -1.780 | 0.559 |
|  | *Amblygliphidodon silolona* | -0.263 | -0.089 | -1.743 | 0.709 |
|  | *Chrysiptera talboti* | -0.253 | -0.108 | -1.461 | 0.554 |
|  | *Chrysiptera unimaculata* | -0.152 | -0.039 | -1.554 | 0.908 |
|  | *Dischistodus melanotus* | 0.100 | 0.036 | -0.732 | 1.061 |
|  | *Neopomacentrus sororius* | -0.067 | -0.014 | -1.333 | 1.077 |
|  | *Pomacentrus albicaudatus* | -0.191 | -0.098 | -1.029 | 0.426 |
|  | *Pomacentrus alleni* | 0.167 | 0.053 | -0.713 | 1.289 |
|  | *Pomacentrus amboinensis* | 0.513 | 0.276 | -0.312 | 2.035 |
|  | *Pomacentrus andamanensis* | -0.458 | -0.203 | -2.013 | 0.420 |
|  | *Pomacentrus aurifrons* | -0.099 | -0.028 | -1.344 | 0.939 |
|  | *Pomacentrus moluccensis* | -0.280 | -0.089 | -1.698 | 0.618 |
|  | *Pomacentrus philippinus* | 0.757 | 0.434 | -0.169 | 2.689 |
|  | *Pomacentrus xanthosternus* | -0.029 | -0.003 | -1.041 | 0.940 |
| Scaridae | *Cetoscarus ocellatus* | -0.143 | -0.034 | -1.494 | 1.037 |
|  | *Chlorurus capisstratoides* | -0.233 | -0.072 | -1.623 | 0.738 |
|  | *Chlorurus sordidus* | -0.154 | -0.039 | -1.486 | 0.940 |
|  | *Scarus rivulatus* | -0.107 | -0.027 | -1.335 | 0.960 |
|  | *Scarus schlegeli* | 0.201 | 0.083 | -0.638 | 1.346 |
|  | *Sparisoma chrysopterum* | 0.644 | 0.339 | -0.281 | 2.444 |
| Siganidae | *Siganus doliatus* | -0.195 | -0.050 | -1.588 | 0.839 |
|  | *Siganus puelloides* | 0.189 | 0.046 | -0.724 | 1.457 |

**Table 15:** Summary of family-level random intercepts for precision

| **Family** | **Species** | **Mean** | **Median** | **Lower CI** | **Upper CI** |
| --- | --- | --- | --- | --- | --- |
| Acanthuridae | *Acanthurus tristis* | 0.009 | 0.001 | -0.748 | 0.803 |
|  | *Ctenochaetus striatus* | 0.003 | 0.002 | -0.775 | 0.770 |
|  | *Naso elegans* | 0.000 | 0.000 | -0.741 | 0.752 |
|  | *Zebrasoma rostratum* | 0.138 | 0.040 | -0.458 | 1.019 |
| Balistidae | *Balistapus undulatus* | 0.070 | 0.018 | -0.518 | 0.767 |
|  | *Sufflamen chrysopterum* | -0.026 | -0.006 | -0.736 | 0.640 |
| Chaetodontidae | *Chaetodon trifasciatus* | 0.005 | 0.002 | -0.744 | 0.763 |
| Labridae | *Anampses elegans* | 0.070 | 0.018 | -0.587 | 0.832 |
|  | *Anampses geographicus* | 0.009 | 0.001 | -0.750 | 0.793 |
|  | *Coris batuensis* | 0.087 | 0.021 | -0.496 | 0.857 |
|  | *Coris centralis* | -0.039 | -0.008 | -0.787 | 0.630 |
|  | *Halichoeres bicolor* | 0.079 | 0.033 | -0.397 | 0.671 |
|  | *Halichoeres chierchiae* | -0.005 | -0.001 | -0.729 | 0.729 |
|  | *Halichoeres dussumieri* | 0.001 | 0.002 | -0.759 | 0.761 |
|  | *Halichoeres hortulanus* | -0.274 | -0.120 | -1.261 | 0.257 |
|  | *Halichoeres leucurus* | 0.135 | 0.042 | -0.457 | 0.990 |
|  | *Labroides dimidiatus* | -0.015 | -0.002 | -0.703 | 0.643 |
|  | *Thalassoma jansenii* | -0.040 | -0.005 | -0.741 | 0.545 |
| Mullidae | *Parupeneus macronemus* | 0.140 | 0.046 | -0.500 | 1.043 |
|  | *Upeneus oligospilus* | -0.003 | -0.002 | -0.719 | 0.726 |
| Pomacentridae | *Acanthochromis polyacanthus* | 0.003 | -0.001 | -0.742 | 0.773 |
|  | *Amblygliphidodon silolona* | 0.010 | -0.001 | -0.745 | 0.748 |
|  | *Chrysiptera talboti* | -0.038 | -0.008 | -0.747 | 0.630 |
|  | *Chrysiptera unimaculata* | -0.005 | -0.001 | -0.766 | 0.713 |
|  | *Dischistodus melanotus* | 0.010 | 0.003 | -0.598 | 0.619 |
|  | *Neopomacentrus sororius* | -0.002 | 0.000 | -0.716 | 0.736 |
|  | *Pomacentrus albicaudatus* | 0.068 | 0.028 | -0.423 | 0.637 |
|  | *Pomacentrus alleni* | -0.035 | -0.010 | -0.694 | 0.539 |
|  | *Pomacentrus amboinensis* | -0.097 | -0.028 | -0.809 | 0.434 |
|  | *Pomacentrus andamanensis* | -0.007 | 0.000 | -0.746 | 0.699 |
|  | *Pomacentrus aurifrons* | 0.002 | -0.001 | -0.745 | 0.771 |
|  | *Pomacentrus moluccensis* | 0.002 | 0.000 | -0.711 | 0.735 |
|  | *Pomacentrus philippinus* | -0.260 | -0.122 | -1.182 | 0.270 |
|  | *Pomacentrus xanthosternus* | 0.153 | 0.042 | -0.481 | 1.131 |
| Scaridae | *Cetoscarus ocellatus* | 0.010 | 0.002 | -0.731 | 0.762 |
|  | *Chlorurus capisstratoides* | 0.004 | 0.001 | -0.745 | 0.746 |
|  | *Chlorurus sordidus* | 0.005 | 0.002 | -0.768 | 0.784 |
|  | *Scarus rivulatus* | 0.001 | -0.001 | -0.747 | 0.764 |
|  | *Scarus schlegeli* | -0.052 | -0.016 | -0.695 | 0.504 |
|  | *Sparisoma chrysopterum* | -0.224 | -0.099 | -1.136 | 0.345 |
| Siganidae | *Siganus doliatus* | -0.005 | 0.000 | -0.756 | 0.742 |
|  | *Siganus puelloides* | 0.115 | 0.037 | -0.521 | 0.970 |

**Table 16:** Summary of fixed-effects for foraging time.

| **Variable** | **Mean** | **Median** | **Lower CI** | **Upper CI** | **P(>0)** |
| --- | --- | --- | --- | --- | --- |
| Intercept | 0.052 | 0.073 | -0.874 | 0.899 | 0.552 |
| Protected | -0.550 | -0.548 | -1.255 | 0.153 | 0.100 |
| Rugosity | -0.206 | -0.212 | -0.672 | 0.274 | 0.236 |
| Biomass | -0.435 | -0.437 | -0.938 | 0.084 | 0.079 |
| Predator | 0.220 | 0.224 | -0.436 | 0.871 | 0.708 |
| Group | -0.153 | -0.150 | -0.890 | 0.587 | 0.370 |
| Rugosity:Guild (Invertivore) | -0.273 | -0.278 | -0.855 | 0.300 | 0.224 |
| Biomass:Guild (Invertivore) | -0.026 | -0.025 | -0.605 | 0.542 | 0.471 |
| Protected:Guild (Invertivore) | -0.047 | -0.044 | -0.766 | 0.678 | 0.461 |

**Table 17:** Summary of zero and one hyperparameters for foraging time.

| **Variable** | **Mean** | **Median** | **Lower CI** | **Upper CI** | **P(>0)** |
| --- | --- | --- | --- | --- | --- |
| Cut One | 1.685 | 1.685 | 1.333 | 2.053 | 1.000 |
| Cut Zero | 0.927 | 0.933 | 0.247 | 1.594 | 0.985 |

**Table 18:** Summary of goodness of fit for foraging probability.

| **Type** | **Estimate** | **SE** | **Lower CI** | **Upper CI** |
| --- | --- | --- | --- | --- |
| Conditional | 0.521 | 0.098 | 0.318 | 0.706 |
| Marginal | 0.160 | 0.086 | 0.024 | 0.339 |

### ­Appendix G: Summary of analysis of protection, plot-level predictors and individual-level predictors on foraging rates.

**Table 1:** Summary of plot-level random intercepts.

| **Site** | **Plot** | **Mean** | **Median** | **Lower CI** | **Upper CI** |
| --- | --- | --- | --- | --- | --- |
| Aquarium | 1 | 0.038 | 0.011 | -0.476 | 0.597 |
|  | 5 | -0.107 | -0.055 | -0.768 | 0.446 |
| Pilot | 8 | -0.190 | -0.107 | -0.934 | 0.362 |
| Boat | 16 | -0.004 | -0.001 | -0.502 | 0.504 |
|  | 20 | -0.200 | -0.131 | -0.826 | 0.236 |
| Twins | 21 | -0.064 | -0.028 | -0.678 | 0.469 |
|  | 22 | -0.043 | -0.013 | -0.674 | 0.528 |
|  | 23 | -0.051 | -0.022 | -0.654 | 0.506 |
|  | 24 | 0.149 | 0.076 | -0.347 | 0.803 |
|  | 25 | -0.065 | -0.032 | -0.598 | 0.429 |
| Tarmugli | 26 | -0.022 | -0.003 | -0.600 | 0.499 |
| Babu's Bar | 31 | -0.127 | -0.064 | -0.761 | 0.395 |
|  | 32 | 0.201 | 0.130 | -0.265 | 0.838 |
|  | 33 | 0.114 | 0.054 | -0.366 | 0.689 |
|  | 34 | -0.082 | -0.036 | -0.666 | 0.435 |
|  | 35 | -0.096 | -0.046 | -0.717 | 0.429 |
| Deep Deception | 36 | 0.017 | 0.001 | -0.525 | 0.598 |
|  | 37 | 0.285 | 0.167 | -0.264 | 1.148 |
|  | 39 | -0.127 | -0.064 | -0.794 | 0.404 |
| Juvi | 42 | 0.076 | 0.040 | -0.376 | 0.603 |
| Oshark | 46 | 0.442 | 0.367 | -0.062 | 1.252 |
|  | 47 | -0.036 | -0.013 | -0.597 | 0.507 |
|  | 48 | 0.103 | 0.044 | -0.409 | 0.719 |
|  | 49 | -0.015 | -0.013 | -0.511 | 0.497 |
|  | 50 | -0.032 | -0.010 | -0.642 | 0.549 |
| Playground | 51 | 0.113 | 0.049 | -0.375 | 0.740 |
|  | 52 | 0.359 | 0.285 | -0.128 | 1.098 |
|  | 53 | 0.172 | 0.102 | -0.276 | 0.783 |
|  | 54 | -0.134 | -0.067 | -0.761 | 0.384 |
|  | 55 | -0.328 | -0.231 | -1.125 | 0.182 |
| Tarmugli-West | 58 | -0.359 | -0.242 | -1.231 | 0.169 |
|  | 59 | -0.006 | -0.001 | -0.588 | 0.564 |
|  | 60 | -0.266 | -0.143 | -1.140 | 0.272 |

**Table 2:** Summary of species-level random intercepts.

| **Family** | **Species** | **Mean** | **Median** | **Lower CI** | **Upper CI** |
| --- | --- | --- | --- | --- | --- |
| Acanthuridae | *Acanthurus tristis* | 0.046 | 0.012 | -0.306 | 0.493 |
|  | *Ctenochaetus striatus* | 0.009 | 0.002 | -0.367 | 0.400 |
|  | *Naso elegans* | -0.045 | -0.013 | -0.500 | 0.327 |
|  | *Zebrasoma rostratum* | 0.096 | 0.028 | -0.239 | 0.645 |
| Balistidae | *Balistapus undulatus* | 0.048 | 0.010 | -0.292 | 0.491 |
| Chaetodontidae | *Chaetodon trifasciatus* | -0.055 | -0.013 | -0.525 | 0.310 |
| Labridae | *Halichoeres bicolor* | -0.166 | -0.057 | -0.863 | 0.165 |
|  | *Halichoeres chierchiae* | -0.106 | -0.027 | -0.660 | 0.232 |
|  | *Halichoeres dussumieri* | 0.038 | 0.009 | -0.295 | 0.426 |
|  | *Halichoeres hortulanus* | -0.019 | -0.002 | -0.464 | 0.358 |
|  | *Halichoeres leucurus* | -0.013 | -0.002 | -0.423 | 0.373 |
| Mullidae | *Parupeneus macronemus* | -0.031 | -0.007 | -0.447 | 0.309 |
|  | *Upeneus oligospilus* | 0.132 | 0.038 | -0.201 | 0.758 |
| Pomacentridae | *Acanthochromis polyacanthus* | -0.021 | -0.003 | -0.477 | 0.366 |
|  | *Chrysiptera unimaculata* | -0.088 | -0.024 | -0.642 | 0.273 |
|  | *Neopomacentrus sororius* | -0.014 | -0.002 | -0.416 | 0.359 |
|  | *Pomacentrus albicaudatus* | -0.050 | -0.023 | -0.382 | 0.227 |
|  | *Pomacentrus amboinensis* | -0.013 | -0.001 | -0.413 | 0.391 |
|  | *Pomacentrus andamanensis* | 0.004 | 0.000 | -0.371 | 0.396 |
|  | *Pomacentrus moluccensis* | 0.023 | 0.003 | -0.336 | 0.424 |
|  | *Pomacentrus xanthosternus* | -0.014 | -0.002 | -0.394 | 0.338 |
| Scaridae | *Cetoscarus ocellatus* | -0.058 | -0.016 | -0.544 | 0.288 |
|  | *Chlorurus capisstratoides* | 0.089 | 0.026 | -0.233 | 0.590 |
|  | *Chlorurus sordidus* | 0.036 | 0.011 | -0.318 | 0.460 |
|  | *Scarus rivulatus* | 0.140 | 0.049 | -0.172 | 0.703 |
| Siganidae | *Siganus doliatus* | -0.054 | -0.012 | -0.510 | 0.283 |
|  | *Siganus puelloides* | -0.042 | -0.010 | -0.494 | 0.344 |

**Table 3:** Summary of fixed effects.

| **Variable** | **Mean** | **Median** | **Lower CI** | **Upper CI** | **P(>0)** |
| --- | --- | --- | --- | --- | --- |
| Intercept | -1.657 | -1.654 | -2.117 | -1.193 | 0.000 |
| Protected | 0.198 | 0.200 | -0.227 | 0.605 | 0.780 |
| Rugosity | 0.064 | 0.062 | -0.191 | 0.324 | 0.659 |
| Biomass | 0.140 | 0.141 | -0.186 | 0.458 | 0.753 |
| Predator | 0.270 | 0.267 | -0.145 | 0.690 | 0.862 |
| Group | -0.195 | -0.201 | -0.742 | 0.368 | 0.274 |
| Rugosity: Guild (Invertivore) | 0.186 | 0.191 | -0.238 | 0.592 | 0.778 |
| Biomass: Guild (Invertivore) | -0.146 | -0.135 | -0.642 | 0.339 | 0.315 |
| Protected: Guild (Invertivore) | 0.558 | 0.563 | -0.070 | 1.161 | 0.926 |
| Shape | 2.158 | 2.082 | 1.378 | 3.221 | 1 |

**Table 4:** Summary of goodness of fit.

| **Type** | **Estimate** | **SE** | **Lower CI** | **Upper CI** |
| --- | --- | --- | --- | --- |
| Conditional | 0.444 | 0.124 | 0.209 | 0.690 |
| Marginal | 0.325 | 0.109 | 0.133 | 0.557 |

### Appendix H: Marginal effects of protections and plot-level predictors on fish behaviour.

**Table 1:** Effect of protection on herbivore and invertivore foraging rates.

| **Foraging Guild** | **Median** | **90% Lower CI** | **50% Lower CI** | **50% Upper CI** | **90% Upper CI** | **P(>0)** |
| --- | --- | --- | --- | --- | --- | --- |
| Herbivore | 0.288 | -0.327 | 0.040 | 0.547 | 0.873 | 0.780 |
| Invertivore | 1.099 | 0.078 | 0.698 | 1.489 | 2.069 | 0.964 |

**Table 2:** Effect of habitat complexity (rugosity) on the probability of a fish showing foraging, vigilance or movement.

| **Behaviour** | **Foraging Guild** | **Median** | **90% Lower CI** | **50% Lower CI** | **50% Upper CI** | **90% Upper CI** | **P(>0)** |
| --- | --- | --- | --- | --- | --- | --- | --- |
| Foraging | Herbivore | 0.106 | -0.440 | -0.117 | 0.323 | 0.650 | 0.627 |
|  | Invertivore | 0.569 | -0.183 | 0.262 | 0.880 | 1.300 | 0.892 |
| Move | Herbivore | 0.049 | -0.604 | -0.214 | 0.317 | 0.698 | 0.550 |
|  | Invertivore | -0.295 | -1.115 | -0.625 | 0.058 | 0.594 | 0.287 |
| Vigilance | Herbivore | -0.087 | -0.671 | -0.311 | 0.141 | 0.508 | 0.401 |
|  | Invertivore | -0.145 | -0.962 | -0.476 | 0.222 | 0.718 | 0.394 |

**Table 3:** Effect of habitat complexity (rugosity) on the proportion of time fish were foraging, vigilant or moving.

| **Behaviour** | **Foraging Guild** | **Median** | **90% Lower CI** | **50% Lower CI** | **50% Upper CI** | **90% Upper CI** | **P(>0)** |
| --- | --- | --- | --- | --- | --- | --- | --- |
| Foraging | Herbivore | 0.184 | -0.676 | -0.157 | 0.525 | 1.062 | 0.645 |
|  | Invertivore | 0.629 | -0.772 | 0.108 | 1.151 | 1.995 | 0.788 |
| Move | Herbivore | -0.442 | -1.619 | -0.884 | -0.019 | 0.537 | 0.238 |
|  | Invertivore | -1.206 | -2.972 | -1.834 | -0.638 | 0.140 | 0.070 |
| Vigilance | Herbivore | 0.068 | -0.600 | -0.209 | 0.245 | 0.636 | 0.560 |
|  | Invertivore | 0.630 | -0.360 | 0.217 | 1.243 | 1.669 | 0.855 |

**Table 4:** Effect of habitat complexity (rugosity) on the foraging rate of herbivores and invertivores.

| **Foraging Guild** | **Median** | **90% Lower CI** | **50% Lower CI** | **50% Upper CI** | **90% Upper CI** | **P(>0)** |
| --- | --- | --- | --- | --- | --- | --- |
| Herbivore | 0.288 | -0.327 | 0.040 | 0.547 | 0.873 | 0.780 |
| Invertivore | 1.099 | 0.078 | 0.698 | 1.489 | 2.069 | 0.964 |

**Table 5:** Effect of resource availability (algal cover) on the probability of a fish showing foraging, vigilance or movement.

| **Behavour** | **Foraging Guild** | **Median** | **90% Lower CI** | **50% Lower CI** | **50% Upper CI** | **90% Upper CI** | **P(>0)** |
| --- | --- | --- | --- | --- | --- | --- | --- |
| Foraging | Herbivore | 0.344 | -0.261 | 0.102 | 0.594 | 0.951 | 0.828 |
|  | Invertivore | 0.494 | -0.345 | 0.154 | 0.848 | 1.368 | 0.836 |
| Move | Herbivore | -0.340 | -1.044 | -0.627 | -0.044 | 0.394 | 0.221 |
|  | Invertivore | -0.367 | -1.280 | -0.729 | 0.000 | 0.571 | 0.250 |
| Vigilance | Herbivore | 0.207 | -0.454 | -0.071 | 0.484 | 0.851 | 0.693 |
|  | Invertivore | 0.110 | -0.796 | -0.267 | 0.481 | 1.026 | 0.580 |

**Table 6:** Effect of resource availability (algal cover) on the proportion of time fish were foraging, vigilant or moving.

| **Behaviour** | **Foraging Guild** | **Median** | **90% Lower CI** | **50% Lower CI** | **50% Upper CI** | **90% Upper CI** | **P(>0)** |
| --- | --- | --- | --- | --- | --- | --- | --- |
| Foraging | Herbivore | 0.184 | -0.676 | -0.157 | 0.525 | 1.062 | 0.645 |
|  | Invertivore | 0.629 | -0.772 | 0.108 | 1.151 | 1.995 | 0.788 |
| Move | Herbivore | -0.442 | -1.619 | -0.884 | -0.019 | 0.537 | 0.238 |
|  | Invertivore | -1.206 | -2.972 | -1.834 | -0.638 | 0.140 | 0.070 |
| Vigilance | Herbivore | 0.068 | -0.600 | -0.209 | 0.245 | 0.636 | 0.560 |
|  | Invertivore | 0.630 | -0.360 | 0.217 | 1.243 | 1.669 | 0.855 |

**Table 7:** Effect of resource availability (algal cover) on the foraging rate of herbivores and invertivores.

| **Foraging Guild** | **Median** | **90% Lower CI** | **50% Lower CI** | **50% Upper CI** | **90% Upper CI** | **P(>0)** |
| --- | --- | --- | --- | --- | --- | --- |
| Herbivore | 0.203 | -0.268 | 0.004 | 0.396 | 0.661 | 0.753 |
| Invertivore | 0.001 | -0.770 | -0.312 | 0.309 | 0.730 | 0.501 |

**Table 8:** Effect of predator presence on the probability of a fish showing foraging, vigilance or movement.

| **Behaviour** | **Median** | **90% Lower CI** | **50% Lower CI** | **50% Upper CI** | **90% Upper CI** | **P(>0)** |
| --- | --- | --- | --- | --- | --- | --- |
| Foraging | -0.629 | -1.392 | -0.944 | -0.313 | 0.144 | 0.084 |
| Move | 0.189 | -0.686 | -0.177 | 0.556 | 1.088 | 0.635 |
| Vigilance | 0.402 | -0.443 | 0.065 | 0.726 | 1.229 | 0.785 |

**Table 9:** Effect of predator presence on the proportion of time fish were foraging, vigilant or moving.

| **Behaviour** | **Median** | **90% Lower CI** | **50% Lower CI** | **50% Upper CI** | **90% Upper CI** | **P(>0)** |
| --- | --- | --- | --- | --- | --- | --- |
| Foraging | -1.222 | -2.698 | -1.766 | -0.729 | -0.005 | 0.050 |
| Move | 0.526 | -0.893 | -0.019 | 1.103 | 2.006 | 0.744 |
| Vigilance | 0.630 | -0.659 | 0.074 | 1.091 | 1.761 | 0.769 |

**Table 10:** Effect of grouping on the probability of a fish showing foraging, vigilance or movement.

| **Behaviour** | **Median** | **90% Lower CI** | **50% Lower CI** | **50% Upper CI** | **90% Upper CI** | **P(>0)** |
| --- | --- | --- | --- | --- | --- | --- |
| Foraging | 0.923 | -0.072 | 0.538 | 1.312 | 1.889 | 0.938 |
| Move | -0.189 | -1.184 | -0.592 | 0.227 | 0.823 | 0.381 |
| Vigilance | -0.785 | -1.844 | -1.217 | -0.340 | 0.268 | 0.115 |

**Table 11:** Effect of grouping on the proportion of time fish were foraging, vigilant or moving.

| **Behaviour** | **Median** | **90% Lower CI** | **50% Lower CI** | **50% Upper CI** | **90% Upper CI** | **P(>0)** |
| --- | --- | --- | --- | --- | --- | --- |
| Foraging | 1.022 | -0.382 | 0.461 | 1.640 | 2.643 | 0.889 |
| Move | -0.269 | -1.909 | -0.905 | 0.312 | 1.272 | 0.379 |
| Vigilance | -1.087 | -2.563 | -1.718 | -0.653 | 0.056 | 0.059 |
